# Generalization of Optimal Control Saturation Pulse Design for Robust and High CEST Contrast

**DOI:** 10.1101/2025.08.21.671490

**Authors:** Clemens Stilianu, Markus Huemer, Moritz Zaiss, Rudolf Stollberger

## Abstract

**Purpose:** Optimal Control (OC) CEST pulse design for singular pulses that can be used flexibly and robustly with high saturation at different duty cycles, saturation durations and magnetic field strengths.

**Theory and Methods:** An OC framework was developed to design a single pulse shape that can be flexibly applied for arbitrary pulse train parameters and outperform typically used CEST saturation pulses shapes. The pulse design was developed primarily with a continuous wave spectrum (CW) as the optimization target, but can be easily adapted to specific scenarios. The generalized OC pulse was evaluated through simulations, phantom, and in vivo measurements on a 3 T clinical scanner. Performance was assessed in terms of contrast, robustness to field inhomogeneities, and resilience against artifacts such as Rabi oscillations and sidebands, compared to established saturation techniques.

**Results:** Investigations showed that the generalized OC pulse achieved a contrast matching CW saturation and also functioned well under field inhomogeneities. Low-pass filtering of the optimized pulse shape effectively suppressed artifacts outside the initial optimization frequency range, enabling generalization across different field strengths. Phantom experiments consistently showed higher contrast than Gaussian, Fermi, and adiabatic spin-lock pulses for various CEST agents covering most clinically relevant regimes. In vivo imaging demonstrated substantially enhanced CEST contrast for both creatine/phosphocreatine in muscle and Amide Proton Transfer (APT) in brain compared to Gaussian saturation.

**Conclusion:** The generalized OC pulse provides a robust and flexible alternative to conventional CEST saturation strategies. Its integration into the open-source Pulseq-CEST framework supports simple reproducibility and a vendor-independent implementation.

## 1 Introduction

A conventional CEST experiment uses off-resonant RF saturation pulses, followed by an imaging sequence repeated at multiple frequencies to generate a Z-spectrum. While continuous wave (CW) saturation is common in preclinical settings due to its high exchange weighting and robust spectra, it is not feasible or unreliable on clinical systems due to hardware limitations. Pulsed saturation is typically used in clinical systems, but is prone to Rabi oscillations [1] and sideband artifacts [2], which are influenced by pulse shape. Additionally, the CEST contrast is highly dependent on the chosen pulse shape and usually lower than with CW saturation [3, 4, 5, 6].

Pulsed CEST saturation can also be used to enhance the temporal resolution of CEST experiments by interleaving pulses with image acquisition. This approach allows further use of magnetization dynamics for studying exchange rates, *T*_1_ mapping, or motion correction [7, 8, 9, 10, 11, 12]. Furthermore, pulsed CEST is required for parallel transmit *B*_1_ shimming [13], [14], [15].

RF pulse shape design has been investigated by various groups in the past. Rancan et al. (2015) applied gradient-based optimization for free-form RF pulse design targeting paramagnetic CEST contrast agents with high exchange rates in preclinical settings [16]. Yoshimaru et al. (2016) employed metaheuristic multi-objective optimization to design pulsed CEST saturation based on a Fourier series for clinical applications [17]. More recently, Mohanta et al. (2024) used gradientbased optimization to design 100 ms pulses for high exchange rate and high chemical shift CEST contrast agents at 11.7 T in preclinical settings.

Off-resonant Spin Lock (SL) experiments extend conventional pulsed CEST saturation. [18]. The optimization of adiabatic pulses for SL measurements has improved robustness against field inhomogeneities, enabling high exchange weighting and stable SL saturation for imaging [19, 20, 21, 22, 23, 24, 25, 26]. However, adiabatic tipping pulses are time- and energy-intensive. . Adiabatic pulses still have limited robustness to field inhomogeneities, with performance rapidly degrading outside defined *B*_0_ and *B*_1_ tolerance ranges [27, 28, 19, 6].

A recent study shows that CEST saturation pulse trains optimized by Optimal Control (OC) offer superior contrast and robustness over Gaussian, Block Pulse Train (BPT), and Adiabatic Spin Lock (aSL) techniques for equivalent saturation times (Tsat) and Duty Cycle (DC) [6]. However, these Fully Optimized Pulse Trains (FOPTs) have a fixed *T_sat_*and while combining pulse trains can extend *T_sat_*, they lack the flexibility of single pulses (e.g., Gaussian pulses) in being applied with arbitrary DC and *T_sat_*.

This work investigates a generalization of the OC-based saturation pulse design to allow high flexibility for CEST experiments. The goal is to develop a versatile pulse that operates effectively across various duty cycles, pulse times, total saturation times, and *B*_0_ field strengths. Furthermore, we investigate whether pulses that have been designed for a specific duration can be used with different pulse durations by scaling them accordingly.

We evaluate the generalized pulse under varying conditions in both simulations and phantom experiments. Additionally, the OC performance is tested in a thigh muscle Creatine (Cr) and Phosphocreatine (PCr) contrast, as well as in a brain Amide Proton Transfer (APT) model. The resulting pulses are integrated into the open-source Pulseq CEST framework [29, 30], enabling reproducibility and easy implementation across different scanners.

## 2 Methods

### 2.1 RF Pulse design

The CEST saturation pulses are designed by minimization of the objective function *J* :

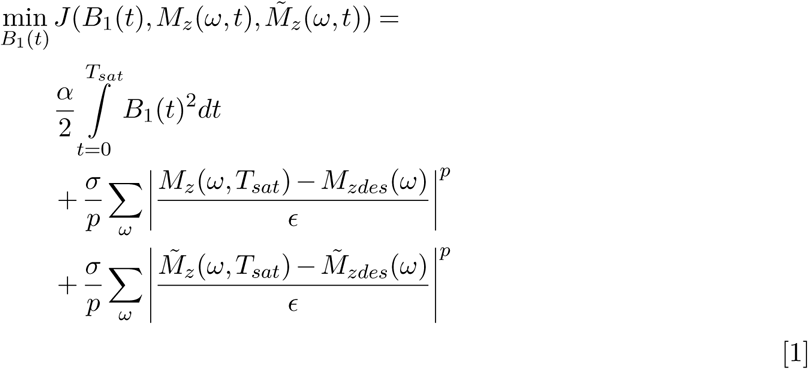

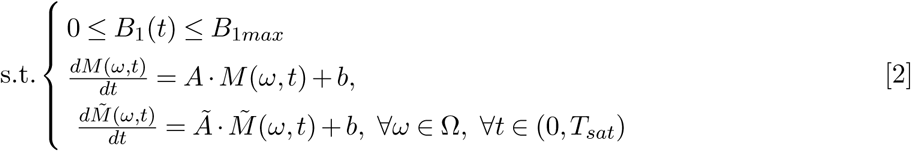

The control, *B*_1_(*t*), describes the time-dependent magnitude of the RF pulse. The first term in the cost function minimizes the energy of the RF pulse, regularized by *α >* 0. The second and third terms minimize the difference to a target continuous-wave (CW) spectrum (*M_zdes_*): first using a two-pool Bloch-McConnell simulation, where *M_z_*(*ω, T_sat_*) represents the longitudinal magnetization at frequency *ω* at the end of the saturation period *T_sat_*; and secondly using the same simulation but considering only the water pool (*M_z_*^~^*_des_* and *M*^~^*_z_*). The RF pulse is defined as a single pulse within a train of repeated pulses, separated by pauses of duration *t_p_*. The simulation is discretized in time, *t*, using a step size of Δ*t* = 10*^−^*^4^ s. Each RF pulse in the train has a duration of *t_d_*, with a duty cycle of 90 %. The gradient for optimization is computed over the entire pulse train and then averaged over the pulses to obtain a single gradient update for one pulse [31, 32]. The numerical optimization is based on a trust-region semi-smooth quasi-Newton method [33]. Further details of the optimization can be found in [6], optimization time, convergence criteria and sensitivity to start parameters are described in the supporting material.

The optimization was constrained to *T_sat_* = 1 s and a DC of 90 %. To investigate different pulse saturation times the pulse duration was set to *t_d_* = 50 and 100 ms with pulse pauses of *t_p_* = 6 and 12.5 ms. The regularization was set to *α* = 1 and the weightings were set to *σ* = 100. We iterate over *p*, starting at *p* =2 and incrementing successively as suggested by [33]. The optimization was initialized with a BPT and converged at *p* =4.

For the optimization the simulated z-spectra were discretized between 20 and −20 ppm with a step size of *δω* of 0.01 ppm, which results in *N_ω_* = 4001 off resonant points. The water pool was set to resemble in vivo-like tissue relaxation times at 3 T: *T*_1_*_w_* = 1200 ms and *T*_2_*_w_* = 80 ms. These values were chosen to be within the range of white and gray matter relaxation times at 3 T [34]. The solute pool was simulated with relaxation times *T*_1_*_s_* = 1000 ms and *T*_2_*_s_* = 160 ms and a fraction rate of *f_s_* = 0.002 at an off resonance of 3.5 ppm [35]. The exchange rate between the solute pool and the water pool was set to *k_sw_* = 200 s*^−^*^1^. These pool parameters were chosen to strike a balance between labeling efficiency and direct saturation. The *B*_0_ field was 3 T. The target spectrum was simulated with *T_sat_* = 1 s CW saturation pulse with an RF amplitude of *B*_1_ = 1 µT.

### 2.2 Comparison of a repeated generalized OC pulses against CW and the FOPT in simulation

The train of the generalized pulse is compared to the saturation of a CW pulse and the performance of the FOPT. This comparison is made for CEST pool offsets between 1 and 5 ppm, *B*_1_*_RMS_* values from 1 to 5 µT, exchange rates *k* = 250, 1000, and 3000 s*^−^*^1^, and a fraction rate *f_s_* = 0.001. The simulations are conducted with 90 % DC and *T_sat_* = 1 s in two-pool models at 3 T with a field inhomogeneity of Δ*B*_0_ = 1 ppm. The phases across the pulse trains were kept constant. The difference in performance is calculated with:

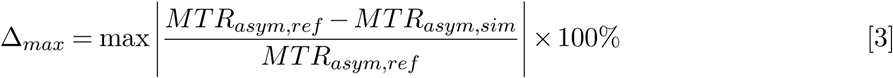

where *MTR_asym,ref_*is the contrast metric obtained with the reference simulation (e.g., CW pulse) and *MTR_asym,sim_* is that from the simulation under comparison (e.g., generalized OC pulse train).

### 2.3 Generalization of OC pulses over time and magnetic fields

To assess the influence of high-frequency Fourier components in the OC pulse on the resulting CEST spectra, the OC pulse was low-pass-filtered using an 11th-order Butterworth filter and compared to the unfiltered pulse. Simulations were conducted with a frequency sampling rate of *δω* = 0.01 ppm over a range of −30 to 30 ppm, using a pulse train of 9 pulses, a duty cycle of 90%, and *B*_1_ = 1 µT with *T*_1_*_w_* = 1200 ms, *T*_2_*_w_* = 120 ms, *T*_1_*_s_* = 1000 ms, and *T*_2_*_s_* = 160 ms.

To investigate the influence of doubling the single pulse saturation time and of the *B*_0_ field strength the filtered and unfiltered OC pulses were stretched by linear interpolation to 200 ms and simulated at 3 T and 7 T. Additionally, the filtered pulse was compressed to 75 ms and 50 ms for simulations at 3 T.

### 2.4 Pulseq implementation

The OC pulses were incorporated into the Pulseq CEST framework with a newly added function makeOCPulse(fa_sat, ’duration’, td, ’useLowPass’, true_or_false). Here the fa_sat is the flip angle over one pulse, td is the duration of a single pulse in ms and true_or_false is an option for low pass filtering the OC pulses. The function generates a Pulseq CEST native pulse object that can be used arbitrarily. The default value for td is 100 ms and for the useLowPass is false. More details about the implementation can be seen in the supporting information section 11.

### 2.5 Phantom measurements

The performance of OC saturation was compared to Gaussian, aSL and Fermi saturation in phantom measurements on a Siemens Vida 3 T clinical scanner system (Siemens Healthineers, Erlangen, Germany) with a 20 channel head coil. The Sequences were implemented in Pulseq CEST. The phantom consists of four 50 mL Falcon tubes, each containing one of the following: 0.125 g nicotinamide (NA), 0.4 g Cr monohydrate, 5 mL Ultravist (300 mg/mL iopromide (IOP)), or 3 g sucrose as a model for hydroxyl (OH) groups (example image of the phantom can be seen in the supporting information Figure S13). The pH of the phantoms was stabilized at pH 7.0 with phosphate buffer. The relaxation times were reduced to approximately *T*_1_ = 1400 ms and *T*_2_ = 120 ms with MnCl_2_. *B*_1_ and *B*_0_ maps were measured using the WASABI method [36].

Two experiments were performed:

1. For 50 ms pulses: duty cycles (DC) of 50 % (10 pulses) and 90 % (18 pulses) were applied.
2. For 100 ms pulses: duty cycles (DC) of 50 % (5 pulses) and 90 % (9 pulses) were applied.

All experiments were conducted with *B*_1_*_RMS_* values of 1, 1.5, and 2 µT. The spectra were measured at 45 frequency offsets, with higher sampling density around the CEST peaks at 1.2 ppm for OH groups, 1.7 ppm for Cr, 3.2 ppm for NA, and 4.1 ppm for IOP. *B*_0_ inhomogeneities were corrected using *B*_0_ maps from WASABI spectra, and the CEST effect was evaluated using the Magnetization Transfer Ratio asymmetry (*MTR_asym_*).

Furthermore, the nominal frequency spectra and the bandwidth of the single 100 ms Gaussian, Fermi, Block and OC pulses were calculated using a FFT and the FWHM is determined.

### 2.6 In vivo measurements

#### 2.6.1 Muscle creatine measurement

In vivo measurements were performed on a Siemens Vida 3 T clinical scanner (Siemens Healthineers, Erlangen, Germany) using an 18-channel body coil on a generally healthy volunteer. The volunteer had a prior sports-related muscle injury, which was incorporated as a variation in the scanned region. A *T*_2_-weighted transversal turbo spin echo image was acquired with the following scan parameters: *T_R_*= 3000 ms, *T_E_*= 15 ms, slice thickness = 5.5 mm, 30 slices, base resolution = 576×576, and FOV = 253×253 mm.

CEST images were generated using two different measurements comparing a state-of-the-art Gaussian saturation with an OC saturation. The Gaussian saturation was optimized to generate the highest Cr and PCr *MTR_asym_* at 3 T in calf muscle [37]. Gaussian saturation parameters: *B*_1_*_RMS_* = 1.74 µT, 90 % DC, *t_p_* = 50 ms, 11 pulses, *T_sat_* = 600 ms. OC saturation was applied with the same *B*_1_*_RMS_* and DC but with 6 pulses of 100 ms each, resulting in a saturation time of *T_sat_* = 650 ms. The readout was a centric reordered 3 D gradient echo with parameters: *T_R_* = 4 ms, *T_E_* = 2.1 ms, slice thickness = 5 mm, 5 slices, base resolution = 128 x 128, FOV = 256 x 256 x 25 mm, flip angle *α* = 8 *^◦^*. *B*_1_ and *B*_0_ maps were measured using the WASABI method. The CEST measurement times were 11:23 minutes for Gaussian and 11:28 minutes for the OC-based saturation.

The CEST spectra were generated with 101 offsets equidistant between −5 and 5 ppm. The spectra were *B*_0_ corrected using the WASABI *B*_0_ maps. The spectra were denoised using principal component analysis (PCA) using 14 principal components, the number of components was determined with the median criterion as suggested by Breitling et al. 2019 [38]. CEST images were generated using the *MTR_asym_* peak at 1.9 ppm. The *MTR_asym_*images were denoised using an NLM filter with parameters: degree of smoothing = 0.015, comparison window = 3 and search window size = 31. The images were *B*_1_ corrected using decorrelation of the CEST image with the *B*_1_ map as suggested by Papageorgakis et al. 2024 [39]. For comparing the CEST spectra in vivo, 3 regions of interest (ROIs) were analyzed: one within the muscle lesion, one in a high-SNR region with homogeneous *B*_1_, and one in a region with high *B*_1_ but low CEST SNR. Furthermore, SNR was calculated in a homogeneous ROI (124 pixels) within the vastus lateralis from the PCA denoised z-spectra.

#### 2.6.2 Brain APT measurement

Brain APT measurements were performed on the same scanner using a 20-channel head coil on a healthy volunteer (female, 32 years). The same 3D gradient echo readout sequence was used as described above. CEST saturation was based on the consensus paper protocol B2 for 3 T APT measurements [40]. Gaussian saturation parameters followed the consensus recommendations: 100 ms Gaussian pulses applied 18 times with 10 ms pauses, resulting in a 91 % DC, *B*_1_*_RMS_* = 1 µT, and *T_sat_*= 2 s to enable Lorentzian fitting. Measurement time per protocol was 9:28 min. OC saturation was applied with identical parameters but using the OC pulse shape.

CEST spectra were acquired using offsets recommended for Lorentzian fitting [41] with 10 frequency points between 3 and 4 ppm, in total 59 offsets. Spectra were *B*_0_ corrected using WASABI *B*_0_ maps. APT contrast was estimated by subtracting the background using 2-pool Lorentzian fitting for water and MT, with Lorentzian difference analysis for APT contrast quantification. The measurement spectra were denoised using PCA.

For comparison between Gaussian and OC saturation, 3 ROIs were analyzed and the mean of the spectra with maximum intensity at 3.5 ppm was used as the comparison metric.

Written informed consent was obtained from the volunteers and the study was approved by the local ethics committee.

## 3 Results

### 3.1 Comparison of a repeated generalized OC pulses against CW and the FOPT in simulation

Figure 1 presents contour plots of the simulated CEST effect for CW, the FOPT, and the proposed generalized OC saturation pulse train across *B*_1_*_RMS_* values (1–5 µT) and CEST pool offsets (1–5 ppm). Panels (a–c) display results for *k* = 250 s^-1^, (d–f) for *k* = 1000 s^-1^, and (g-i) for *k* = 3000 s^-1^. All saturation strategies show a similar well-known trend: the CEST effect increases with offset and peaks at a specific *B*_1_*_RMS_*, depending on the exchange rate. For *k* = 250 s^-1^, the maximum occurs at 2 µT, for *k* = 1000 s^-1^ at higher *B*_1_*_RMS_* around 2–4 µT, and for *k* = 3000 s^-1^ between 3–5 µT.

**Figure 1:**
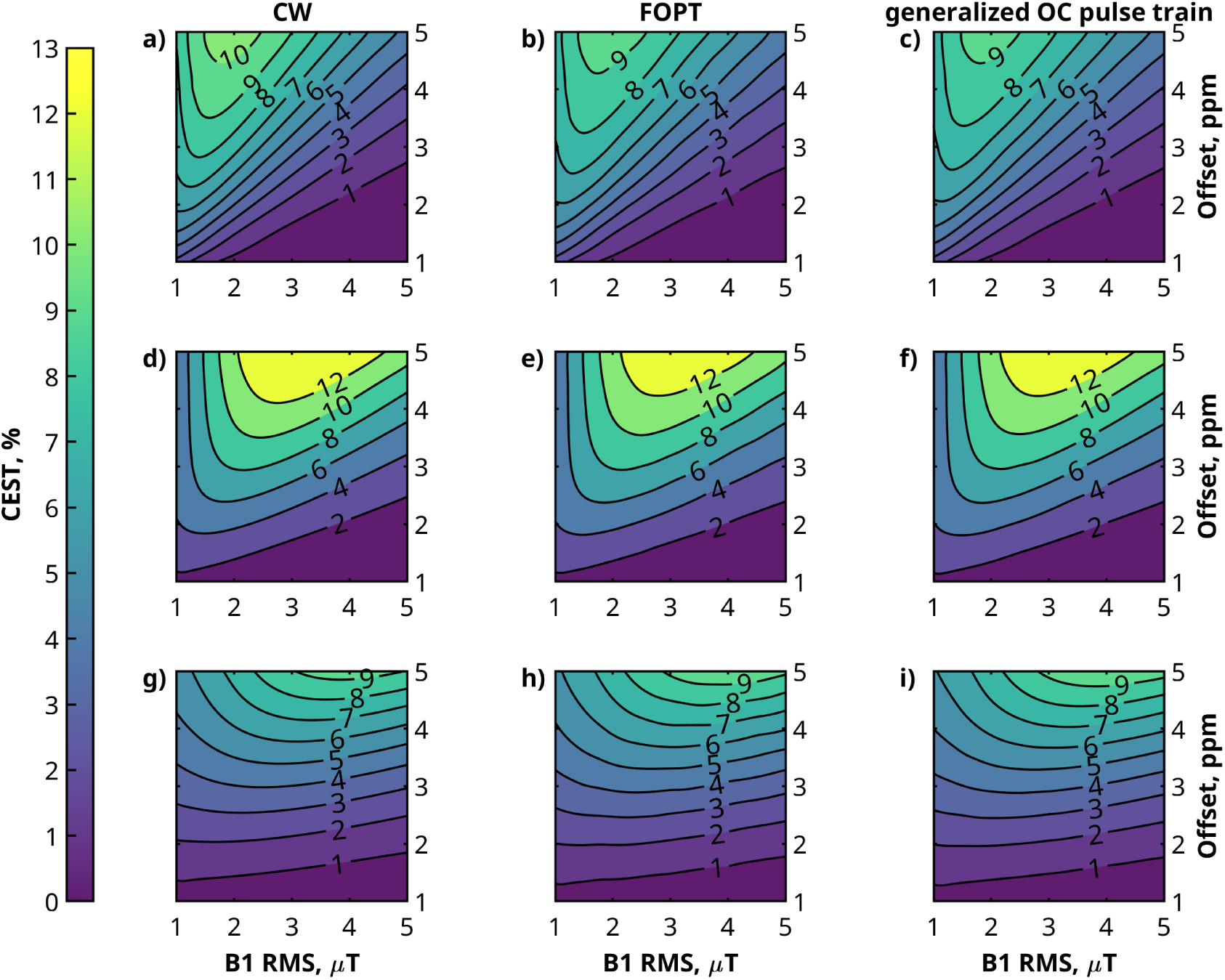
CEST effect generated by the theoretically optimal CW saturation (a, d, g) in comparison to the FOPT (b, e, h) [6] and the generalized OC pulse (c, f, i). All pulses were tested in a two-pool simulation with *T_sat_* = 1 s, for clinically relevant *B*_1_*_RMS_* values and off-resonances of the CEST pool. The OC trains have a DC of 90 %. Simulations were performed for different exchange rates: 250 *s^−^*^1^ (a, b, c), 1000 *s^−^*^1^ (d, e, f), and 3000 *s^−^*^1^ (g, h, i). For all cases a *B*_0_ inhomogeneity of 1 ppm was used.

Near the peak CEST effect, CW saturation generates a slightly larger effect than both the FOPT and generalized OC saturation strategies. For *k* = 250 s^-1^, the highest difference are 9.3 % (FOPT) and 10.4 % (generalized OC); for *k* = 1000 s^-1^, 4.1 % and 4.4 %; for *k* = 3000 s^-1^ both OC saturations are slightly higher than the CW saturation at the maximum. The *B*_0_ inhomogeneities of 1 ppm lead to no visible artifacts in the performances of the OC strategies.

### 3.2 Generalization of OC pulses over time and magnetic fields

Figure 2 (a) shows the optimized 100 ms OC pulse as obtained from the optimization process, alongside its low-pass-filtered version. The corresponding FFT of the original and filtered pulses, and the low-pass filter response, are displayed in (b). The low-pass filter suppresses frequencies above 250 Hz, preserving the overall pulse shape while smoothing oscillations, particularly at the pulse’s beginning and end.

**Figure 2:**
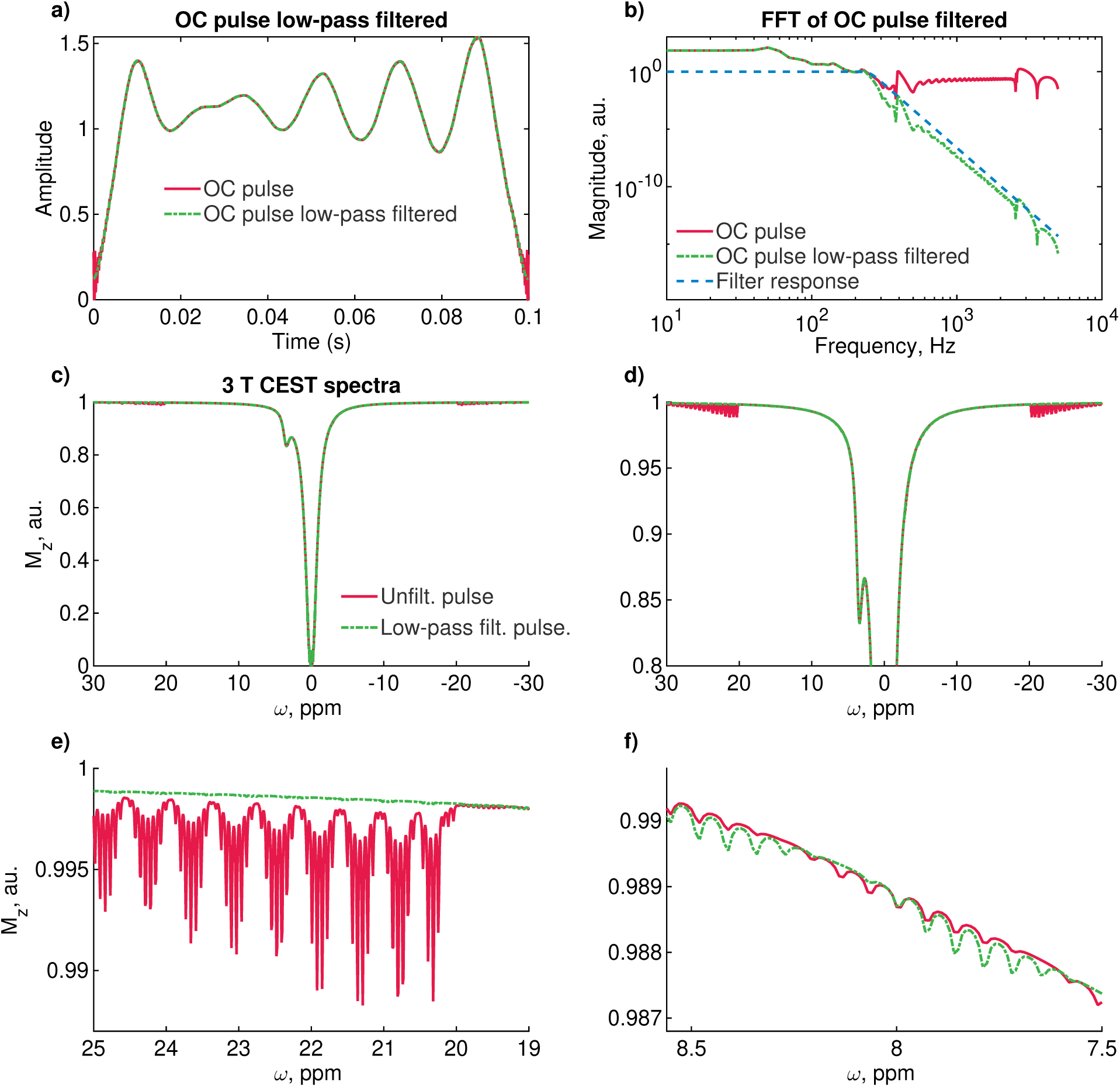
Low-pass filtering of the OC pulse: a) The original pulse from the optimization and the low-pass-filtered pulse. b) The frequency spectrum of the original and low-pass-filtered OC pulse, along with the response of the low-pass filter. c) Simulated CEST spectrum at 3 T with very fine frequency sampling of ≈ 1.3 Hz (0.01 ppm). d) The same spectrum, zoomed in to highlight sidebands at offsets > 2.6 kHz (20 ppm at 3 T). e) Further zoom into the sideband structure. f) Zoom into sidebands between 0.96 and 1.09 kHz (7.5 and 8.5 ppm at 3 T).

The impact of the low-pass-filtered pulse on the simulated CEST spectrum at 3 T is shown in Figure 2(cf). For the unfiltered OC pulse, prominent sidebands are observed at offsets outside ±2.6 kHz (±20 ppm at 3 T). The low-pass-filtered pulse generates a smoother spectrum in these regions. OC saturation also introduces sidebands inside the optimized frequency range, as exemplified in (f), with a maximum amplitude in the order of 1e-4 of the water peak. The low-pass-filtered pulse introduces wiggles with slightly higher amplitude in the CEST spectrum.

Figure 3 (a,b) shows simulated CEST spectra with 100 ms unfiltered OC pulses. (a) is at 3 T over ±3.8 kHz (±30 ppm), and (b) is at 7 T. Sidebands appear outside the optimized range (±2.6 kHz, ±20 ppm). Interpolating the 100 ms unfiltered OC pulse to 200 ms shifts the sidebands from outside ±2.6 kHz (±20 ppm) to within ±1.3 kHz (±10 ppm), as shown in (c).

**Figure 3:**
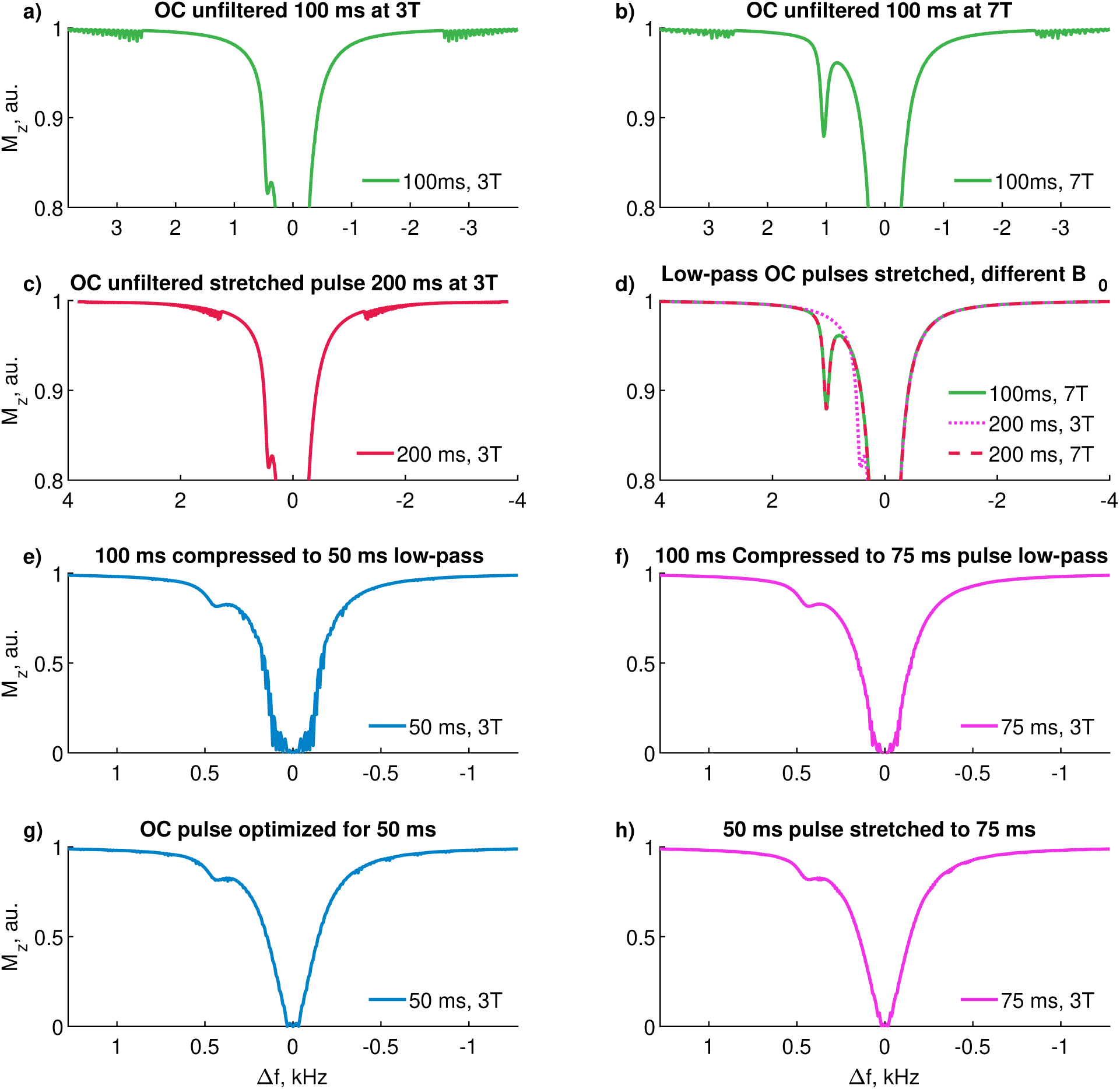
Simulations for filtered and unfiltered OC pulses stretched and compressed in time at different *B*_0_ field strengths: (a) Unfiltered 100 ms pulse simulated at 3 T with sampling of *dω* = 1.3 Hz (0.01 ppm), showing sidebands at offsets *>* 2.6 kHz (*>* 20 ppm). (b) The same pulse simulated at 7 T with sidebands at offsets *>* 3 kHz (*>* 10 ppm). (c) The 100 ms pulse interpolated in time and stretched to 200 ms, showing sidebands at offsets *>* 3 kHz (*>* 10 ppm). (d) Low-pass-filtered OC pulse simulated with 100 ms at 7 T and the filtered OC pulse stretched to 200 ms, simulated at 3 T and 7 T. (e) Simulation of the OC 100 ms pulse compressed to 75 ms, showing oscillations in the water peak. (f) Simulation of the OC 100 ms pulse compressed to 50 ms, showing stronger oscillations in the water peak.

Figure 3 (d) shows simulated CEST spectra for low-pass-filtered OC saturation pulses under various conditions: 100 ms at 7 T, 200 ms at 3 T, 200 ms at 7 T. Sidebands observed with the unfiltered pulse were not detectable in the low-pass-filtered version across all stretched pulse durations and *B*_0_ field strengths.

Figure 3 (e, f) shows CEST spectra simulated with a 100 ms pulse compressed and resampled to 75 ms and 50 ms, respectively. Figures (g, h) depict simulations of an OC pulse optimized for 50 ms and the same pulse stretched to 75 ms. Resampling the 100 ms pulse to 75 ms introduces noticeable oscillations in the water peak, which become more pronounced when further reduced to 50 ms. In contrast, the pulse optimized for 50 ms exhibits fewer oscillations, even when stretched to 75 ms. The OC pulse optimized for 50 ms and its interpolation to 75 ms exhibit minor sidebands distributed throughout the spectra.

### 3.3 Phantom measurements

Figure 4 compares the performance of four saturation pulses (Gaussian, Fermi, aSL, and OC) under varying conditions of duty cycle (50 % and 90 %) and *B*_1_*_RMS_* of (1, 1.5, and 2 µT). Across all measured regimes, the *MTR_asym_* values varied depending on the pulse and the experimental conditions. Generally, 90 % DC generated higher contrast than 50 %. In the Cr and NA phantom, the highest contrast was generated by 1.5 µT *B*_1_*_RMS_* and 90 % DC across all saturation strategies. In the IOP phantom, the generated CEST contrasts increased with higher *B*_1_*_RMS_* and DC values.

**Figure 4:**
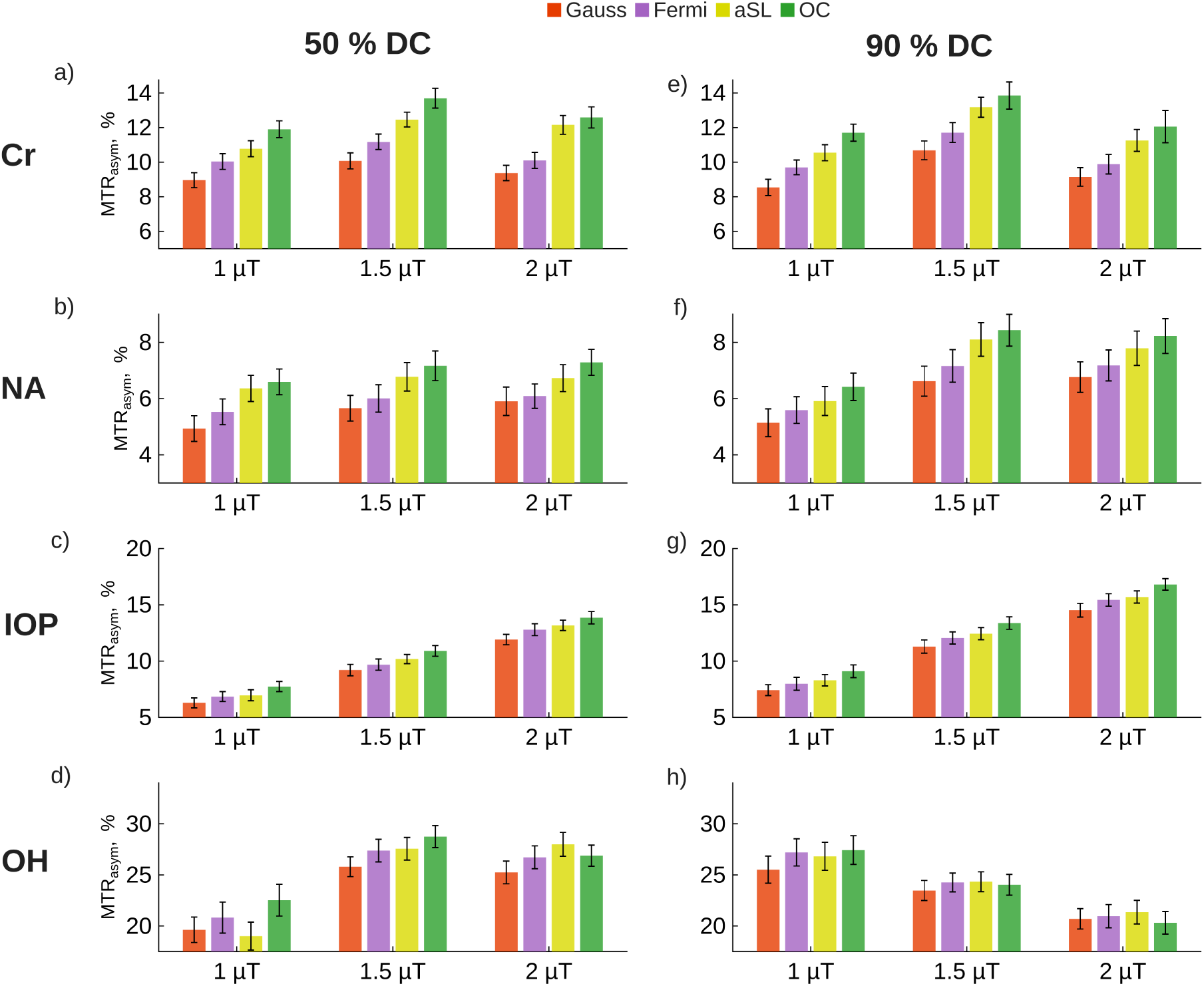
Phantom measurements comparing the performance of Gaussian, Fermi, adiabatic Spin Lock (aSL), and Optimal Control (OC) saturation pulses. All pulses were applied for 100 ms with a total saturation time of 1 s, using 50 % duty cycle (DC) in (a-d) and 90 % DC in (e-h). Saturation was performed at *B*_1_ RMS of 1, 1.5, and 2 µT. The phantom consisted of four Falcon tubes, each containing one of the following: Cr, NA, IOP, and sucrose (OH).

Differences in *MTR_asym_* values between the OC pulse and the Gaussian pulse ranged from 15.7 % to 37.1 % across all offsets and conditions. Differences between the OC and Fermi pulses ranged from 8.3 % to 24.7 %, while differences between the OC and aSL pulses were between 3.5 % and 11.2 %. The highest differences between the OC pulse and other saturation pulses were observed in the Cr measurement, with values ranging from 29.7 % to 37.1 % compared to the Gaussian pulse, 18.4 % to 24.7 % compared to the Fermi pulse, and 3.5 % to 11.0 % compared to the aSL pulse. These differences were followed by those in the NA measurement, which ranged from 21.6 % to 33.7 % for the Gaussian pulse, 14.5 % to 19.6 % for the Fermi pulse, and 3.7 % to 8.5 % for the aSL pulse. The lowest differences were observed in the IOP measurement, with ranges of 15.7 % to 22.9 % for the Gaussian pulse, 8.3 % to 14.0 % for the Fermi pulse, and 5.2 % to 11.2 % for the aSL pulse.

In Figure S1 (supporting information) the same phantom and parameters are compared as in Figure 4 but for 50 ms pulses. Across all measured regimes, the *MTR_asym_* values followed similar trends to the 100 ms pulse durations, with higher contrast generally observed at 90 % DC compared to 50 %. In the Cr and NA phantoms, higher *B*_1_*_RMS_* and DC yielded greater contrast, while in the IOP phantom, this trend was consistent across all conditions.

The *MTR_asym_* differences between the OC pulse and the Gaussian pulse ranged from 8.1 % to 29.0 %, and between the OC and Fermi pulses from 3.1 % to 17.9 %. Differences between the OC and aSL pulses ranged from 10.2 % to 33.4 %, with the highest differences observed in the Cr measurement, followed by the NA and IOP measurements. In the Cr measurement, the differences ranged from 21.2 % to 29.0 % for the Gaussian pulse, 10.1 % to 17.9 % for the Fermi pulse, and 14.3 % to 26.6 % for the aSL pulse. For the NA measurement, the differences ranged from 13.5 % to 19.2 % for the Gaussian pulse, 5.5 % to 13.6 % for the Fermi pulse, and 10.2 % to 26.9 % for the aSL pulse. The IOP measurement showed the lowest differences, ranging from 8.1 % to 25.0 % for the Gaussian pulse, 3.1 % to 16.0 % for the Fermi pulse, and 11.2 % to 33.4 % for the aSL pulse.

The frequency spectra of single 100 ms Gaussian, Fermi, Block and OC pulses, show bandwidths of 20 Hz, 17 Hz, 12 Hz, and 13 Hz respectively (see supporting material (Figure S12, supporting information))

### 3.4 In vivo measurement

#### 3.4.1 Muscle creatine measurement

Figure 5 shows a transverse section through the right thigh. (a) displays a *T*_2_-weighted image highlighting the injury in the rectus femoris, with increased *T*_2_ signal intensity (red arrow). (b) shows a *B*_1_ map, with *B*_1_ values ranging from 0.5 to 1.4 µT. (c, d) show Cr and PCr *MTR_asym_* CEST images acquired with the Gaussian saturation protocol and the generalized OC saturation. Elevated Cr and PCr contrast is visible in the rectus femoris and parts of the vastus intermedius in both Gaussian and OC images. The OC images exhibit higher contrast and more uniform contrast distribution, with fewer hyper-intense regions and higher contrast to noise compared to the Gaussian images.

**Figure 5:**
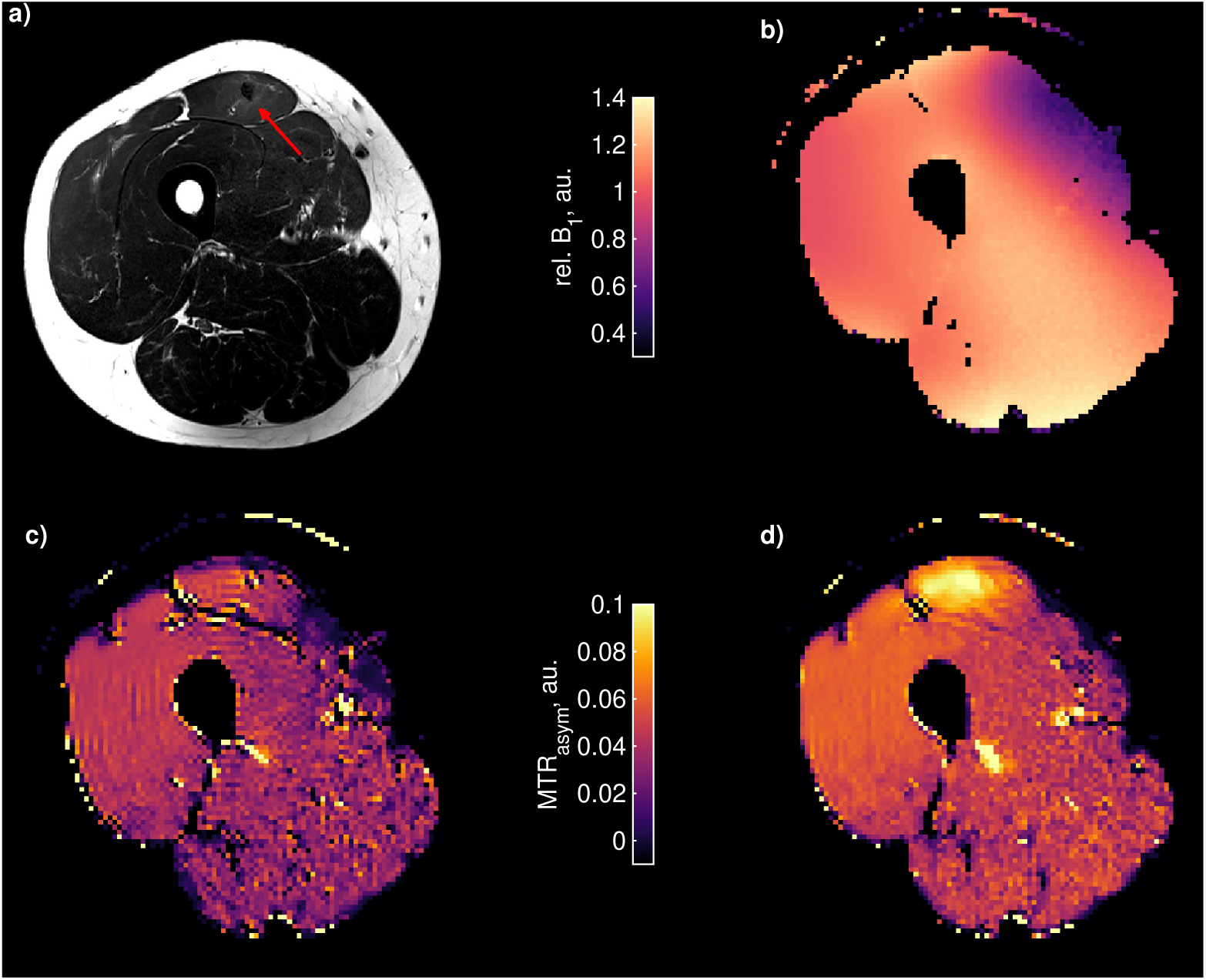
Comparison of Gaussian versus OC saturation for creatine CEST imaging in thigh muscle with an injury in the rectus femoris: (a) *T*_2_-weighted image with increased values see red arrow, (b) relative *B*_1_ map, (c) Gaussian saturation protocol with 11 pulses of 50 ms each and a duty cycle (DC) of 90 %, with a *B*_1,RMS_ of 1.74 µT, and (d) generalized OC saturation with 6 pulses of 100 ms each, also with a 90 % duty cycle, and a *B*_1,RMS_ of 1.74 µT. The measurement spectra were denoised using PCA, while the creatine CEST maps were denoised using NLM. The creatine CEST maps were *B*_1_ corrected using decorrelation of the CEST and *B*_1_ map as proposed by [39]. Some hyperintensities are apparent in the CEST images, these are flow-related artifacts due to blood vessels.

Figure 6 shows the mean value of *MTR_asym_*from ROIs in three regions of the CEST images generated with (a) the Gaussian saturation and (b) the OC saturation. (c, e, g) display the full CEST spectra in the rectus femoris lesion area ROI, while (d, f, h) highlight the *MTR_asym_* peaks.

**Figure 6:**
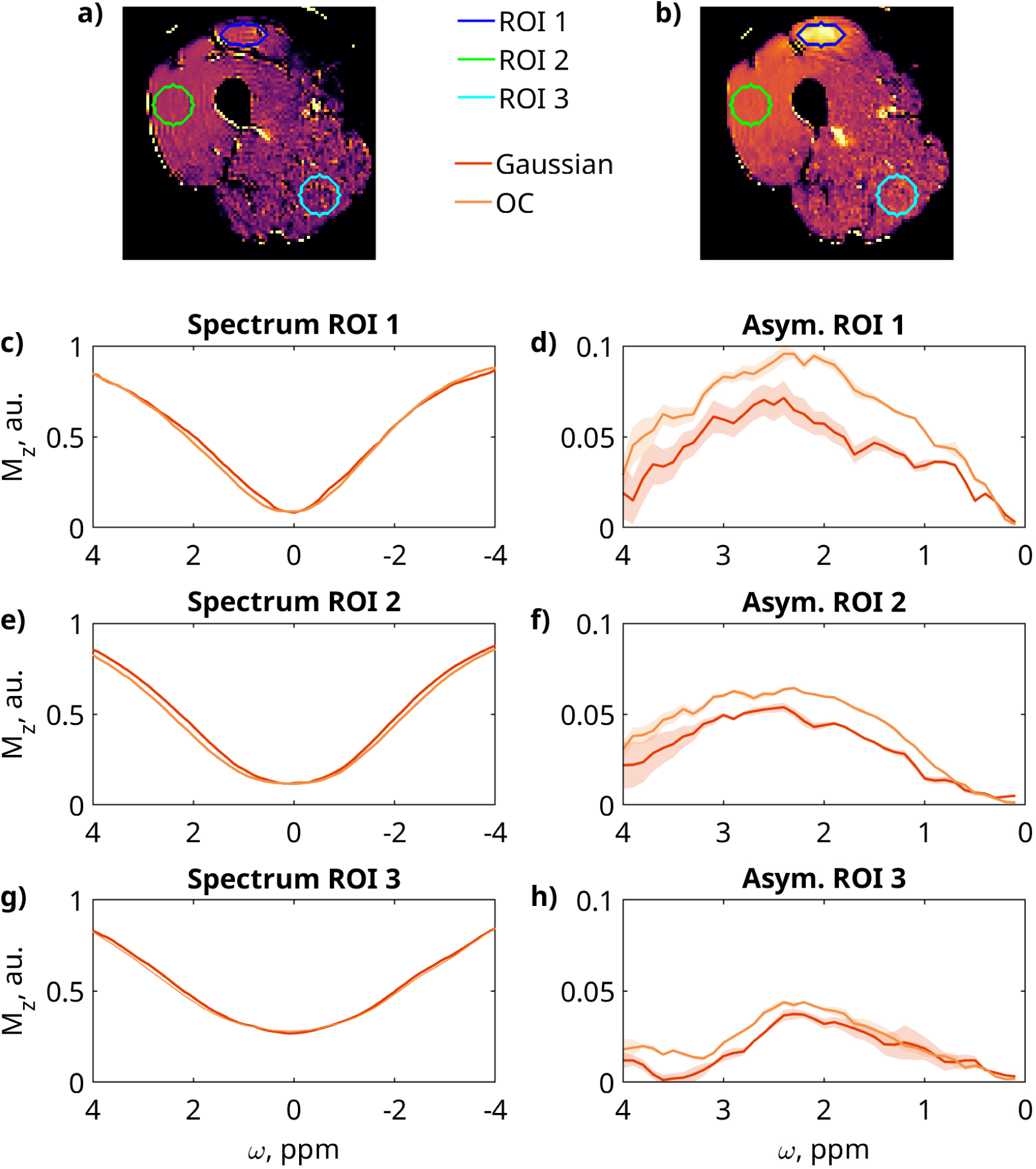
Comparing in vivo CEST spectra in different ROIs: a) Creatine CEST image generated with Gaussian saturation with ROIs. b) Creatine CEST image generated with OC saturation with ROIs. CEST spectra and *MTR_asym_*with standard deviation in the lesion region of the rectus femoris, (c,d) in the vastus lateralis and (g,h) in a noisy, low *B*_1_ region.

The mean *MTR_asym_* values and standard deviations for each ROI are as follows: For ROI 1, OC saturation was 0.090 ± 0.003 and Gaussian saturation was 0.053 ± 0.006, resulting in 70.4 % higher mean and 50.5 % lower standard deviation for OC saturation. For ROI 2, OC saturation was 0.0592 ± 0.0006 and Gaussian saturation was 0.0448 ± 0.0011, leading to 32.1 % higher mean and 47.3 % lower standard deviation for OC saturation. For ROI 3, OC saturation was 0.0391 ± 0.0022 and Gaussian saturation was 0.0330 ± 0.0028, yielding 18.6 % higher mean and 20.3 % lower standard deviation for OC saturation. Across all ROIs, the OC saturation consistently yields higher *MTR_asym_* peak values compared to the Gaussian saturation. SNR calculation in the vastus lateralis yielded values of 4.16 for Gaussian saturation and 6.68 for OC saturation. The ROI used for the calculation can be seen in Figure S21 of the supporting information.

#### 3.4.2 Brain APT measurement

Figure 7 shows the results of the APT measurement in brain using Lorentzian fitting. (a) displays the measurement points in ROI 1 with the fitted Lorentzian background for Gaussian and OC saturation, (b) shows the Lorentzian difference magnetization transfer ratio (*MTR_LD_*) in ROI 1. (c) depicts the mean values of the *MTR_LD_* in ROI 1, 2, and 3 which are shown in (e). The *B*_1_ and *B*_0_ distributions (d,f) in the ROIs were: ROI 1: (1.05 ± 0.03) µT and (0.01 ± 0.02) ppm, ROI 2: (1.00±0.03) µT and (0.13±0.05) ppm, ROI 3: (0.95±0.03) µT and (−0.04±0.04) ppm, respectively. The OC APT contrast was 33.1 %, 28.3 % and 33.6 % higher in ROIs 1, 2 and 3, respectively. The parameter maps for water, MT and APT can be seen in supporting information Figure S20.

**Figure 7:**
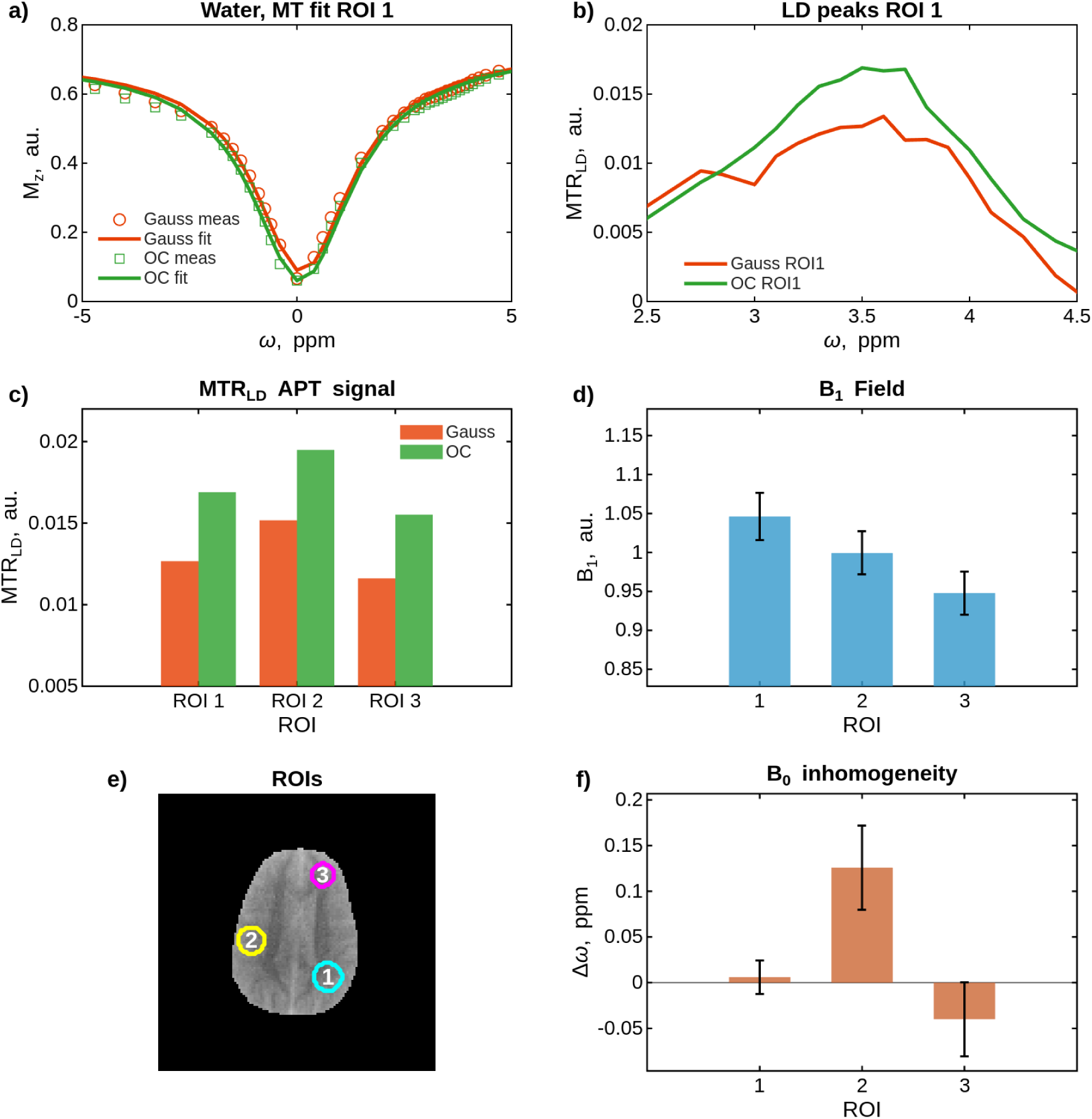
In vivo brain APT-CEST imaging at 3T with 90 % DC, *T_sat_* = 2s, *B*_1_*_RMS_* = 1 *µ*T for *t_p_* = 100 ms Gaussian and OC saturation pulses. (a) Z-spectra fitted with two-pool Lorentzian model (water and MT pools), (b) Lorentzian difference spectra after background subtraction, (c) MTR_LD_ values at 3.5 ppm across three ROIs, (d) *B*_1_ field distribution and standard deviation within ROIs, (e) *T*_1_-weighted image showing ROI locations, (f) *B*_0_ distribution within ROIs.

## 4 Discussion

In previous work, we introduced an OC saturation pulse train for enhanced and robust contrast in CEST MRI on clinical scanners [6]. This pulse train was optimized as a whole for a specific saturation duration with a fixed duty cycle. Changing these parameters requires a new optimization, which limits the flexibility for experimental use.

Here, we propose a generalized OC pulse that is optimized as a single unit and can be repeated to form a pulse train, enabling adaptation to arbitrary DC, *T_sat_*, *t_d_ >* 50 ms, *B*_1_ levels, offsets, exchange rates, and all frequency ranges, making it easy to use for all *B*_0_ fields. We demonstrate its excellent performance across all investigated regimes by comparing it to CW saturation in simulations, to state-of-the-art saturation strategies in phantom experiments across various CEST agents, and to Gaussian saturation optimized for Cr and PCr CEST contrast in an in vivo example. The proposed pulses are integrated into the Pulseq-CEST framework, providing an easily accessible alternative to conventional saturation pulses.

The optimization of RF saturation pulses for CEST MRI has been explored also by other groups. Yoshimaru et al. [17] applied a genetic algorithm using smooth basis functions to optimize pulsed CEST saturation, ensuring a controlled spectral profile but inherently limiting the achievable contrast due to constraints in the solution space. More recently, Mohanta et al. introduced a 100 ms free-form optimized pulse for 11.7 T and 99 % DC, optimizing for a single spectral offset but without explicitly addressing spectral properties for imaging. In this work, we introduce a generalized OC pulse that optimizes across the full spectral range, ensuring high and robust contrast while maintaining spectral stability and mitigating sideband artifacts and Rabi oscillations across various physical and measurement parameters.

### 4.1 Comparison to CW and FOPT in simulation

Figure 1 demonstrate that the proposed OC pulse generates CEST contrast comparable to the published pulse across all tested conditions. This includes offsets between 1-5 ppm, *B*_1_*_RMS_* values from 1-5 µT, and exchange rates of 250, 1000, and 3000 s*^−^*^1^, covering most clinically relevant CEST agents. Near the peak CEST effect, CW saturation produces about 10 % higher contrast than OC strategies at *k* = 250 s*^−^*^1^ and about 4 % higher at *k* = 1000 s*^−^*^1^. These deviations are mainly due to the shorter effective saturation time for a duty cycle of 90 % and the same *T_sat_*. However, at *k* = 3000 s*^−^*^1^, both OC and FOPT pulses slightly exceed CW saturation due to a slightly lower spillover. Moreover, the simulated *B*_0_ field inhomogeneity of 1 ppm demonstrates the robustness of the OC saturation. The *B*_0_ robustness is indirectly enforced in the simulation by maintaining a constant phase offset between pulses. For a more detailed explanation, see [6]. Further simulations with different tissue parameters of OC pulses in comparison to CW can be seen in [6].

### 4.2 Generalization over frequency, ***B*_0_** and ***t_d_***

To ensure a stable spectrum generated by OC pulses, the simulated spectrum is optimized to resemble a CW spectrum with 4001 equidistant frequency components. The high-frequency sampling (0.01 ppm) is necessary to show sideband artifacts [2] and consequently suppress sidebands and Rabi oscillations. However, this stability is only ensured within a limited frequency range, i.e., ±2.6 kHz (20 ppm at 3 T). Beyond this range, saturation introduces small artifacts (Figure 2 (c-d)). Theoretically, wider ranges are possible, but ±20 ppm cover more than the relevant frequency range and avoid unnecessarily inflating the optimization problem.

The artifacts outside the optimized spectral range could be mitigated by restricting high frequencies in the pulse via low-pass filtering. Suppressing frequencies above 250 Hz in the OC pulse resulted in a smoother spectrum outside the optimized frequency range (*<* −2.6 kHz and *>* 2.6 kHz). Yet, OC saturation introduces slight oscillations in the spectrum (Figure 2 (f)), which slightly increase with low-pass filtering. These oscillations, on the order of 10*^−^*^4^ of the water signal, are negligible in measurements. Thus, low-pass filtering improves the pulse behavior beyond the optimized frequency range.

The relevant frequency range for CEST experiments depends on the *B*_0_ field strength. When the unfiltered pulse is simulated at 7 T at ±20 ppm (or ±6.0 kHz), wiggles appear at approximately 8.6 ppm (2.6 kHz) (see Figure 3(b)). A similar effect is observed when the OC pulse is stretched to 200 ms at 3 T (Figure 3(c)), where the wiggles appear at 1.3 kHz (10 ppm). Figure 3(d) shows that low-pass filtering eliminates these artifacts in all cases, including the combination of 200 ms at 7 T. Therefore, low-pass filtering generalizes OC saturation across different *B*_0_ field strengths and for pulse durations *t_d_ >* 100 ms.

Compressing a 100 ms pulse to *t_d_ <* 100 ms is less straightforward. Figure 3(e,f) shows simulations for 50 ms and 75 ms pulse durations. At 75 ms, artifacts begin to appear within ±160 Hz (1.25 ppm) and become more pronounced at 50 ms, where they can be observed in the range of ±320 Hz (2.5 ppm) at 3 T.

To minimize these artifacts for short OC pulses, a dedicated pulse was optimized for 50 ms. Its performance is shown in Figure 3(g), along with its stretched version at 75 ms. The 50 ms-optimized pulse exhibits significantly fewer artifacts than the compressed 100 ms pulse. Furthermore, stretching this optimized 50 ms pulse to 75 ms reduces wiggles near the water peak compared to simply compressing a 100 ms pulse to 75 ms. However, stretching short pulses too far, e.g., from 50 ms to 100 ms, degrades performance compared to pulses optimized directly for 100 ms.

All shaped CEST pulses generally introduce some amount of Rabi oscillations and sidebands into the spectrum. Rabi-oscillations depend on the adiabatic properties of the pulse shape, slowly varying RF pulses show less oscillations. The origin of sidebands is more complex, they emerge when some pulse shape dependent resonance condition is fulfilled at some off resonant frequencies [2].

### 4.3 Phantom measurements

To assess the performance of generalized OC pulses in phantom CEST measurements, we selected four agents covering a broad range of chemical shifts and exchange rates (1.2-4.1 ppm). Fast-exchanging OH groups at 1.2 ppm and slow-exchanging Cr at 1.7 ppm, which is close to water, are more susceptible to spillover, artifacts, and Rabi oscillations. In contrast, NA (3.2 ppm) and IOP (4.1 ppm) are farther from water, reducing spillover effects.

Across all investigated *B*_1_*_RMS_* levels, offsets, and exchange rates, OC pulses achieved the highest saturation efficiency for 50 ms and 100 ms durations with duty cycles of 50 % and 90 %. The only exception were OH measurements, where aSL led to higher saturation at higher *B*_1_*_RMS_* levels due to its lower direct water saturation. However, OC saturation produced higher overall contrast for OH exchange, even at lower *B*_1_*_RMS_* levels.

For Cr, NA, and IOP, 100 ms pulses outperformed 50 ms, while OH groups responded better to 50 ms. aSL was the exception, performing markedly worse at 50 ms across all regimes. BPT saturation data was excluded due to their sensitivity to *B*_0_ inhomogeneities (up to 0.1 ppm). Additional simulations for the performance of the different pulses over varying *B*_1_*_RMS_* values can be seen in Figure S16 in the supporting information.

### 4.4 In vivo measurements

#### 4.4.1 Muscle creatine measurement

Muscle was chosen for in vivo experiments due to its pronounced Cr and PCr CEST peaks at 3 T. Measurements were performed in the thigh muscle of a generally healthy volunteer (Male, 28 years), with a prior sports-related muscle injury. The experiment compared the generalized OC pulse to a previously optimized Gaussian saturation for Cr and PCr imaging [37]. The OC pulse train was configured to replicate the Gaussian train as closely as possible. To meet the 90 % duty cycle, six OC pulses with longer pauses were used, making the pulse train by 50 ms longer compared to the Gaussian train.

OC saturation provided higher and more homogeneous CEST contrast at 1.9 ppm, with less noise and much higher saturation in low *B*_1_ regions (≈40 % lower) in comparison to Gaussian saturation. In ROI 3, where relative *B*_1_ was approximately 30 % higher, OC saturation yielded a 19 % increase in contrast.

Due to coil inhomogeneities and artifacts at tissue boundaries, it is difficult to specify the SNR for different saturation pulses exactly. However, Figure 6 shows that the mean *MTR_asym_* value for the OC pulse is higher at all *B*_1_ levels in the three ROIs, and that the associated variance for the OC pulse is lower. The contrast in the vastus lateralis which has the fewest artifacts and homogeneous *B*_1_ (rel. *B*_1_ ≈ 1.0), the SNR is 4.16 for Gaussian saturation and 6.68 for OC saturation, representing a 61 % improvement with OC saturation.

Interestingly, the contrast was markedly increased in the region with the muscle lesion (ROI1). To the best of our knowledge, this is the first detection of elevated Cr contrast in muscle tissue adjacent to a rupture detected using CEST imaging.

One possible explanation is that Cr leakage from ruptured muscle cells increases its extracellular concentration, where it has greater exposure to water, leading to higher contrast. Kogan et al. demonstrated that Cr CEST imaging, in contrast to Magnetic Resonance Spectroscopy, is a suitable method for detecting free Cr in muscle [42, 43], supporting this hypothesis. Another possibility is that local Cr metabolism is altered in response to muscle injury.

#### 4.4.2 Brain APT measurement

The APT measurement was chosen to contrast the muscle measurement, as the brain provides a controlled environment with less field inhomogeneities where APT contrast is well-established and Lorentzian fitting can reliably separate individual CEST pools. Unlike the muscle measurement where only *MTR_asym_* analysis was feasible due to the high *B*_1_*_RMS_* (1.74 µT) and low *T*_2_ conditions that produce excessively broad spectra unsuitable for Lorentzian fitting, the brain measurement conditions (*B*_1_*_RMS_* = 1 µT, broad frequency sampling) enabled robust Lorentzian fitting of water and MT for isolated APT quantification. Lorentzian difference analysis was necessary instead of *MTR_asym_* to avoid NOE contamination, which would mask the OC label efficiency since both APT and NOE pools are affected.

The OC saturation demonstrated consistently ≈ 30% higher APT contrast compared to the consensus Gaussian protocol across all ROIs, which showed minimal *B*_0_ and *B*_1_ field variations. This improvement exceeds NA phantom measurements, confirming that OC saturation benefits can be extended to physiologically relevant conditions.

### 4.5 Single pulse optimization trade-off

Transitioning from a pulse train where each pulse is different to individually optimized single pulses significantly increases the versatility of the approach. A key advantage of this design is that each pulse prepares the magnetization completely independent of the previous pulse. This independence enables interleaved measurements, where the spectrum can be acquired after each pulse. This technique allows for studying magnetization dynamics in greater detail. Potential applications include motion correction, exchange rate estimation, and integrating *T*_1_ mapping within the same experiment. This effect is demonstrated in Figure S2 (supporting information), where simulations show how a smooth spectrum forms after each saturation pulse.

Optimizing a single 100 ms pulse introduced slight artifacts in the water spectrum between −1 and 1 ppm at 3 T, which could not be fully suppressed within a single-pulse design. These features are only apparent at high spectral resolution (Δ*ω* = 0.01 ppm) and are also commonly observed in other saturation strategies [2]. An alternative cost functional allowed for solutions where either two alternating pulses or a pulse train terminating with a second, distinct pulse effectively mitigated these wiggles, yielding a spectrum comparable clean to the FOPT [44]. These strategies, along with the modified cost functional, are detailed in the supporting information (Figure S3).

Minor residual oscillations in the single-pulse design typically do not impact the measured CEST effects. RF spoiling after each pulse further suppresses these artifacts, as shown in simulations and phantom data (see supporting information Figures S4, S5). Other RF pulses, for example Gaussian and Fermi, also introduce spectral artifacts (see supporting information Figure S6). Given the complexity of the two-pulse approach, the simpler single-pulse design remains preferable despite minor imperfections. Nonetheless, we propose an alternative method to further reduce spectral artifacts.

### 4.6 Choice of the target spectrum

Many seminal publications are based on CW saturation, or refer to it as an objective to be achieved [16, 45, 46]. Technical adjustments to specific whole-body scanners that adapt the RF system for CW saturation also emphasize this point [40]. For multi-pool systems at 7 T and low exchange rates, we (M. Z.) found that Gaussian saturation pulses of a suitable amplitude can achieve greater saturation than CW saturation. An similar effect was described previously by van Zijl et al. [47], who found that higher fields reduced NOE contrast due to increased selectivity. Simulations show that this effect occurs within a specific *B*_1_*_RMS_* exchange rate window (see Supplementary Information Figures S7, S8 and S9 of the supporting information). For this multipool system, an OC pulse was subsequently optimized Figure S10 and S11 to demonstrate the flexibility of the pulse design.

In single-pool optimizations, OC phase variation was allowed but the optimal solution consistently yielded zero phase. In contrast, multi-pool (5 pool) OC optimization produced pulses with a phase sweep (see Figure S10, supporting information), outperforming CW saturation at low exchange rates (see Figure S11). The target spectrum was based on short Gaussian pulses, but the optimal target in the multi-pool case remains unclear. Future work should explore improved targets and the role of frequency sweeps in enhancing CEST contrast in multi-pool peaks.

This work is a methodological development for optimizing the design of saturation pulses and with the change of the target and more or less degrees of freedom for the RF pulse it is a flexible framework for various challenges.

## 5 Conclusion

With this work, we demonstrate that previously designed OC saturation trains can be largely generalized using a single unit that can be flexibly applied and demonstrates strong performance across all investigated duty cycles, exchange rates, offsets, *B*_1_*_RMS_* values, and *B*_0_ field strengths. The performance of the OC pulses has been validated through simulations against CW saturation and phantom measurements against state-of-the-art saturation strategies. Furthermore, in vivo measurements were employed to demonstrate that OC saturation benefits extend to physiological conditions. The proposed pulses are integrated into the open source Pulseq-CEST framework, providing an easily accessible alternative to conventional saturation pulses.

## 6 Data Availability Statement

All mentioned OC pulses will be provided over a git repo after publishing the paper, including example simulations and suggested usage. The function integrating the OC saturation will be provided in the Pulseq CEST library.

## Acknowledgments

This research was funded in whole, or in part, by the Austrian Science Fund (FWF) I4870. We also acknowledge funding by the German Research Foundation (DFG, Project 442377885).

We want to thank Moritz Fabian for providing code for postprocessing of in vivo measurements, which served as inspiration for our implementation.

## 7 Conflict of Interest Statement

The authors declare no competing interests.

## Funding

This research was funded in whole, or in part, by the Austrian Science Fund (FWF) I4870 and by the Deutsche Forschungsgemeinschaft (DFG, German Research Foundation) Project 442377885. For the purpose of open access, the author has applied a CC BY public copyright license to any Author Accepted Manuscript version arising from this submission.

## 9 Supporting Information

## 10 Phantom measurements for 50 ms pulses

**Figure 8:**
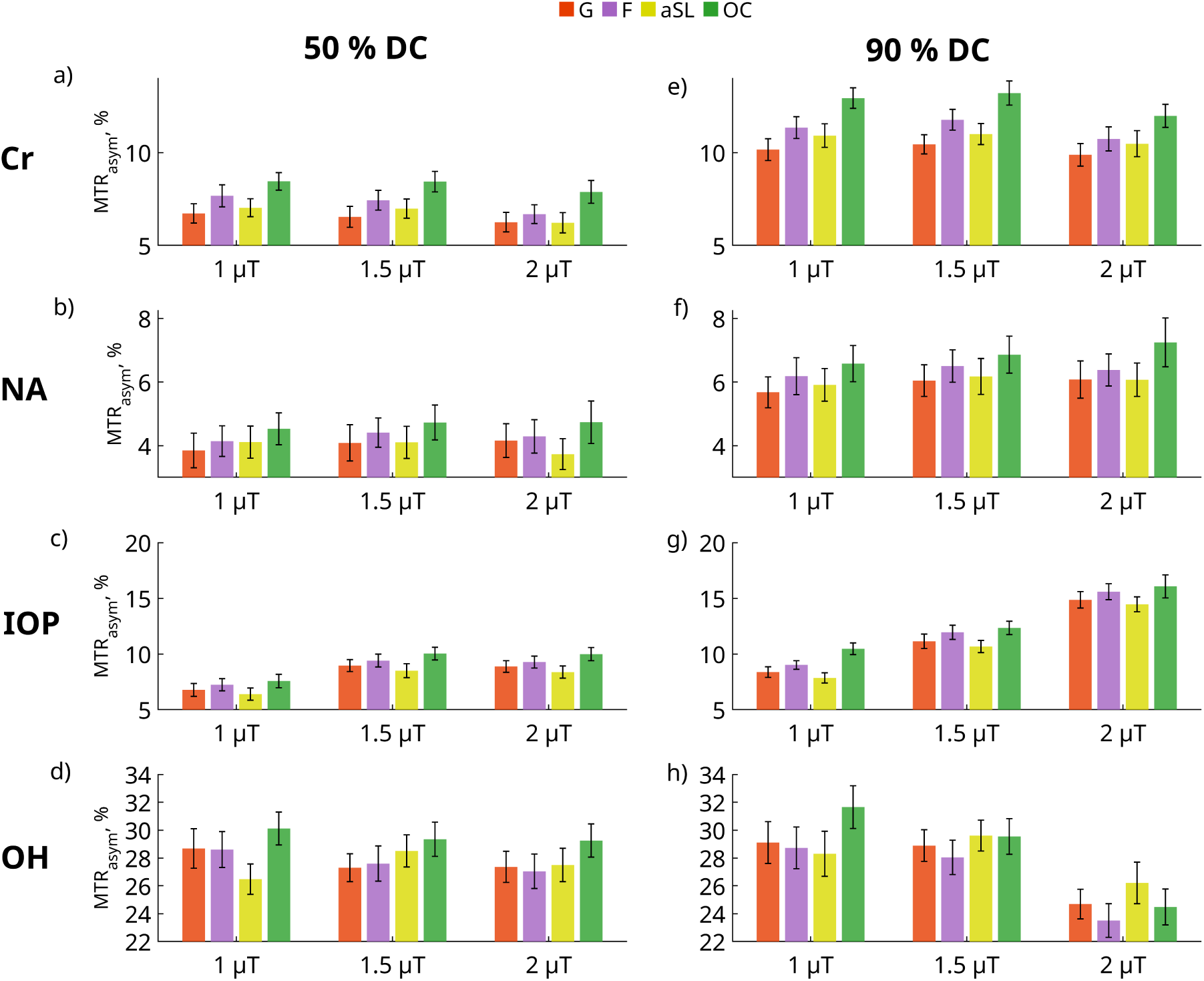
Phantom measurements comparing the performance of Gaussian, Fermi, adiabatic Spin Lock (aSL), and Optimal Control (OC) saturation pulses. All pulses were applied for 50 ms with a total saturation time of 1 s, using 50 % duty cycle (DC) in (a-d) and 90 % DC in (e-h). Saturation was performed at *B*_1_ RMS of 1, 1.5, and 2 µT. The phantom consisted of four Falcon tubes, each containing one of the following: Cr, NA, IOP, and sucrose (OH).

## 11 Interleaved measurements, magnetization after each saturation pulse in a pulse train

Figure 9 depicts the magnetization after each pulse in a pulse train of 20 OC pulses, each 100 ms long with a 90% DC. With the exception of the *T*_2_-dependent Rabi oscillations in the water peak (simulated with *T*_2_ = 60 ms), the magnetization remains smooth. This is crucial for interleaved measurements where an artifact-free magnetization and independence from the preceding pulse are required, as any measurement might alter the previously prepared magnetization and compromise the outcome.

**Figure 9:**
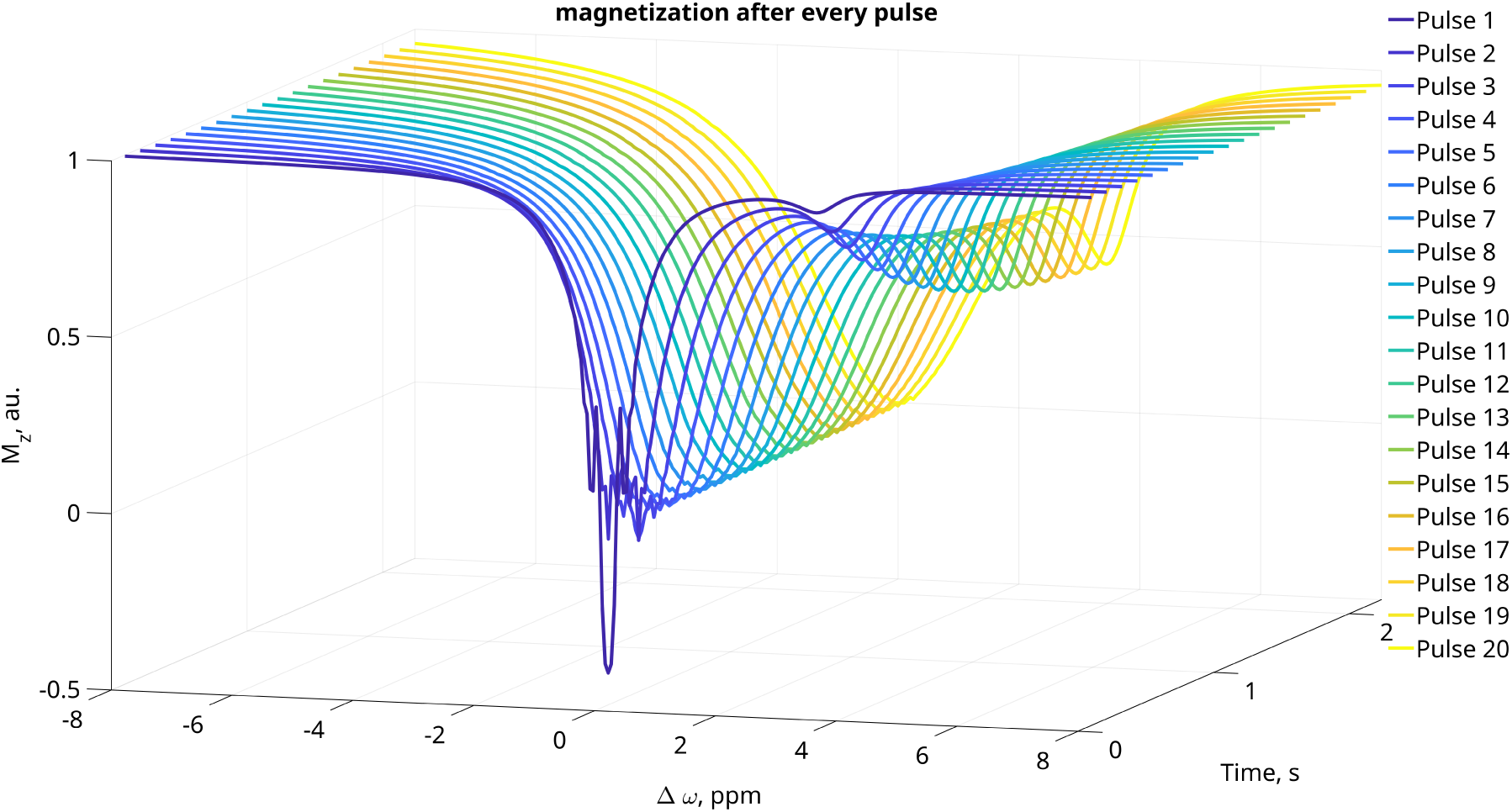
Waterfall plot for the magnetization over time. The Magnetization is measured after every pulse.

## 12 Alternative formulation of cost functional

Since optimizing 1000 free parameters for *B*_1_(*t*) over 100 ms did not yield a spectrum comparable to that obtained with 9000 free parameters over a 1 s pulse train, we introduced a modified cost functional:

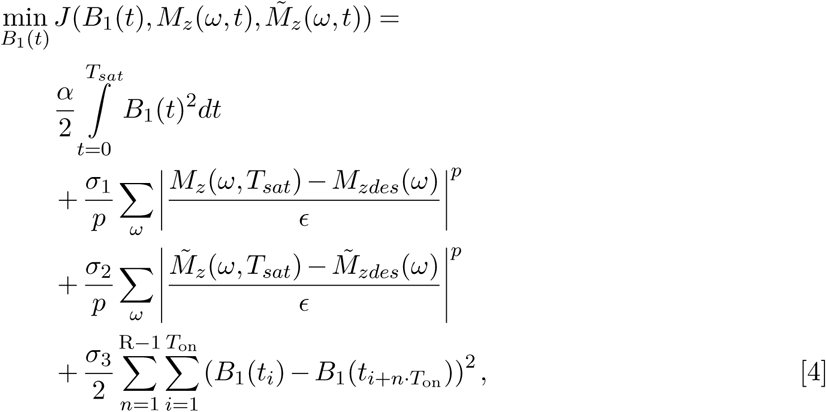

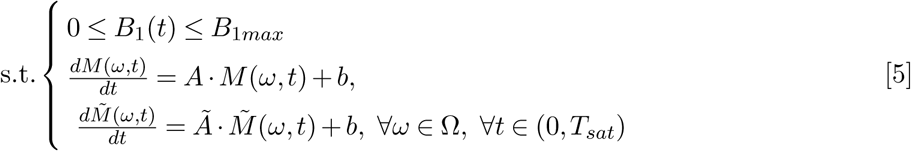

The main difference compared to the cost function in the paper is the third term, weighted by *σ*_3_. This term, depending on the choice of *n*, penalizes differences between the first pulse and other pulses in the train. For example, penalizing all pulses results in a train of identical pulses, similar to the single pulse presented in the paper (see Figure 10 (b, c)). Penalizing all trains except for the last one leads to the *N* + 1 configuration (d, e)), while penalizing every second train results in a pulse pair that can be applied in an alternating manner (f, g). Other configurations may also lead to favorable CEST spectra.

**Figure 10:**
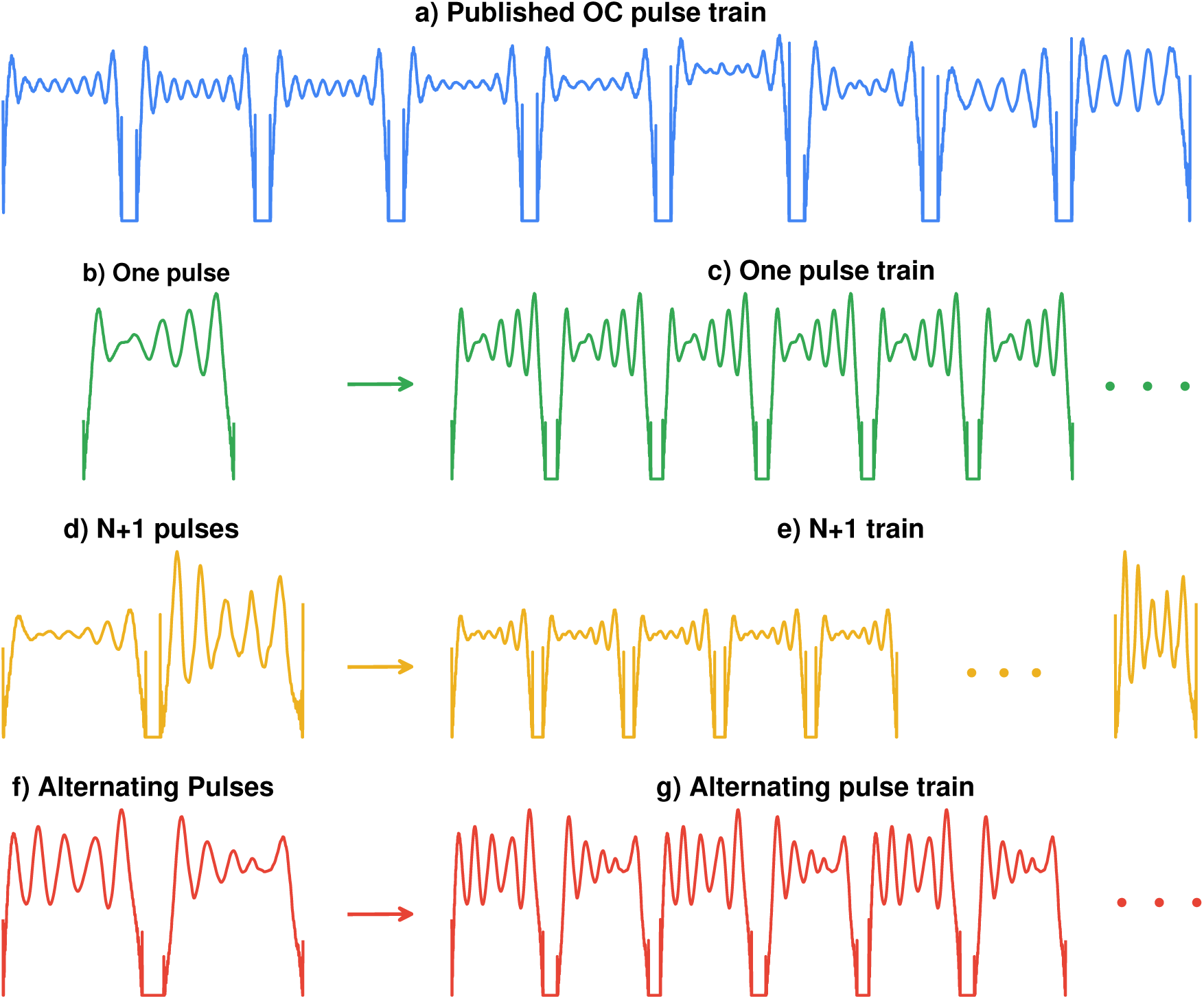
Schematic of different approaches to OC saturation.

This formulation of the cost functional is comparable to modifying the gradient to optimize only specific parts of the pulse train. However, this approach was easier to implement and experiment with. In particular, for very high values of *σ*_3_, the difference between the two approaches should be minimal.

The different OC saturation strategies were simulated with high-frequency sampling (0.01 ppm) to reveal sidebands and Rabi oscillations (see Figure 11(1-d)). The *N* + 1 and alternating approaches produce spectra that are nearly identical to the published pulse train. In contrast, the single-pulse approach, as used in the paper, results in slight oscillations in the water peak between −1 and 1 ppm. However, the overall performance in generating a CEST effect remains nearly identical (e).

**Figure 11:**
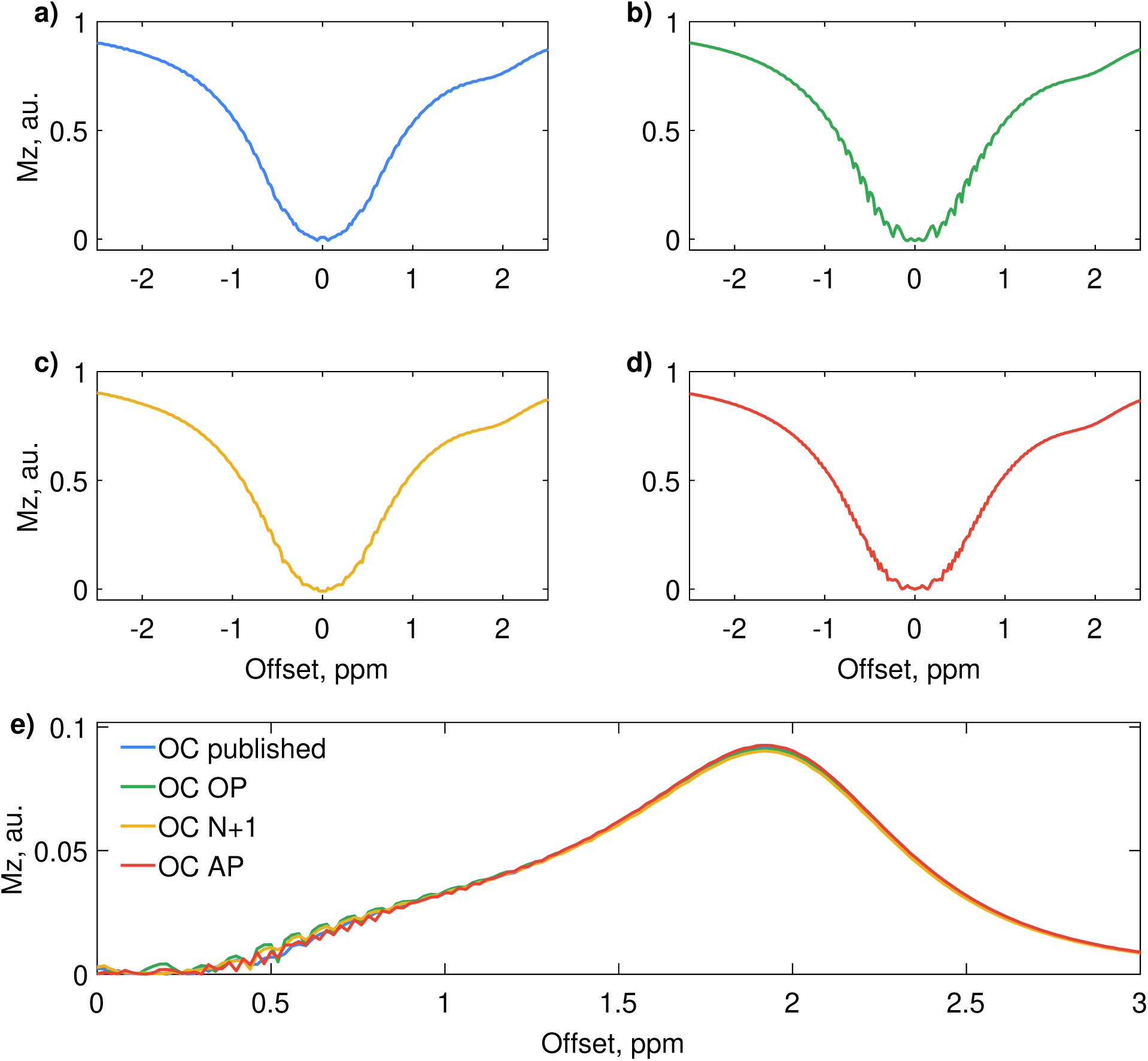
Results of simulations for the the saturation strategies of pulses presented in Figure 10. Simulation parameters are the same as described in the RF Pulse design section of the paper.

Oscillations in the water spectrum between −1 and 1 ppm could be detected in phantom measurements at a clinical 3 T scanner at a frequency sampling of 0.25 ppm (see Figure 12). The published OC pulse, the N+1 and the AP train produce smoother spectra than the one pulse. With an spoiler after each pulse, the spectrum of the OP could be smoothed. This artifacts depend on the *T*_2_ value.

**Figure 12:**
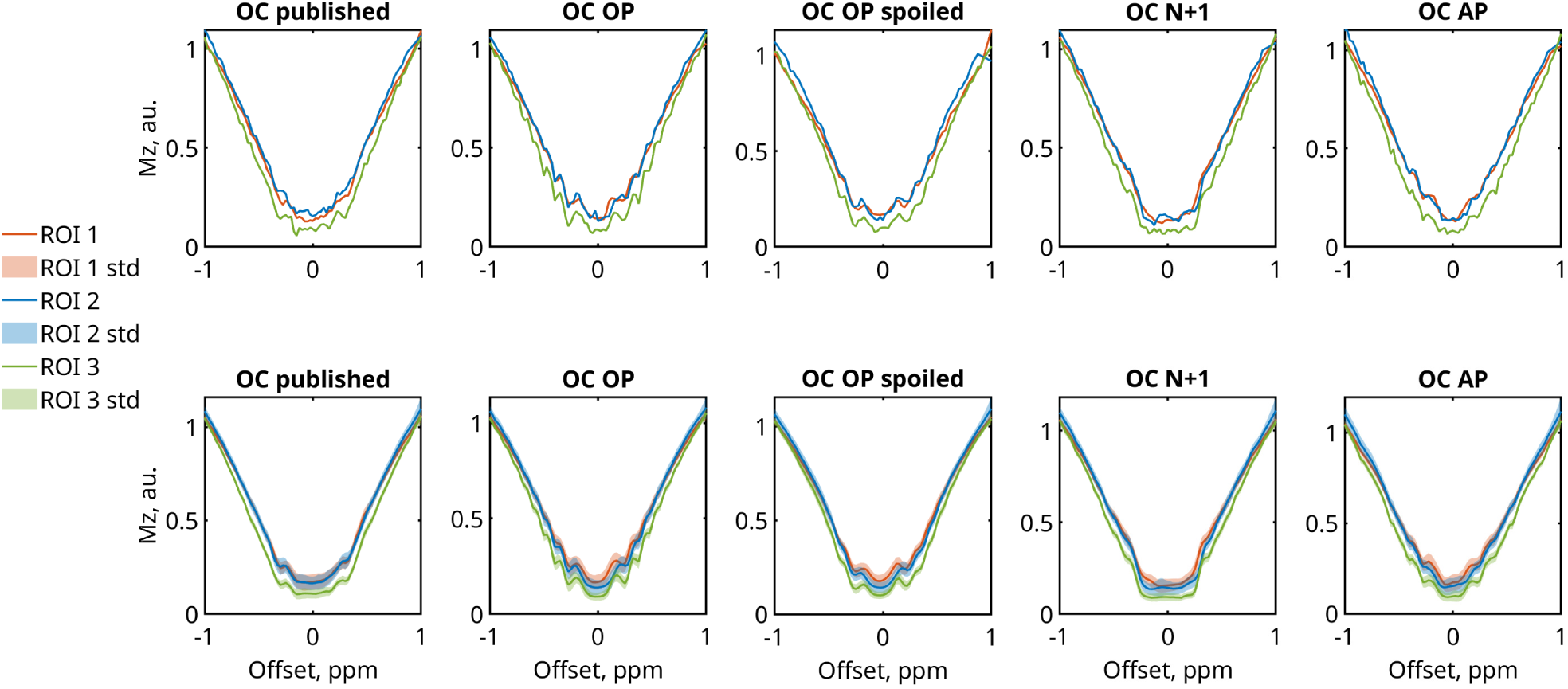
Phantom measurements in water phantoms with different OC saturation strategies presented in Figure 10 for differnt T2 values: ROI1 = 97 ms, ROI2 = 73 ms, ROI3 60 ms. First row single pixels, second row mean and std over ROI with several pixels.

## 13 Rabi oscillations and sidebands of Gauss and Fermi pulses

In Figure 13 it can be seen that other pulses also exhibit spectral artifacts depending on the pulse shape. Gaussian saturation generally behaves well. Especially for longer *t_d_*, the spectrum appears clean, as the adiabatic condition is fulfilled even for lower offsets. This well-behaved spectrum, however, comes at the cost of lower saturation efficiency. Shorter versions of well-performing pulses, such as Gaussian or windowed sinc, can introduce more sideband artifacts [2] and increased Rabi oscillations due to the faster rise in RF amplitude. Fermi pulses, which also have a steep RF onset, tend to produce much more Rabi oscillations and express strong sideband artifacts, particularly at short durations. However, they usually achieve higher saturation efficiency than Gaussian pulses. These Artifacts decay with the *T*_2_ value of the water pool, in this Figure 120 ms.

**Figure 13:**
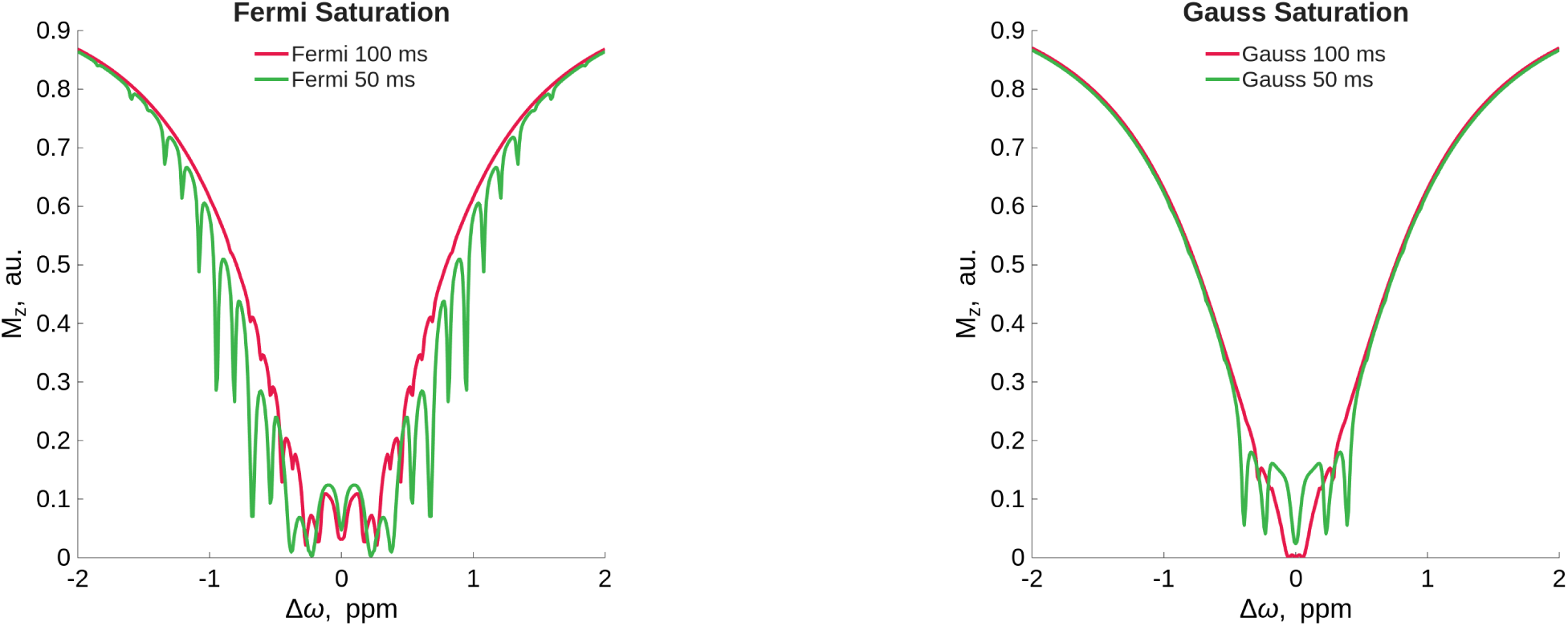
Artifacts apparent with simulations at high spectral resolution Δ*ω* = 0.01 ppm at 3 T.

## 14 CW vs short gaussian pulses in a multi-pool CEST peak

In Figure 14 the simulation of a 5 pool simulation can be seen, where short Gaussian saturation is depicted vs CW saturation. The Gaussian saturation were *t_d_* = 15.4 ms, DC = 90 % as in [48]. Here we simulate with following parameters: *T_sat_*= 2 s, *B*_1_*_RMS_* = µT (over train), CEST pool offsets = 3.3, 3.4, 3.5, 3.6 ppm, *T*_2_ of cest pools = 80 ms, exchange rates *k_s_w* = 25 Hz, CEST pool fraction rates = 0.0003, water relaxation *T*_1_ = 1.5 s, *T*_2_ = 100 ms.

**Figure 14:**
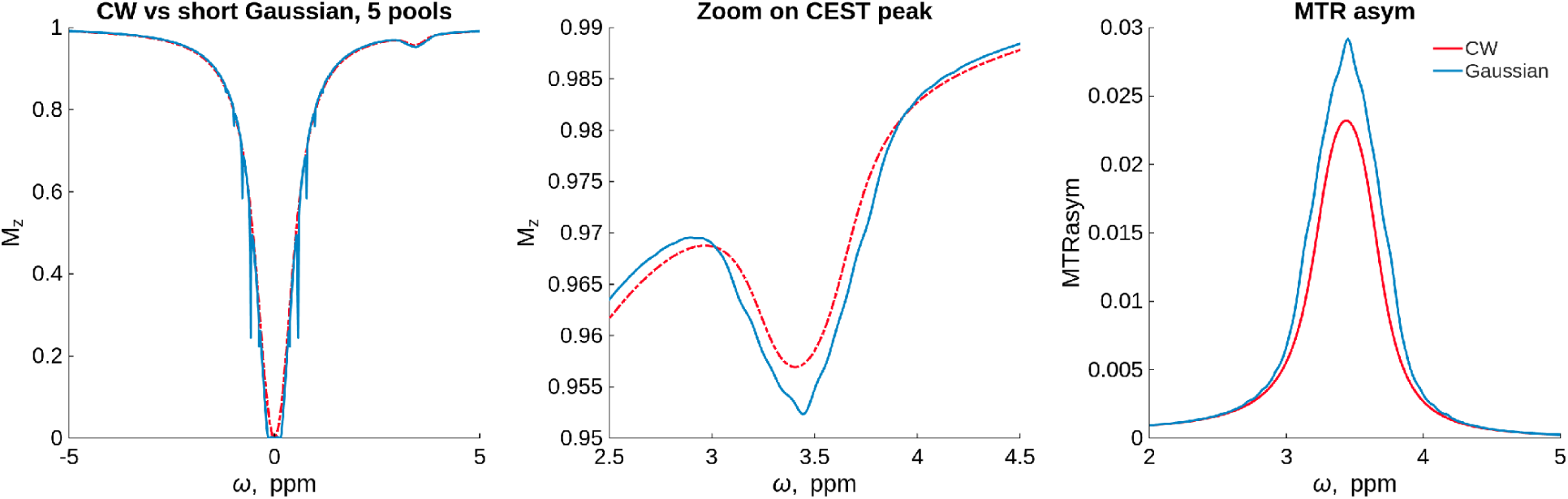
Multipool simulation for 7T. CW vs short Gaussian pulses. At exchange rate 25 Hz.

Interestingly, in this regime, the CEST peak is higher for Gaussian saturation compared to CW saturation. This effect occurs only at low exchange rates and low *B*_1RMS_, as shown in the simulations for different *B*_1RMS_ and exchange rates in Figure 15. Additionally, the presence of sidebands and oscillations at the CEST peak with Gaussian saturation is noteworthy.

**Figure 15:**
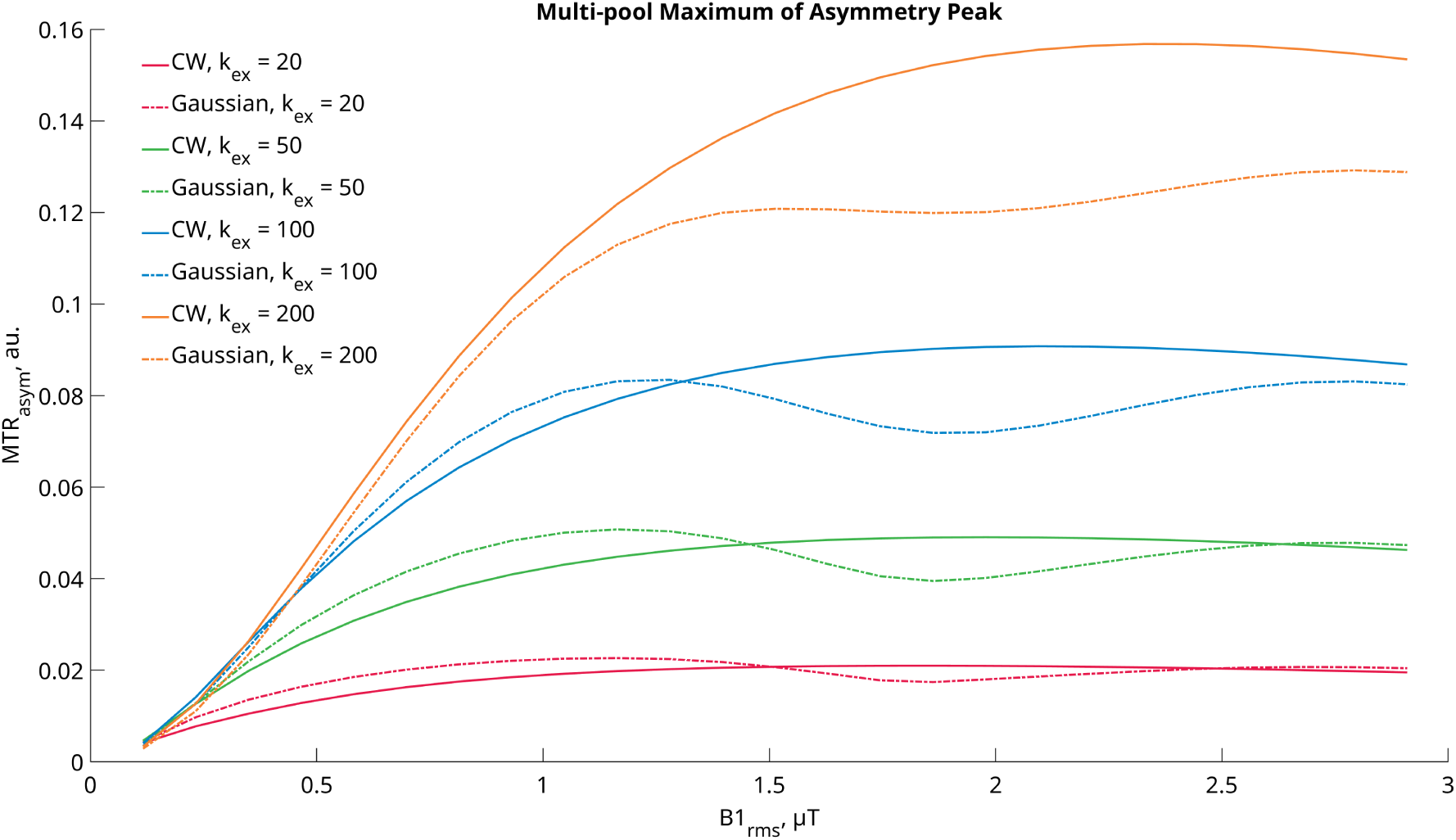
Multipool simulation for 7T. CW vs short Gaussian pulses. At different exchange rates and *B_RMS_*. In this simulations, the *MTR_asym_*peak of short Gaussian pulses is higher for smaller *B*_1_*_RMS_* levels.

Figure 15 further illustrates that for exchange rates of 200 Hz or higher, CW saturation again outperforms Gaussian saturation in this regime. Interestingly, this effect is more pronounced at 7 T than at 3 T (see Figure 16).

**Figure 16:**
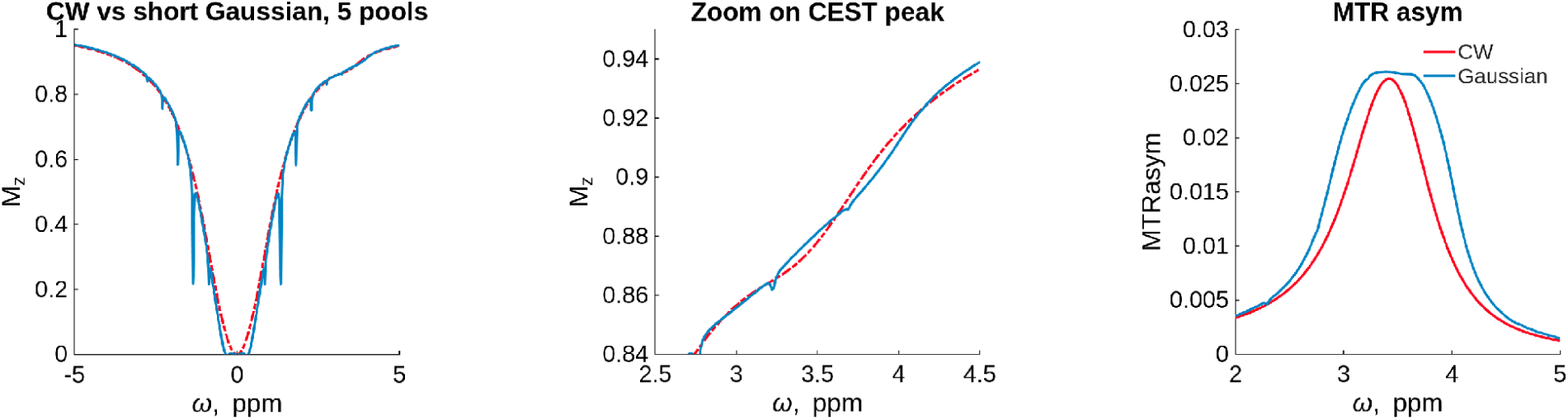
Multipool simulation for 3T. CW vs short Gaussian pulses. At exchange rate 25 Hz.

Of course, these simulations are based on arbitrary parameters and do not fully represent a real in vivo scenario. They are designed based on expected behavior for Amide Proton Transfer (APT) contrast. However, our goal here is to demonstrate that CW saturation does not necessarily yield the highest CEST image contrast in all cases.

### 14.1 OC for the multi pool case

For the multi pool case described in the previous section an OC pulse was designed. The target for the optimization were the spectrum of the short gaussian saturation. The optimization was not carried out for the whole spectrum but only between −4 and −2.5 and between 4 and 2.5 leaving out the water peak. the peak of the short gaussian pulses was not used as a target since we did not want to incorporate the sidebands in the optimization and we do not know what the perfect water spectrum in this case looks like.

As can be seen in Figure 17, the optimization lead to an amplitude comparable to the single pool case, but the optimal phase in this scenario is now not constant 0 anymore but exhibits a frequency sweep over 0.26 ppm. This makes sense since the collective amplitude of the single pulses can be increased by leveraging the a broader saturation in the spectrum.

**Figure 17:**
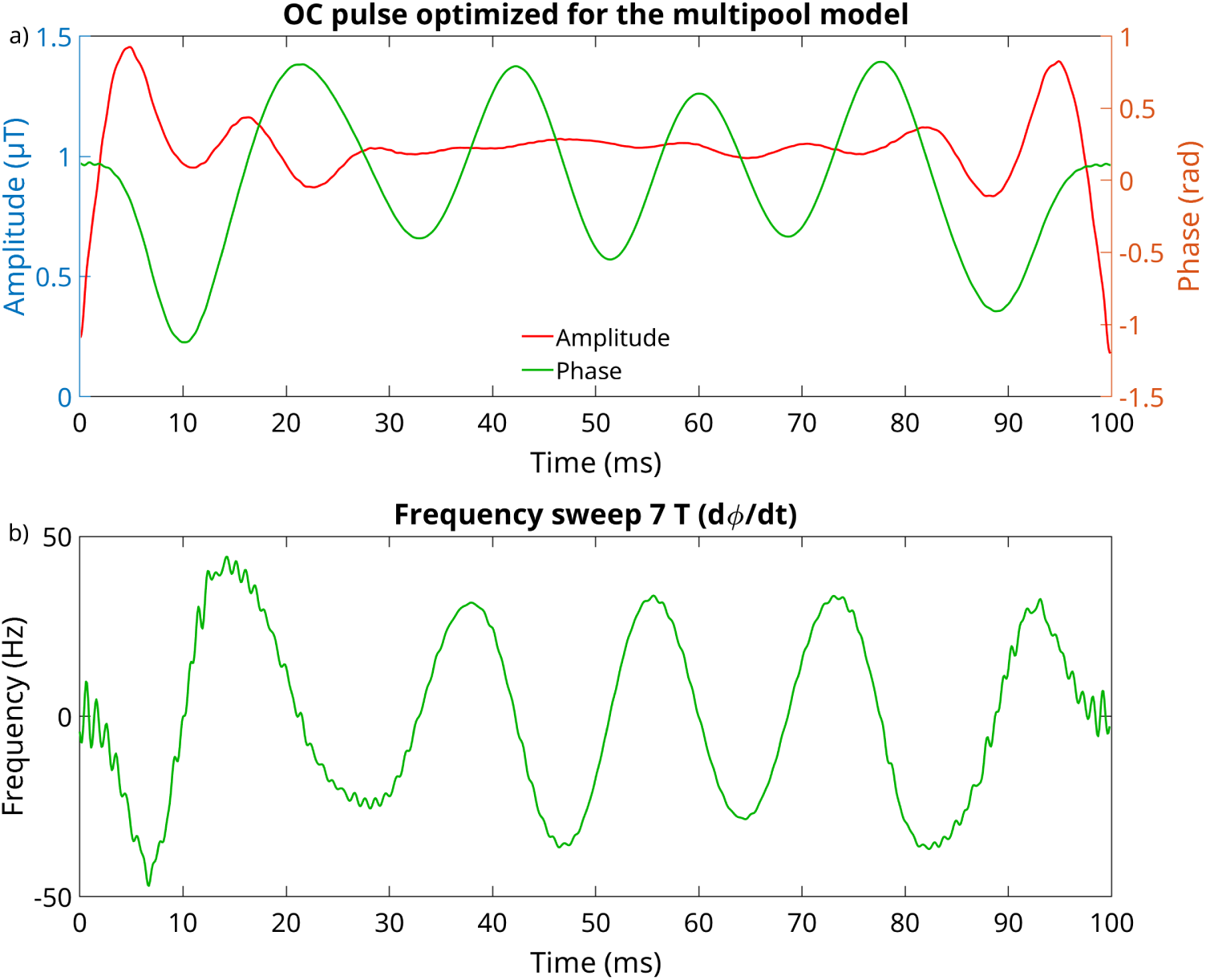
a) OC pulse designed for the multi pool case. b) Frequency sweep at 7 T.

The Multi pool OC pulse performs significantly better in simulations of the multipool case in comparison to CW saturation (Figure 18). And since the target was the spectrum of the short gaussian pulses the performance is comparable. with slightly higher contrast at 25 Hz. Interestingly the oscillations of the contrast over the *B*1*_RMS_* which can be seen with Gaussian saturation was not noticeable with the OC pulses.

**Figure 18:**
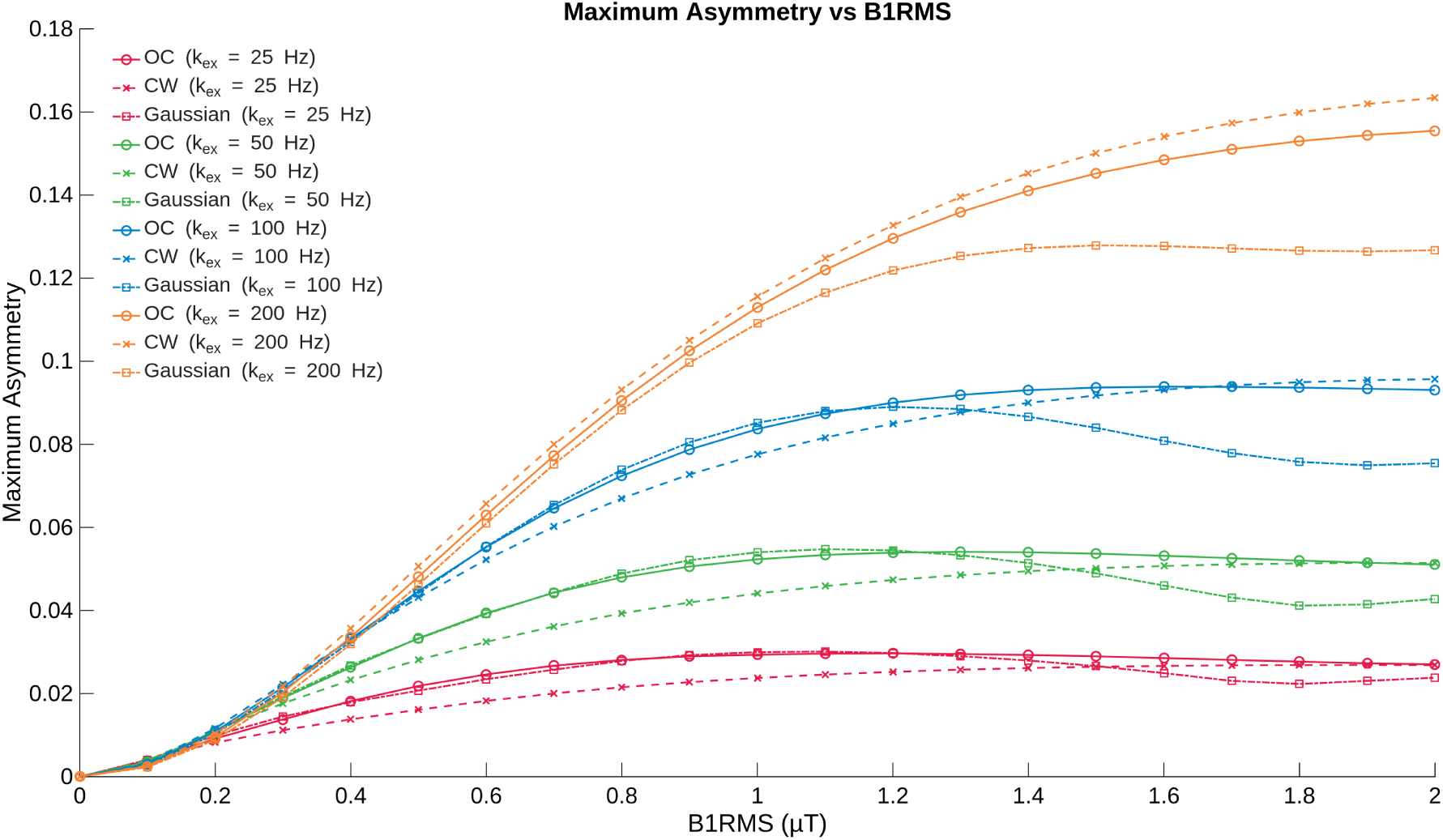
Multipool simulation for 7T.OC vs CW and short Gaussian pulses. At different exchange rates and *B_RMS_*. In this case the OC saturation was able to exceed CW contrast.

For the multi-pool case, an OC pulse was designed using the spectrum of short Gaussian saturation as the optimization target. The optimization was restricted to two spectral regions (−4 to −2.5 ppm and 2.5 to 4 ppm), excluding the water peak. The central peak of the Gaussian pulse was omitted to avoid sideband contributions and because the ideal spectrum in this context is unknown.

As shown in Figure 17, the optimized pulse exhibits an amplitude similar to the single-pool case, but with a non-zero phase featuring a frequency sweep over 0.31 ppm. The sweep range matches the simulated distribution of the four CEST peaks over a range of 0.3 ppm. This broader spectral coverage improves the collective saturation of multiple pools. Interestingly, the pulse starts at the center frequency, sweeps back and forth between negative and positive offsets, and returns to the center frequency at the end. The multi-pool OC pulse outperforms CW saturation in simulations (Figure 18) and achieves contrast comparable to Gaussian pulses, with slightly higher contrast at 25 Hz. Notably, the maximum of the asymmetry of the OC pulse outperformed the Gaussian saturation at higher *B*1_RMS_ values and follows the CW saturation more closely.

## 15 Frequency spectra of used pulses

The frequency spectra of a single 100 ms Gaussian, Fermi, Block and OC pulse are depicted in Figure 20. The frequency spectrum is generated by applying an FFT to the time amplitude signal. The bandwidth of the pulses is calculated using the FWHM. The Gaussian pulse has the highest bandwidth with 20 Hz, followed by the Fermi pulse with 17 Hz. The lowest FWHM values are found in the Block pulse and the OC pulse, with 12 and 13 Hz respectively. However, this analysis is only a rough estimate because it disregards the influence of the nonlinearity of the Bloch-McConnell equation on the z-spectrum.

**Figure 19:**
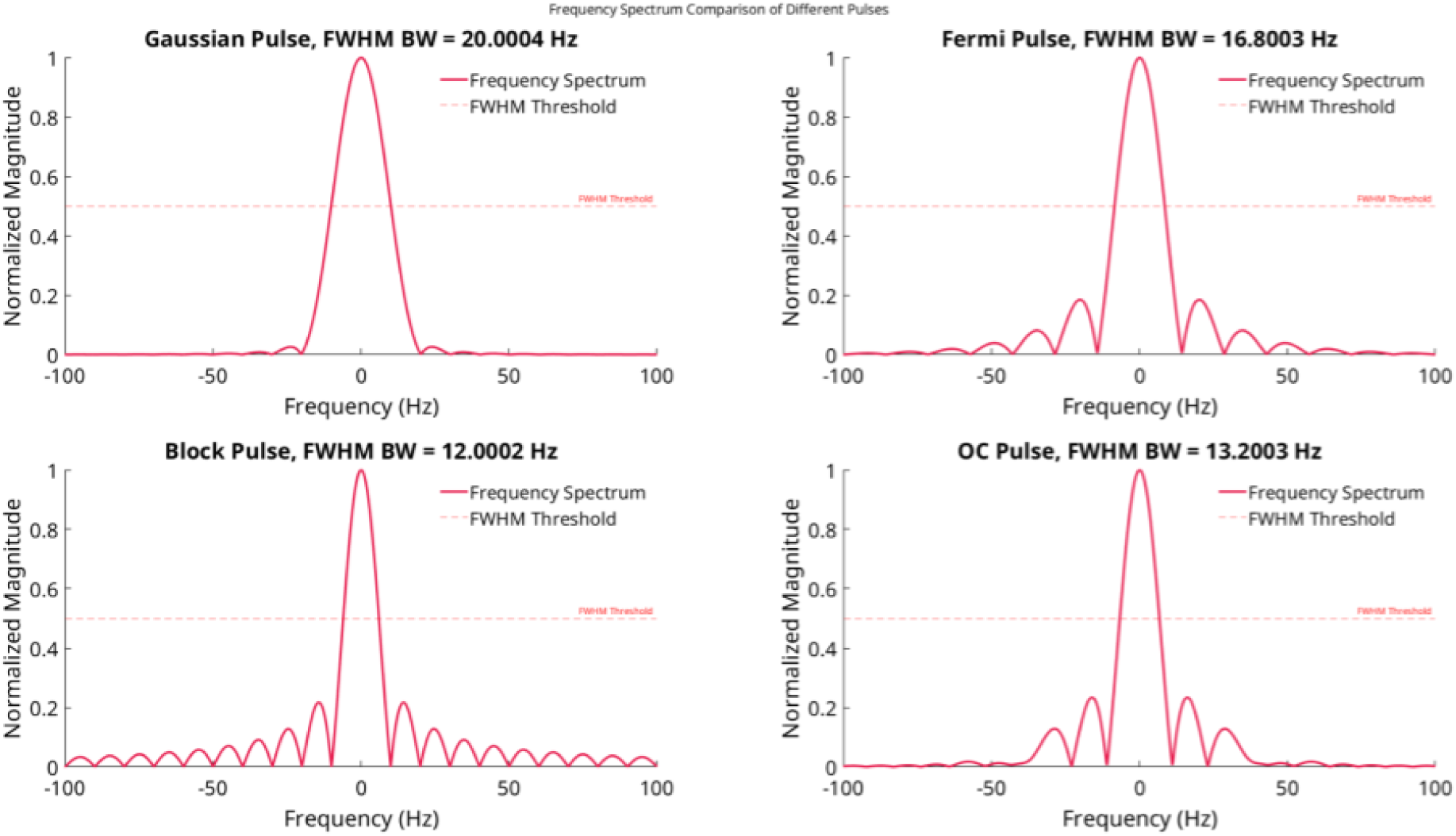
Frequency Spectra of a single 100 ms Gaussian, Fermi, Block and OC pulse

**Figure 20:**
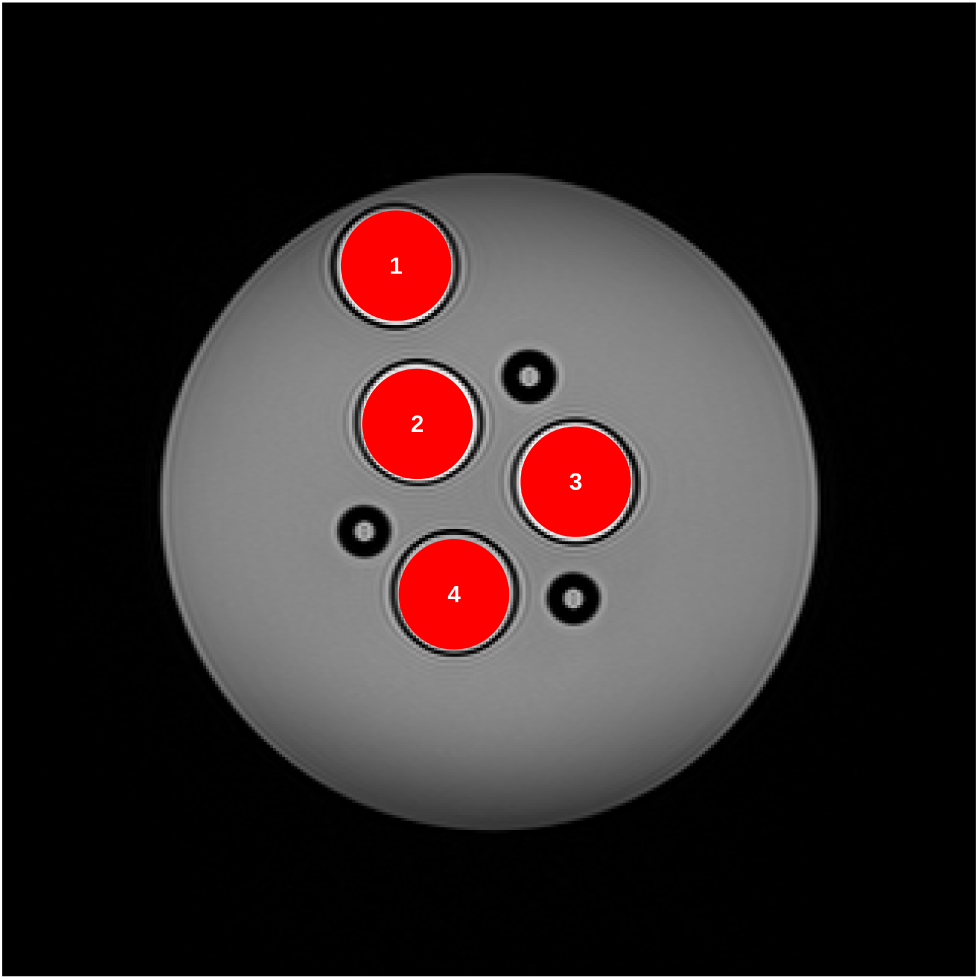
*T*_1_*_w_* weighted image of the CEST phantom, coronal slice with ROIs.

## 16 Phantom measurements

Example *T*_1_*_w_* image of the phantom used for phantom measurements. The phantom consisted of falcon tubes arranged by a 3D printed falcon tube holder in a water bath to make calibration of the scanner easier and to avoid big susceptibility jumps through air. The water outside the falcon tubes was without contrast agent.

## 17 Details to the optimization

The optimization process employs a set of stopping criteria to ensure efficient and accurate convergence. The algorithm terminates when one of the following conditions is met:

1. the relative gradient norm falls below a threshold of 10*^−^*^6^ times the initial gradient norm, indicating sufficient convergence
2. the absolute gradient norm becomes smaller than 10*^−^*^8^
3. the maximum number of iterations, set to 1000, is reached to prevent excessive computation
4. the trust region radius shrinks below 10*^−^*^12^, signaling optimization failure
5. no accepted solution is found within 100 consecutive iterations, indicating stagnation.

For testing the global convergence, the optimization was initialized with a completely random starting pulse. We generate *B*_1_(*t*) by drawing 1000 amplitude points from a uniform random distribution and scaling them to achieve RMS power *P* :

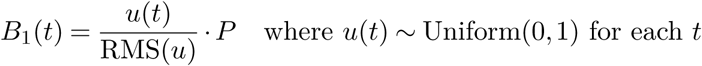

where 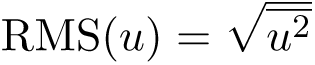 is the root-mean-square over all time points. This provides a stochastic initialization while maintaining the power constraint. The optimization results from multiple runs indicate robustness to local and global minima. The maximum difference in pulse amplitude is smaller than 1 nT, and the maximum difference in the simulated spectra is smaller than 7 × 10*^−^*^6^ % of the maximum water signal.

**Figure 21:**
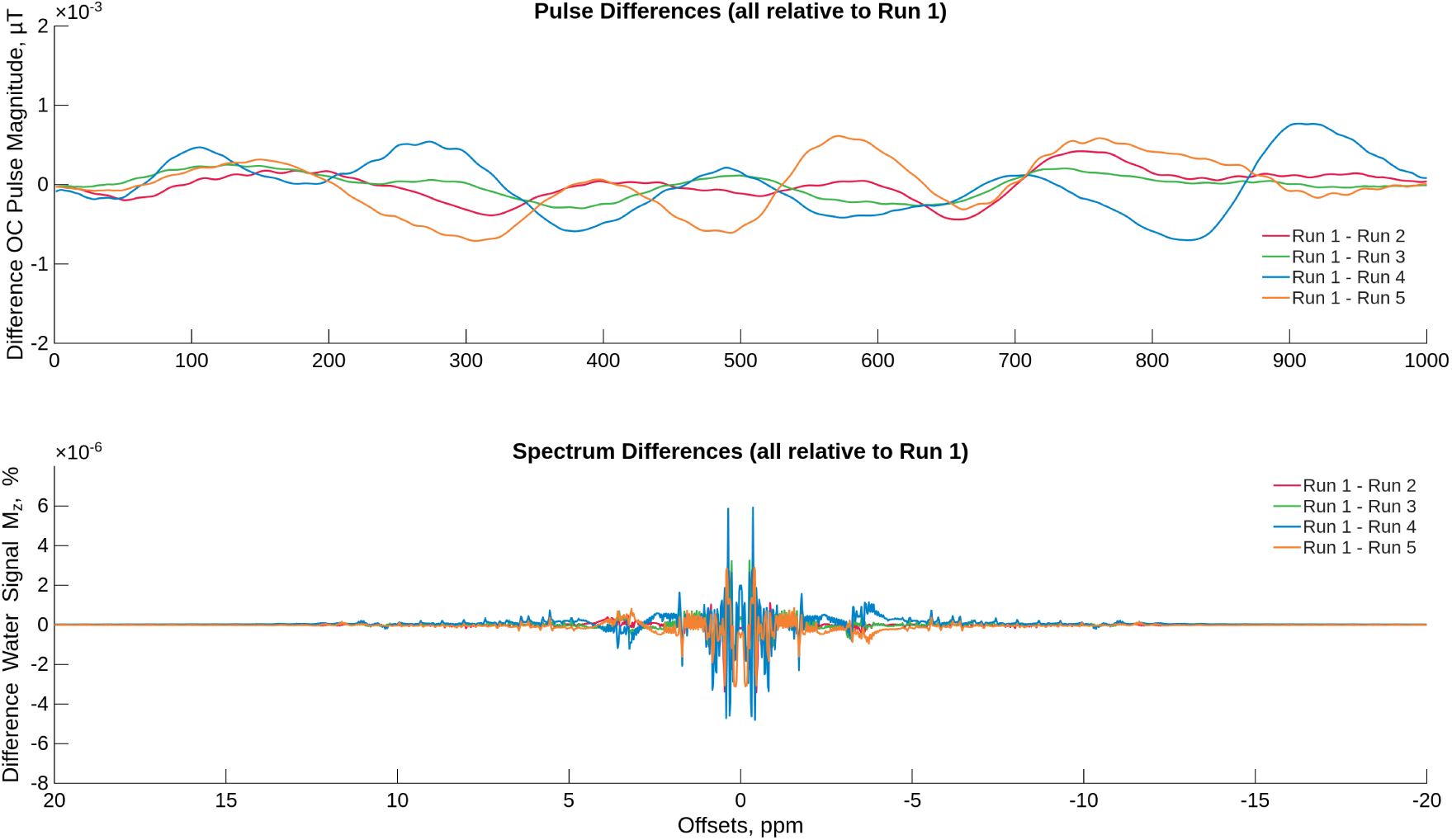
OC optimization results from 5 runs with random intiialization. Difference in pulse shape (top) and difference in simulated spectra (bottom).

## 18 In vivo thigh ***B*_0_** map

WASABI *B*_0_ map for the thigh measurement in Figure 22.

**Figure 22:**
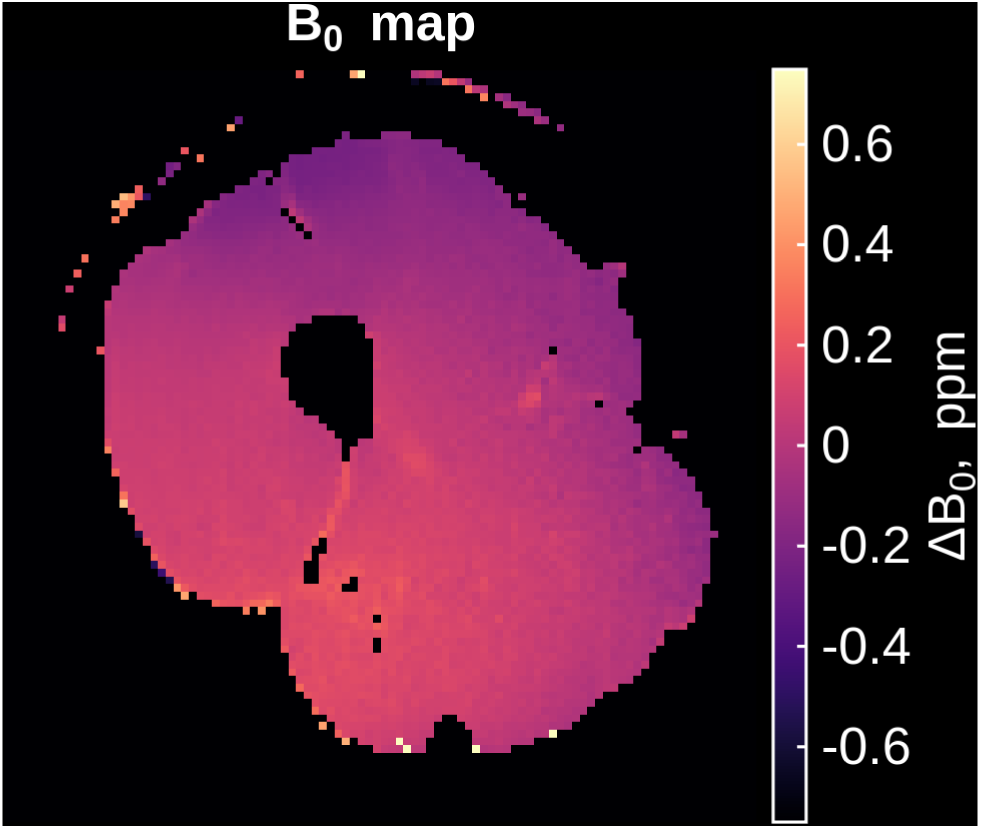
WASABI *B*_0_ map for the thigh measurement.

## 19 Performance of the different pulses in simulation

Pulseq CEST simulation of all 100 ms pulses used in the paper over different *B*1 scalings (Figure 23). The simulated values were:

**Figure 23:**
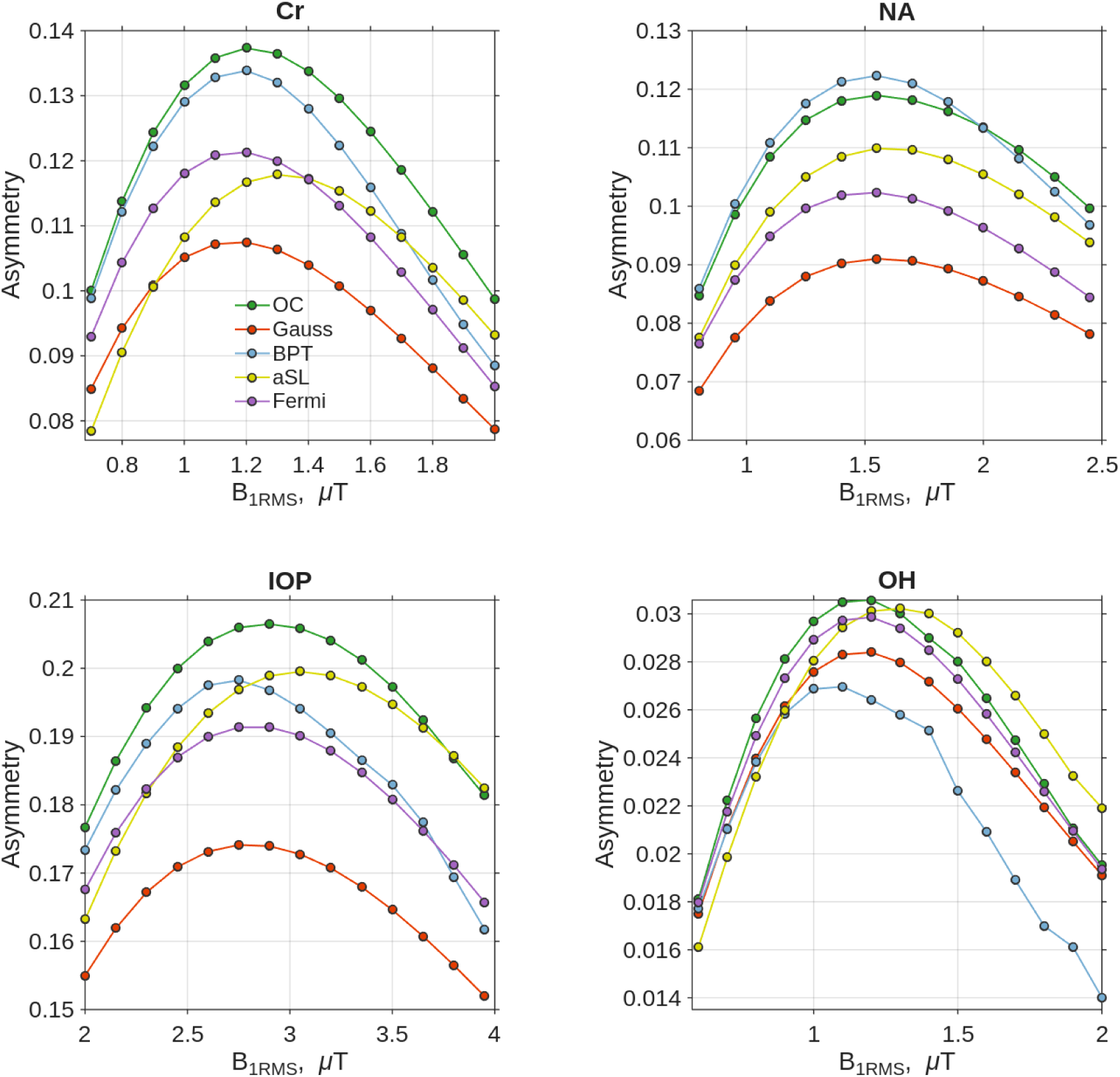
Pulseq CEST simulation: *MTR_asym_* for all pulse shapes used in manuscript, simulated for parameters expected in the phantom measurements. *B*1 level was adjusted to show the maximum point for every pulse

**Figure 24:**
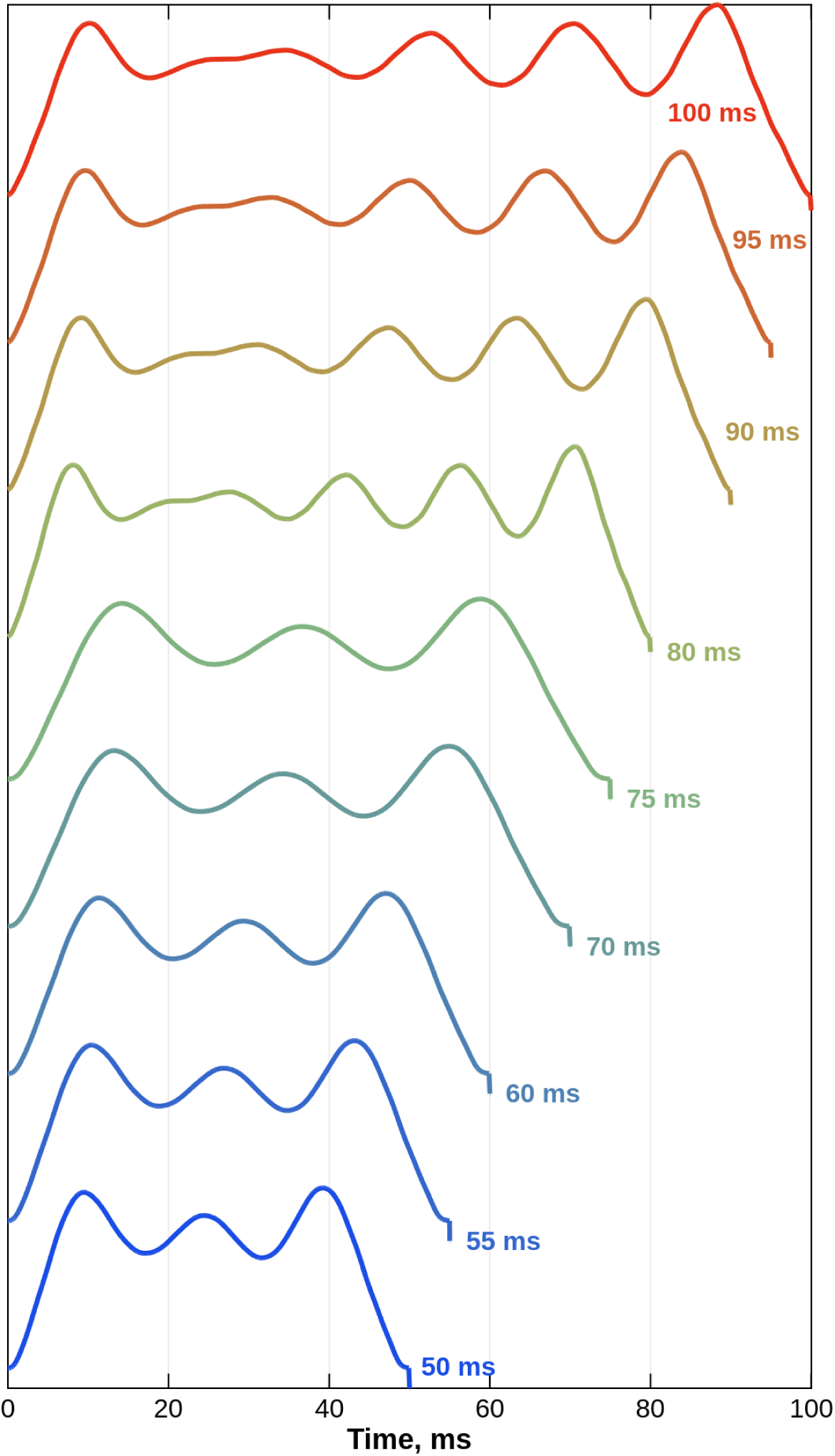
Pulse shapes stretched and compressed to different times *t_p_*.

Pulse train parameters: 8 pulses, *t_p_* = 100 ms, DC = 90 %.

BM pool model: water pool (*f* = 1.0, *T*_1_ = 1.2 s, *T*_2_ = 0.080 s) and four exchangeable pools: creatine (*f* = 0.0035, *T*_1_ = 1.2 s, *T*_2_ = 0.160 s, *k* = 250 Hz, Δ*ω* = 1.7 ppm), IOP (*f* = 0.0021, *T*_1_ = 1.2 s, *T*_2_ = 0.160 s, *k* = 1000 Hz, Δ*ω* = 4.2 ppm), NA (*f* = 0.0017, *T*_1_ = 1.2 s, *T*_2_ = 0.160 s, *k* = 250 Hz, Δ*ω* = 3.2 ppm), and OH (*f* = 0.002, *T*_1_ = 1.2 s, *T*_2_ = 0.160 s, *k* = 1000 Hz, Δ*ω* = 1.2 ppm).

## 20 Performance comparison of temporally stretched and compressed OC pulses

To minimize artifacts in the OC spectra shown in Figure 3 e,f,g,h of the paper, the 100 ms OC pulse was used for scalings between 100 and 76 ms. For scalings between 75 and 50 ms the 50 ms pulse was used. The correct pulses and interpolations are implemented in the makeOCPulse in pulseq CEST and is automatically selected based on the user input of td.

Pulseq CEST simulation for the creatine parameters in section 19 for different pulse times in 1 s pulse train with DC of approximately 90 %.

For different pulse times the OC shows constant highest saturation (Figure 25). The 100 ms pulse generates slightly higher contrast than the 50 ms, independent of the scaling in time.

**Figure 25:**
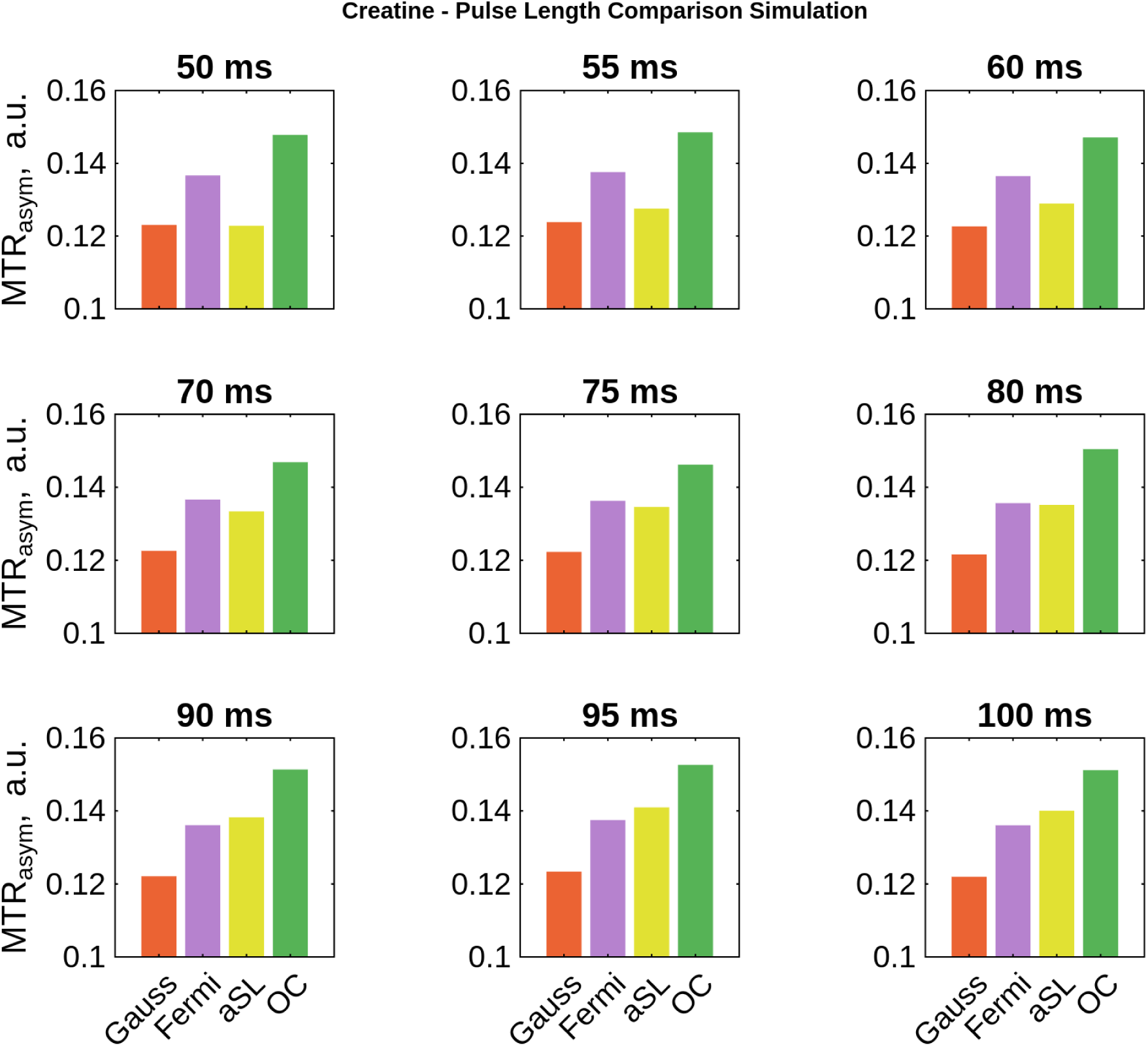
*MTR_asym_* for different pulse times and Gauss, Fermi, aSL and OC pulse simulated for the creatine pahntom.

High resolution (0.01 ppm) spectra for all pulse shapes and different pulse times can be seen in Figure 26. These spectra are simulated with a *B*_0_ inhomogeneity of 0.1 ppm.

**Figure 26:**
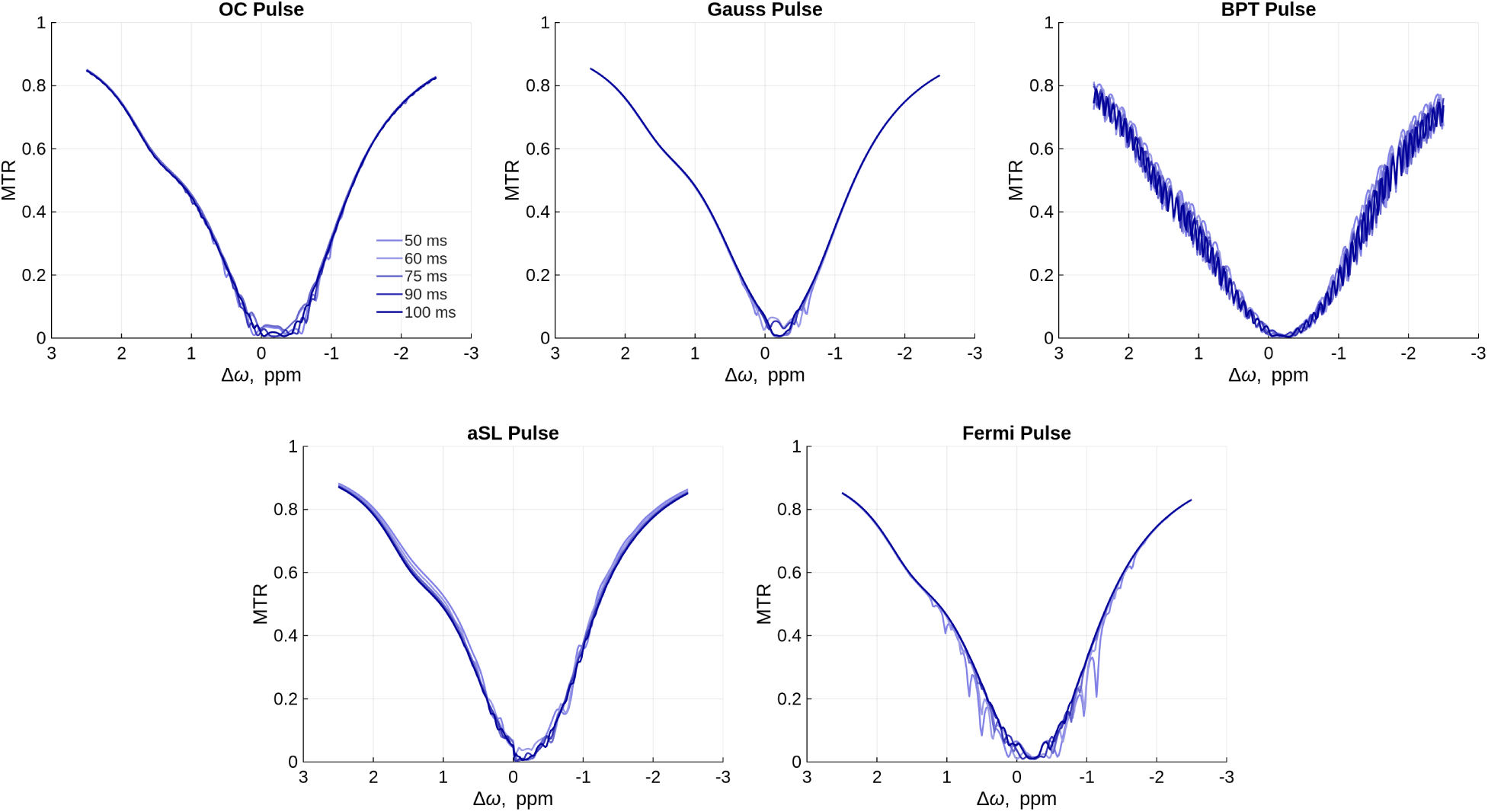
Spectra simulated with different pulse lengths and a high frequency resolution of 0.01 ppm and a *B*_0_ inhomogeneity of 0.1 ppm.

The pulse train parameters for both Figures 25, 26 were chosen to have a constant DC of 91 % a *T_sat_* of approximately 1 s and a *B*_1*rms*_ of 1 µT.

## 21 Lorentzian fitting

Parameter maps for the APT brain Lorentzian fitting can be seen in Figure 28. Water and MT were fitted with a 2 pool Lorentzian model. For the fit a pixel wise Lorentzian model was fitted with a least squares algorithm. Both saturation lead to approximately the same MT maps. The on resonant saturation of the OC was higher than the Gaussian saturation. This is in agreement with simulations. The OC APT contrast generated is approximately 30 % higher than with Gaussian saturation.

**Figure 27:**
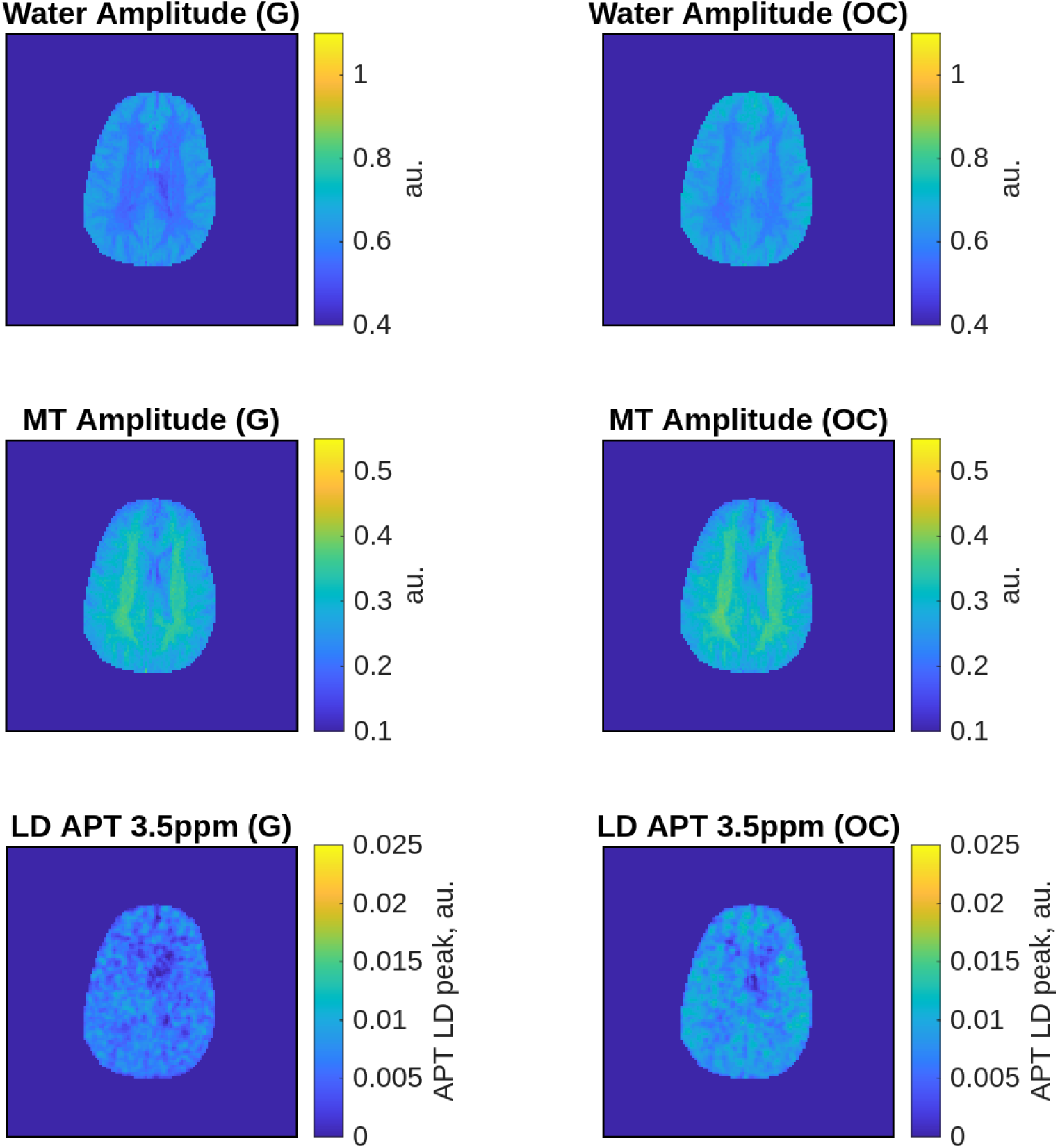
Parameter maps from the Lorentzian fitting. And the Lorentzian difference maps resembling the extracted APT contrast.

**Figure 28:**
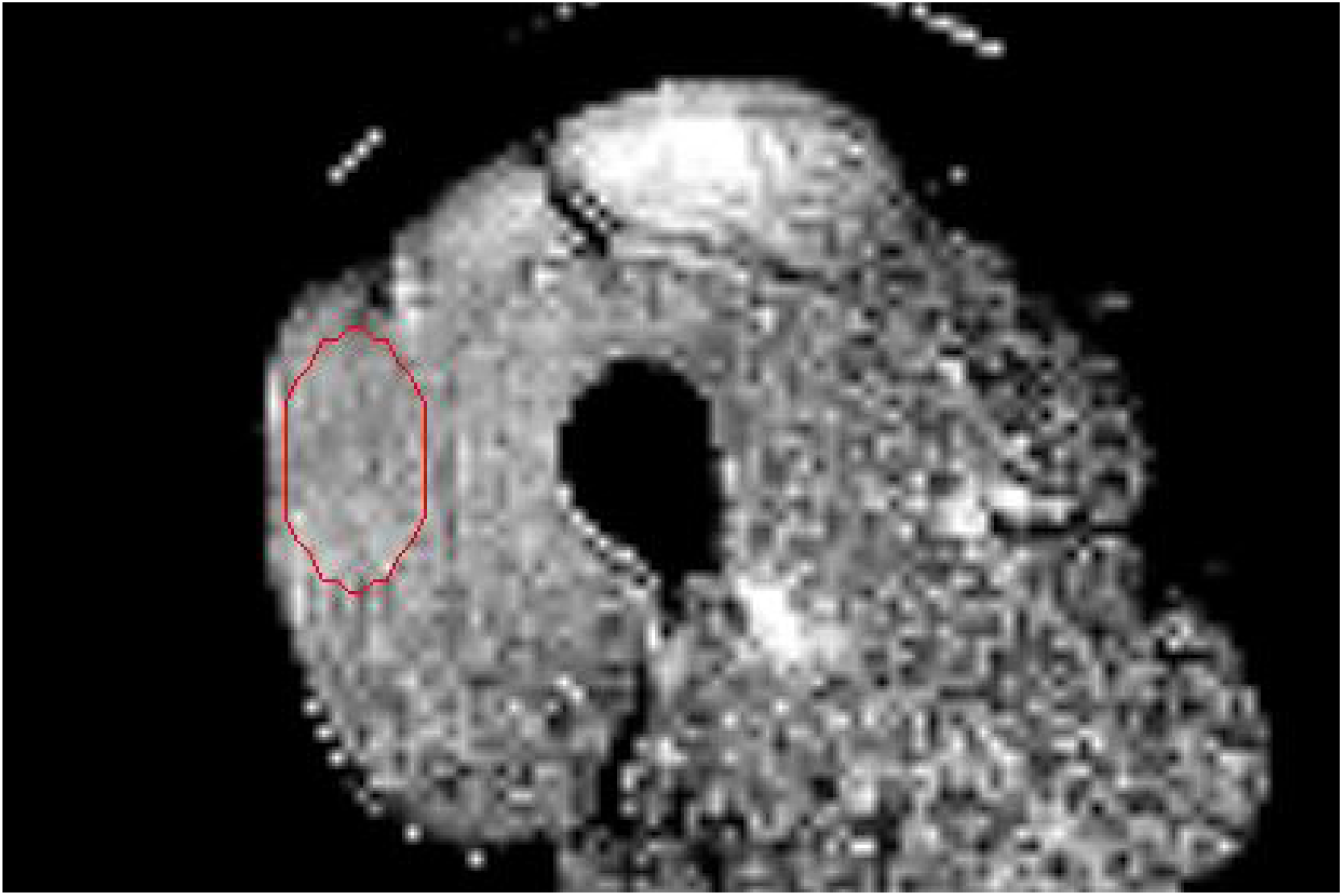
ROI in the PCA-denoised and *B*_0_-corrected OC *MTR_asym_* image used for the calculation of the SNR.

## 22 ROI for the SNR calculation

## Notes

### Competing Interest Statement

The authors have declared no competing interest.

