## Supporting Information for "Generalization of Optimal Control Saturation Pulse Design for Robust and High CEST Contrast"

### 1 Phantom measurements for 50 ms pulses

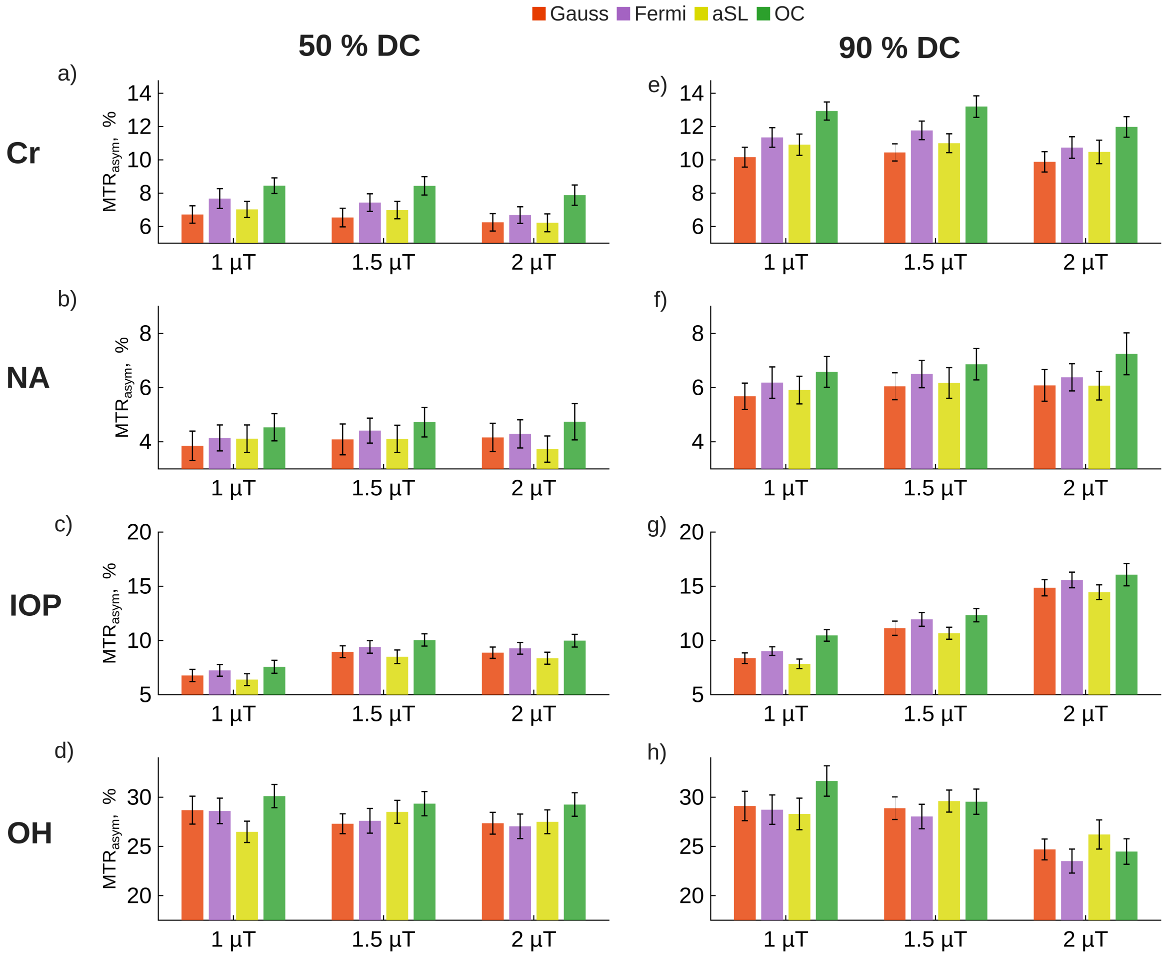


Figure S1: Phantom measurements comparing the performance of Gaussian, Fermi, adiabatic Spin Lock (aSL), and Optimal Control (OC) saturation pulses. All pulses were applied for 50 ms with a total saturation time of 1 s, using 50 % duty cycle (DC) in (a-d) and 90 % DC in (e-h). Saturation was performed at $B_1$ RMS of 1, 1.5, and 2 µT. The phantom consisted of four Falcon tubes, each containing one of the following: Cr, NA, IOP, and sucrose (OH).

### 2 Interleaved measurements, magnetization after each saturation pulse in a pulse train

Figure S2 depicts the magnetization after each pulse in a pulse train of 20 OC pulses, each 100 ms long with a 90% DC. With the exception of the $T_2$-dependent Rabi oscillations in the water peak (simulated with $T_2 = 60$ ms), the magnetization remains smooth. This is crucial for interleaved measurements where an artifact-free magnetization and independence from the preceding pulse are required, as any measurement might alter the previously prepared magnetization and compromise the outcome.

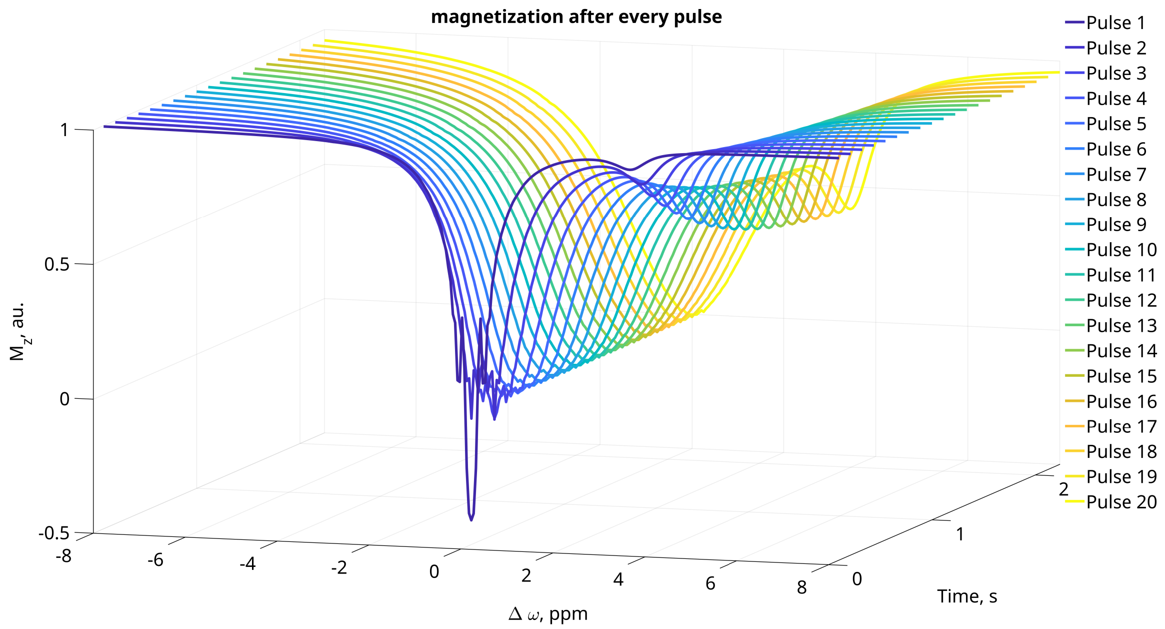


Figure S2: Waterfall plot for the magnetization over time. The Magnetization is measured after every pulse.

#### 3 Alternative formulation of cost functional

Since optimizing 1000 free parameters for $B_1(t)$ over 100 ms did not yield a spectrum comparable to that obtained with 9000 free parameters over a 1 s pulse train, we introduced a modified cost functional:

$$\begin{aligned} \min_{B_1(t)} J(B_1(t), M_z(\omega,t), \tilde{M}_z(\omega,t)) = \\ & \frac{\alpha}{2} \int_{t=0}^{T_{sat}} B_1(t)^2 dt \\ & + \frac{\sigma_1}{p} \sum_{\omega} \left| \frac{M_z(\omega, T_{sat}) - M_{zdes}(\omega)}{\epsilon} \right|^p \\ & + \frac{\sigma_2}{p} \sum_{\omega} \left| \frac{\tilde{M}_z(\omega, T_{sat}) - \tilde{M}_{zdes}(\omega)}{\epsilon} \right|^p \\ & + \frac{\sigma_3}{2} \sum_{n=1}^{\mathrm{R}-1} \sum_{i=1}^{T_{\mathrm{on}}} \left( B_1(t_i) - B_1(t_{i+n \cdot T_{\mathrm{on}}}) \right)^2, \end{aligned} \quad [1]$$

$$\text{s.t.} \begin{cases} 0 \leq B_1(t) \leq B_{1max} \\ \frac{dM(\omega,t)}{dt} = A \cdot M(\omega,t) + b, \\ \frac{d\tilde{M}(\omega,t)}{dt} = \tilde{A} \cdot \tilde{M}(\omega,t) + b, \ \forall \omega \in \Omega, \ \forall t \in (0, T_{sat}) \end{cases} \quad [2]$$

The main difference compared to the cost function in the paper is the third term, weighted by $\sigma_3$. This term, depending on the choice of $n$, penalizes differences between the first pulse and other pulses in the train. For example, penalizing all pulses results in a train of identical pulses, similar to the single pulse presented in the paper (see Figure S3 (b, c)). Penalizing all trains except for the last one leads to the $N+1$ configuration (d, e)), while penalizing every second train results in a pulse pair that can be applied in an alternating manner (f, g). Other configurations may also lead to favorable CEST spectra.

This formulation of the cost functional is comparable to modifying the gradient to optimize only specific parts of the pulse train. However, this approach was easier to implement and experiment with. In particular, for very high values of $\sigma_3$, the difference between the two approaches should be minimal.

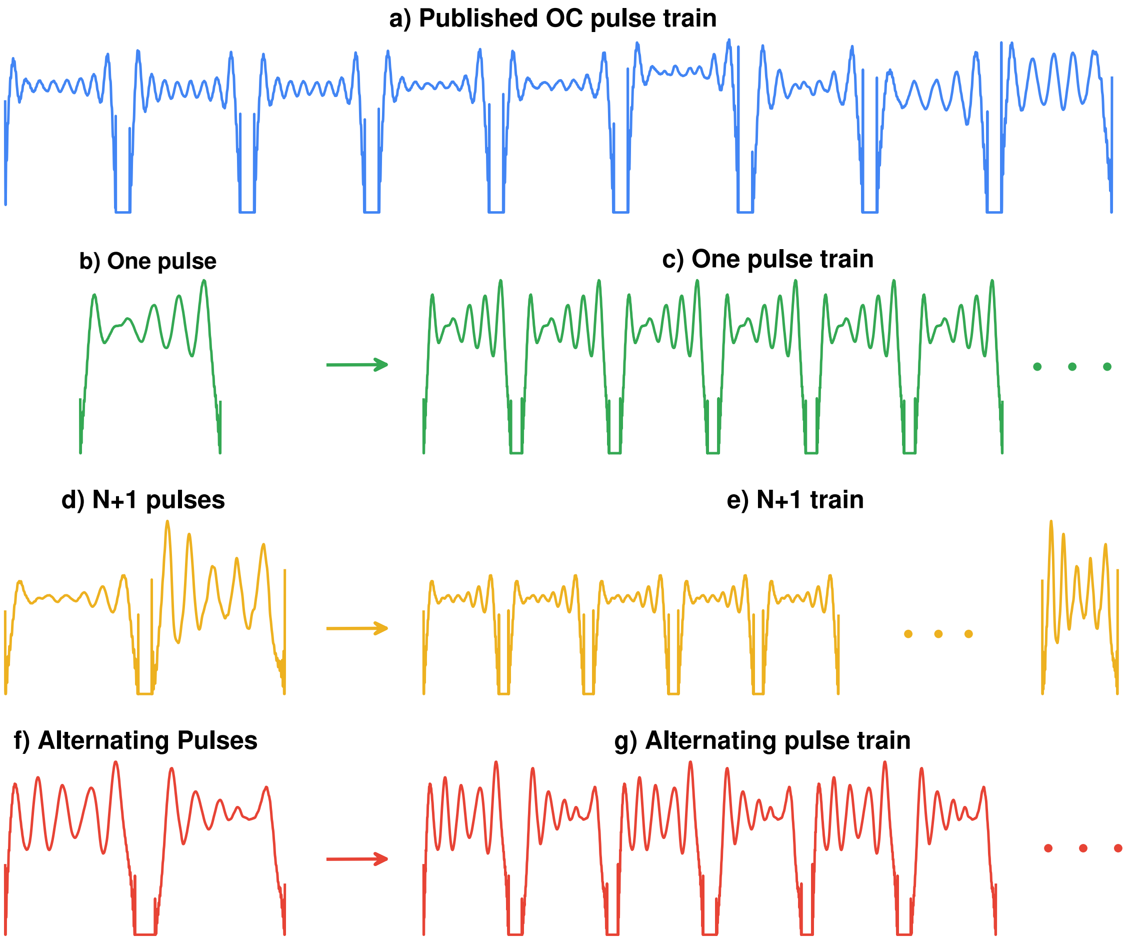


Figure S3: Schematic of different approaches to OC saturation.

The different OC saturation strategies were simulated with high-frequency sampling (0.01 ppm) to reveal sidebands and Rabi oscillations (see Figure S4(1-d)). The $N+1$ and alternating approaches produce spectra that are nearly identical to the published pulse train. In contrast, the single-pulse approach, as used in the paper, results in slight oscillations in the water peak between -1 and 1 ppm. However, the overall performance in generating a CEST effect remains nearly identical (e).

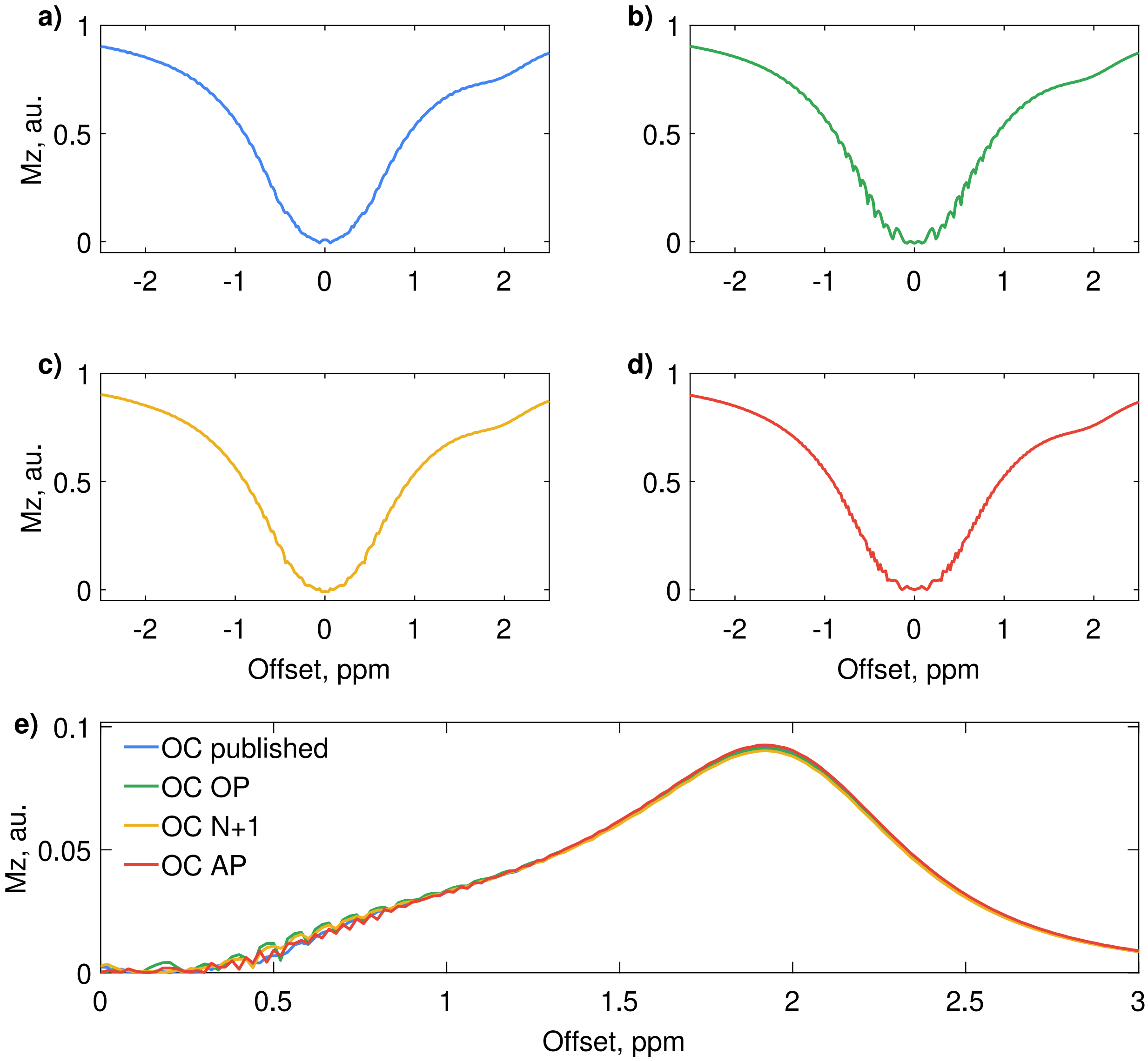


Figure S4: Results of simulations for the the saturation strategies of pulses presented in Figure S3. Simulation parameters are the same as described in the RF Pulse design section of the paper.

Oscillations in the water spectrum between -1 and 1 ppm could be detected in phantom measurements at a clinical 3 T scanner at a frequency sampling of 0.25 ppm (see Figure S5). The published OC pulse, the N+1 and the AP train produce smoother spectra than the one pulse. With an spoiler after each pulse, the spectrum of the OP could be smoothed. This artifacts depend on the $T_2$ value.

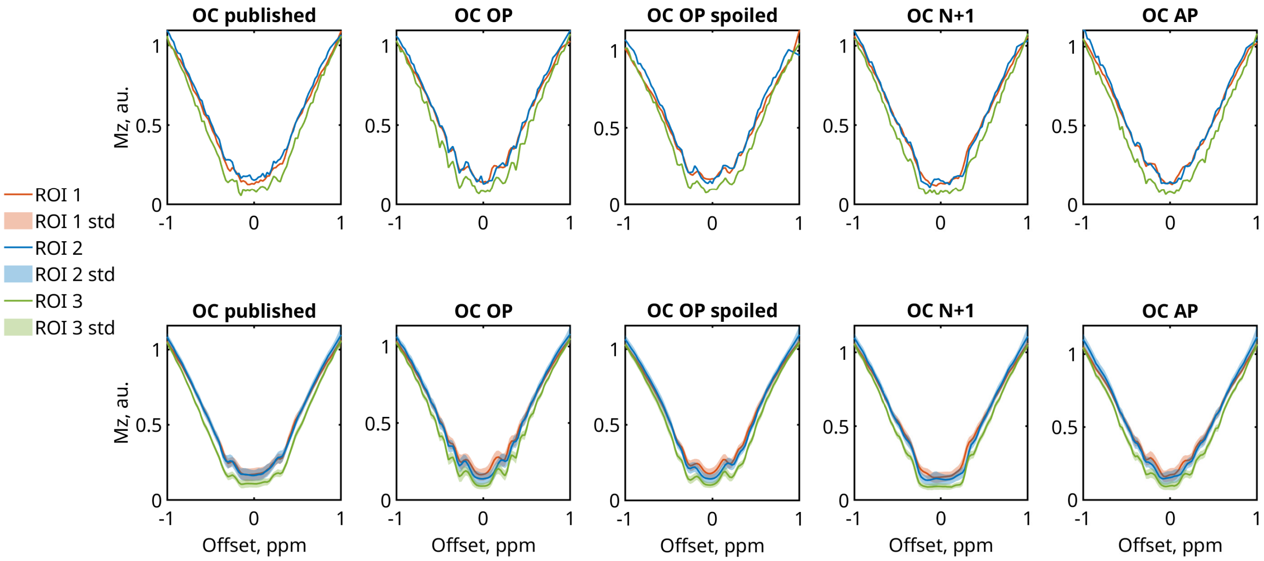


Figure S5: Phantom measurements in water phantoms with different OC saturation strategies presented in Figure S3 for differnt T2 values: ROI1 = 97 ms, ROI2 = 73 ms, ROI3 60 ms. First row single pixels, second row mean and std over ROI with several pixels.

### 4 Rabi oscillations and sidebands of Gauss and Fermi pulses

In Figure S6 it can be seen that other pulses also exhibit spectral artifacts depending on the pulse shape. Gaussian saturation generally behaves well. Especially for longer $t_d$, the spectrum appears clean, as the adiabatic condition is fulfilled even for lower offsets. This well-behaved spectrum, however, comes at the cost of lower saturation efficiency. Shorter versions of well-performing pulses, such as Gaussian or windowed sinc, can introduce more sideband artifacts [1] and increased Rabi oscillations due to the faster rise in RF amplitude. Fermi pulses, which also have a steep RF onset, tend to produce much more Rabi oscillations and express strong sideband artifacts, particularly at short durations. However, they usually achieve higher saturation efficiency than Gaussian pulses. These Artifacts decay with the $T_2$ value of the water pool, in this Figure 120 ms.

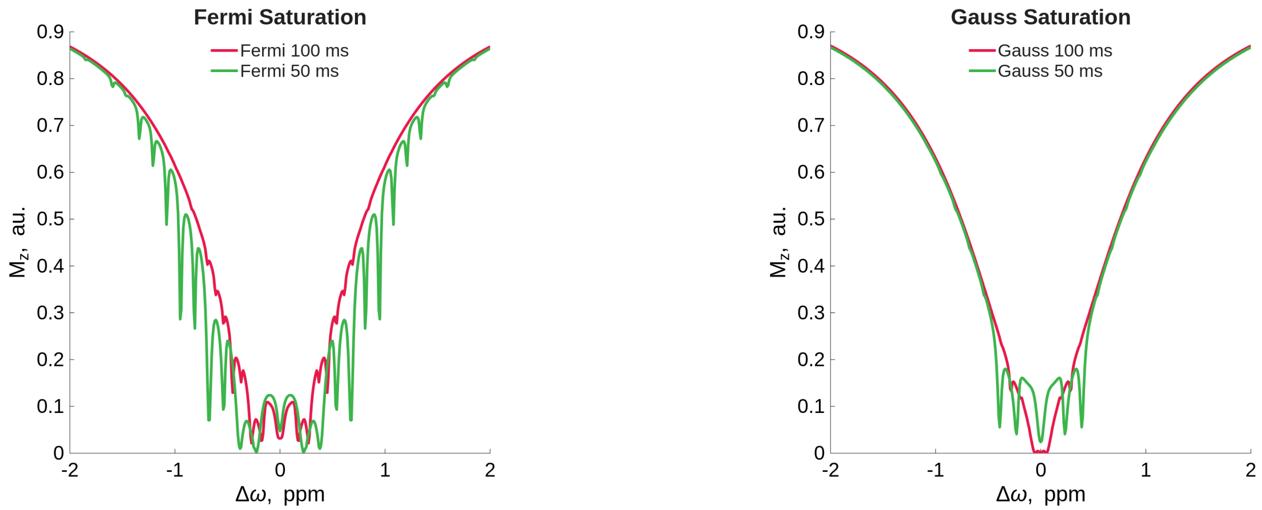


Figure S6: Artifacts apparent with simulations at high spectral resolution $\Delta\omega = 0.01$ ppm at 3 T.

### 5 CW vs short gaussian pulses in a multi-pool CEST peak

In Figure S7 the simulation of a 5 pool simulation can be seen, where short Gaussian saturation is depicted vs CW saturation. The Gaussian saturation were $t_d = 15.4$ ms, DC $= 90$ % as in [2]. Here we simulate with following parameters: $T_{sat} = 2$ s, $B_{1RMS} =$ µT (over train), CEST pool offsets $=$ 3.3, 3.4, 3.5, 3.6 ppm, $T_2$ of cest pools $= 80$ ms, exchange rates $k_s w = 25$ Hz, CEST pool fraction rates $= 0.0003$, water relaxation $T_1 = 1.5$ s, $T_2 = 100$ ms.

Interestingly, in this regime, the CEST peak is higher for Gaussian saturation compared to CW saturation. This effect occurs only at low exchange rates and low $B_{1\mathrm{RMS}}$, as shown in the simulations for different $B_{1\mathrm{RMS}}$ and exchange rates in Figure S8. Additionally, the presence of sidebands and oscillations at the CEST peak with Gaussian saturation is noteworthy.

Figure S8 further illustrates that for exchange rates of 200 Hz or higher, CW saturation again outperforms Gaussian saturation in this regime. Interestingly, this effect is more pronounced at 7 T than at 3 T (see Figure S9).

Of course, these simulations are based on arbitrary parameters and do not fully represent a real in vivo scenario. They are designed based on expected behavior for Amide Proton Transfer (APT) contrast. However, our goal here is to demonstrate that CW saturation does not necessarily yield the highest CEST image contrast in all cases.

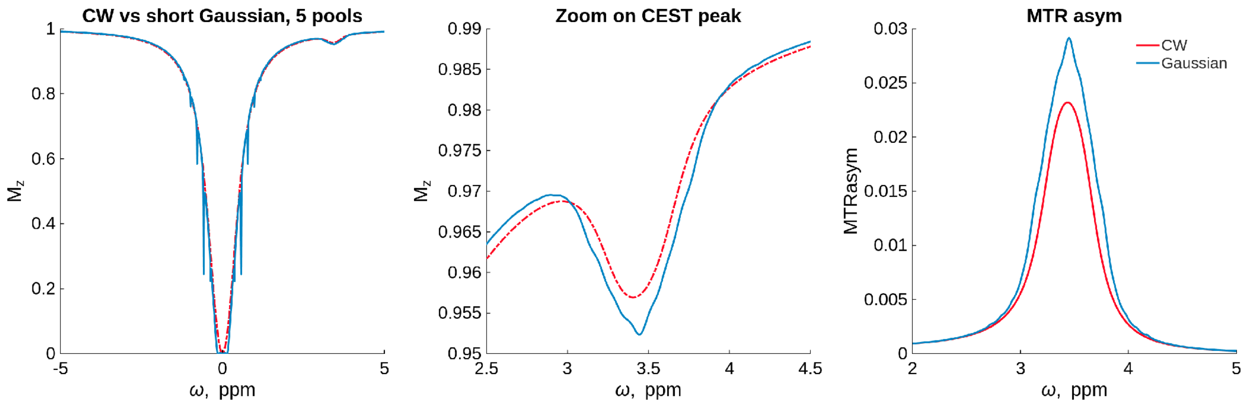


Figure S7: Multipool simulation for 7T. CW vs short Gaussian pulses. At exchange rate 25 Hz.

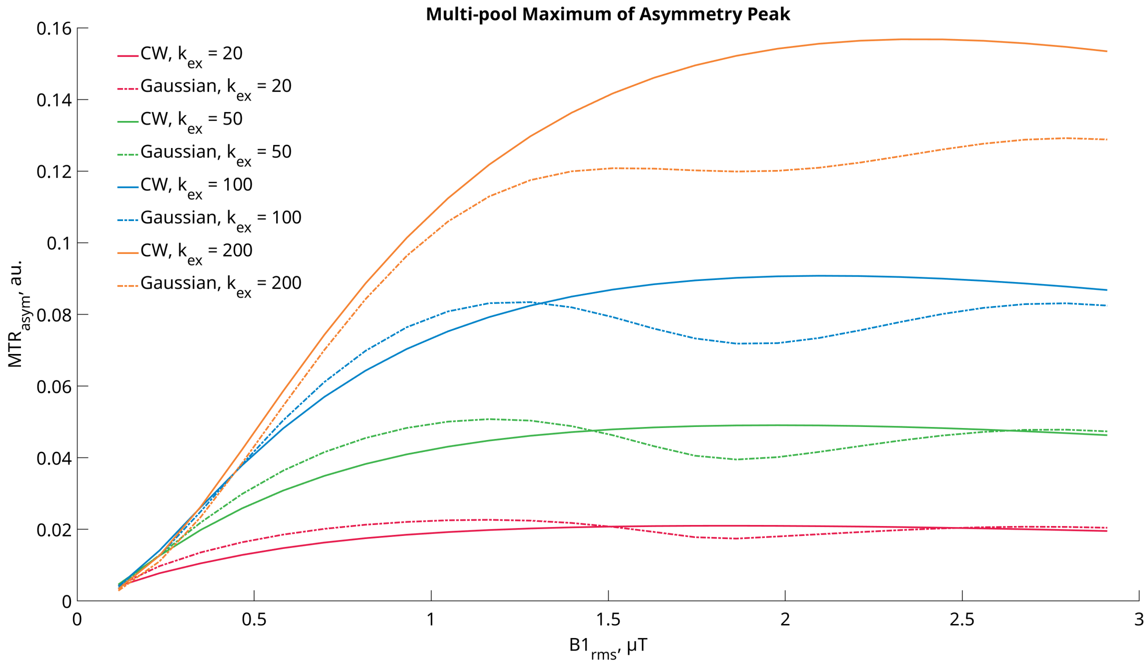


Figure S8: Multipool simulation for 7T. CW vs short Gaussian pulses. At different exchange rates and $B_{RMS}$. In this simulations, the $MTR_{asym}$ peak of short Gaussian pulses is higher for smaller $B_{1RMS}$ levels.

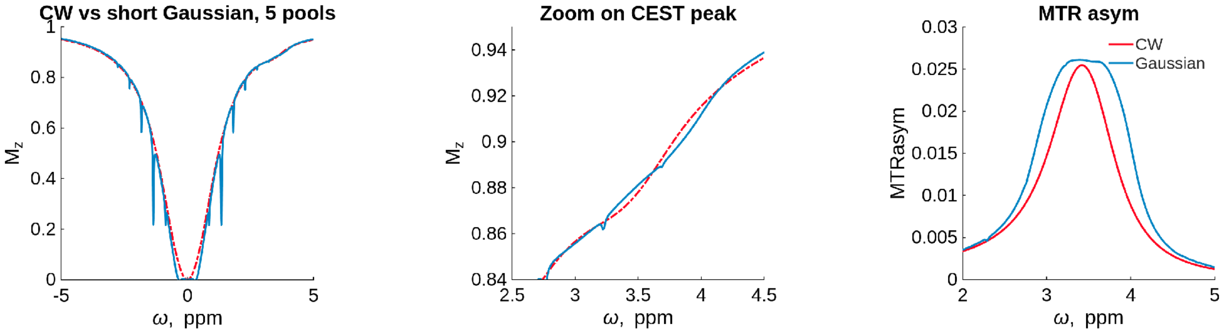


Figure S9: Multipool simulation for 3T. CW vs short Gaussian pulses. At exchange rate 25 Hz.

##### 5.1 OC for the multi pool case

For the multi pool case described in the previous section an OC pulse was designed. The target for the optimization were the spectrum of the short gaussian saturation. The optimization was not carried out for the whole spectrum but only between -4 and -2.5 and between 4 and 2.5 leaving out the water peak. the peak of the short gaussian pulses was not used as a target since we did not want to incorporate the sidebands in the optimization and we do not know what the perfect water spectrum in this case looks like.

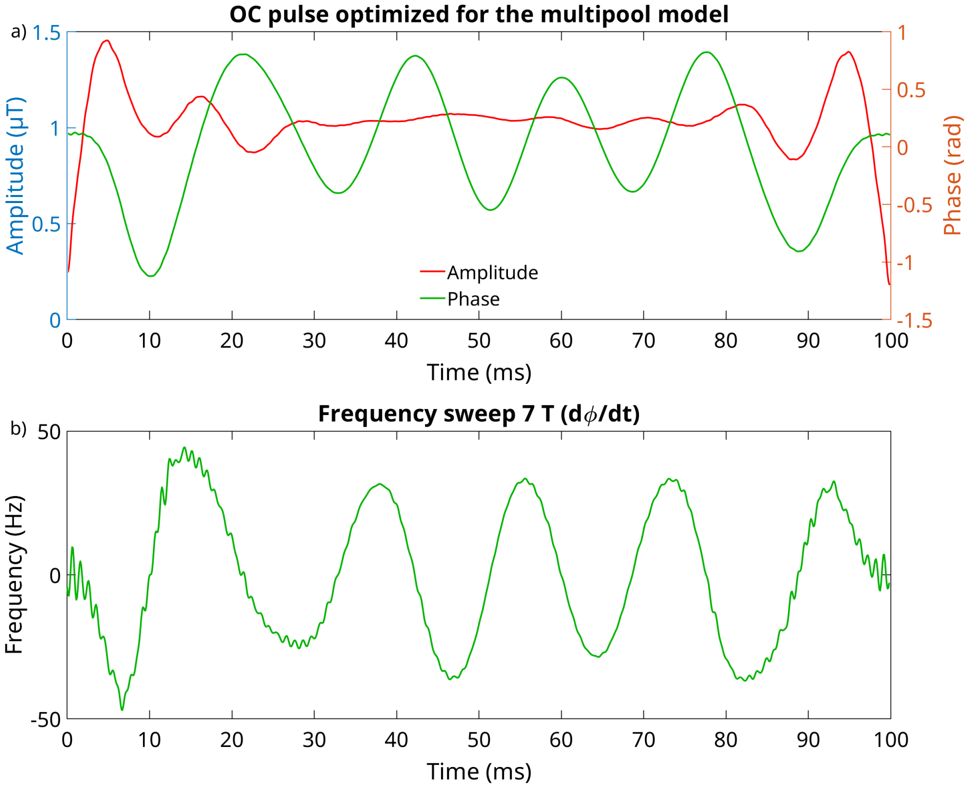


Figure S10: a) OC pulse designed for the multi pool case. b) Frequency sweep at 7 T.

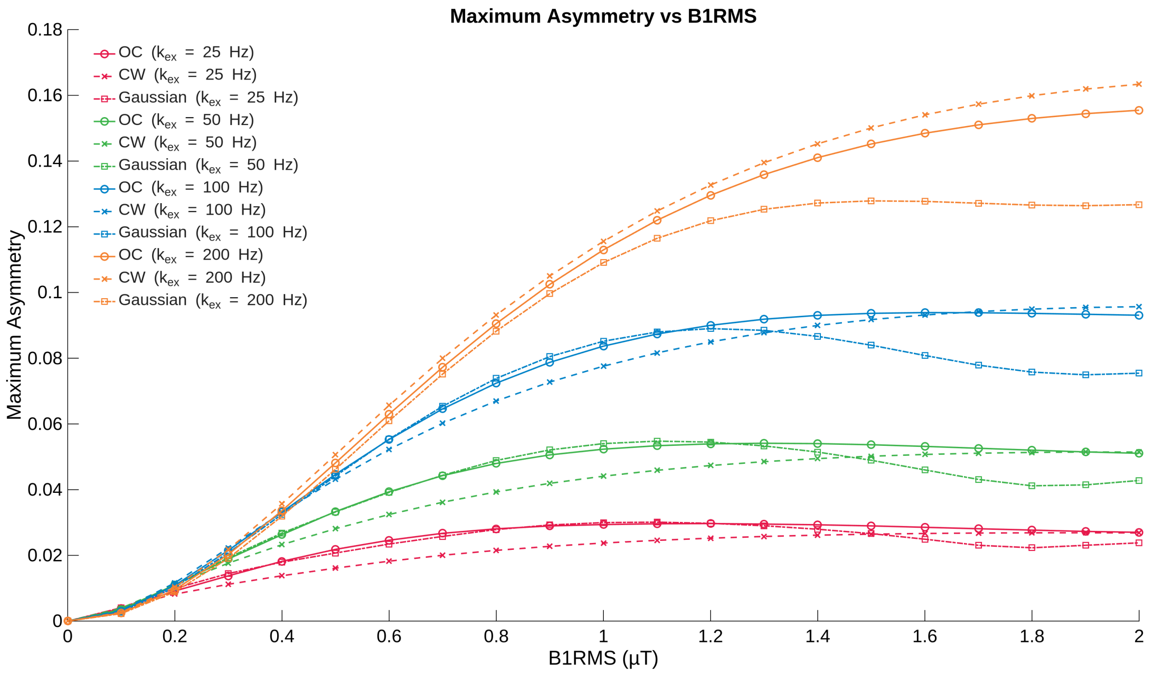


Figure S11: Multipool simulation for 7T.OC vs CW and short Gaussian pulses. At different exchange rates and $B_{RMS}$. In this case the OC saturation was able to exceed CW contrast.

#### 6 Frequency spectra of used pulses

The frequency spectra of a single 100 ms Gaussian, Fermi, Block and OC pulse are depicted in Figure S13. The frequency spectrum is generated by applying an FFT to the time amplitude signal. The bandwidth of the pulses is calculated using the FWHM. The Gaussian pulse has the highest bandwidth with 20 Hz, followed by the Fermi pulse with 17 Hz. The lowest FWHM values are found in the Block pulse and the OC pulse, with 12 and 13 Hz respectively. However, this analysis is only a rough estimate because it disregards the influence of the nonlinearity of the Bloch-McConnell equation on the z-spectrum.

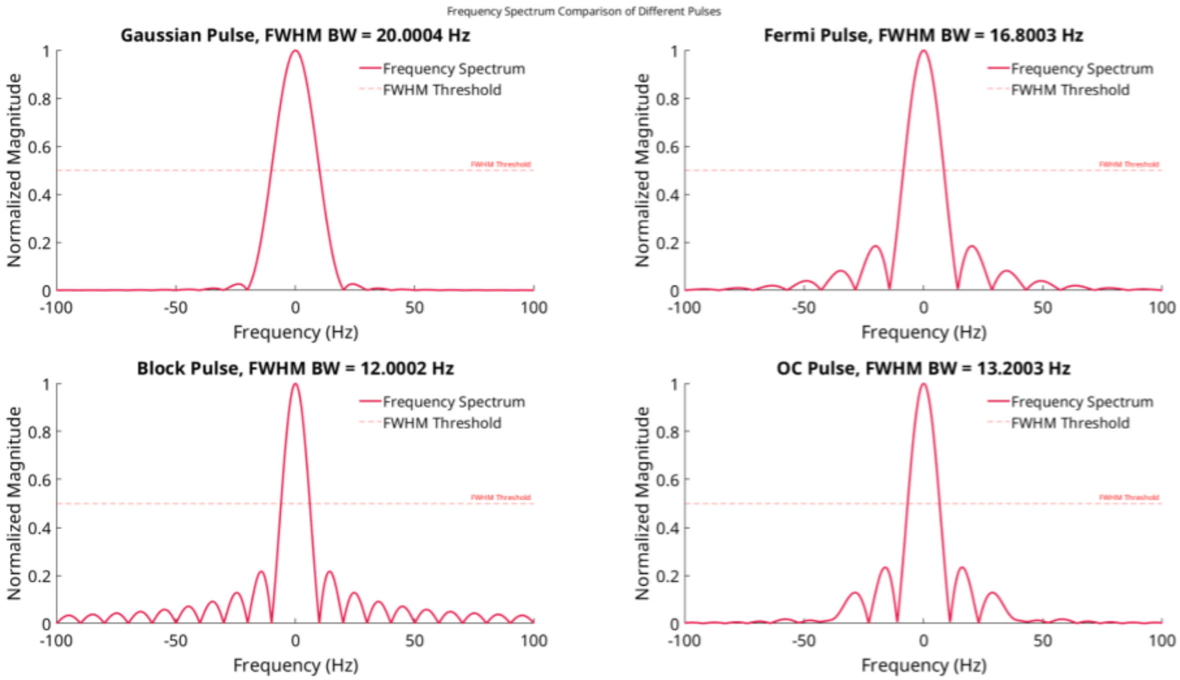


Figure S12: Frequency Spectra of a single 100 ms Gaussian, Fermi, Block and OC pulse

### 7 Phantom measurements

Example $T_{1w}$ image of the phantom used for phantom measurements. The phantom consisted of falcon tubes arranged by a 3D printed falcon tube holder in a water bath to make calibration of the scanner easier and to avoid big susceptibility jumps through air. The water outside the falcon tubes was without contrast agent.

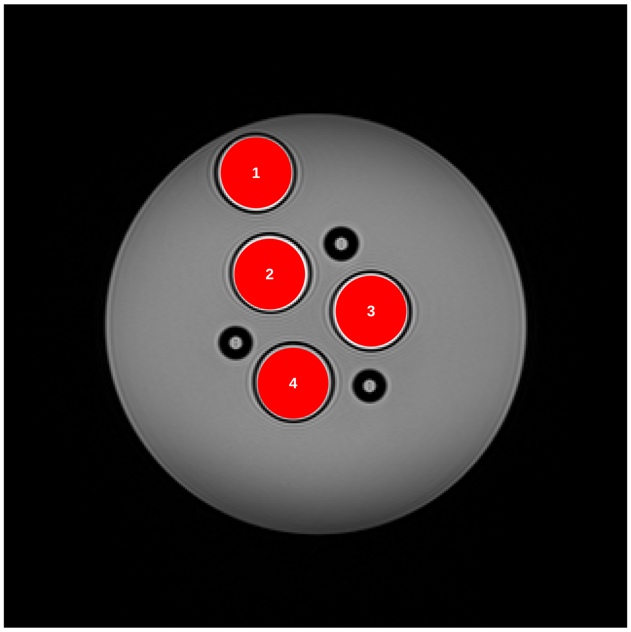


Figure S13: $T_{1w}$ weighted image of the CEST phantom, coronal slice with ROIs.

### 8 Details to the optimization

The optimization process employs a set of stopping criteria to ensure efficient and accurate convergence. The algorithm terminates when one of the following conditions is met:

1. the relative gradient norm falls below a threshold of $10^{-6}$ times the initial gradient norm, indicating sufficient convergence
2. the absolute gradient norm becomes smaller than $10^{-8}$
3. the maximum number of iterations, set to 1000, is reached to prevent excessive computation
4. the trust region radius shrinks below $10^{-12}$, signaling optimization failure
5. no accepted solution is found within 100 consecutive iterations, indicating stagnation.

For testing the global convergence, the optimization was initialized with a completely random starting pulse. We generate $B_1(t)$ by drawing 1000 amplitude points from a uniform random distribution and scaling them to achieve RMS power $P$:

$$B_1(t) = \frac{u(t)}{\text{RMS}(u)} \cdot P \quad \text{where } u(t) \sim \text{Uniform}(0, 1) \text{ for each } t$$

where $\text{RMS}(u) = \sqrt{\overline{u^2}}$ is the root-mean-square over all time points. This provides a stochastic initialization while maintaining the power constraint. The optimization results from multiple runs indicate robustness to local and global minima. The maximum difference in pulse amplitude is smaller than 1 nT, and the maximum difference in the simulated spectra is smaller than $7 \times 10^{-6}$ % of the maximum water signal.

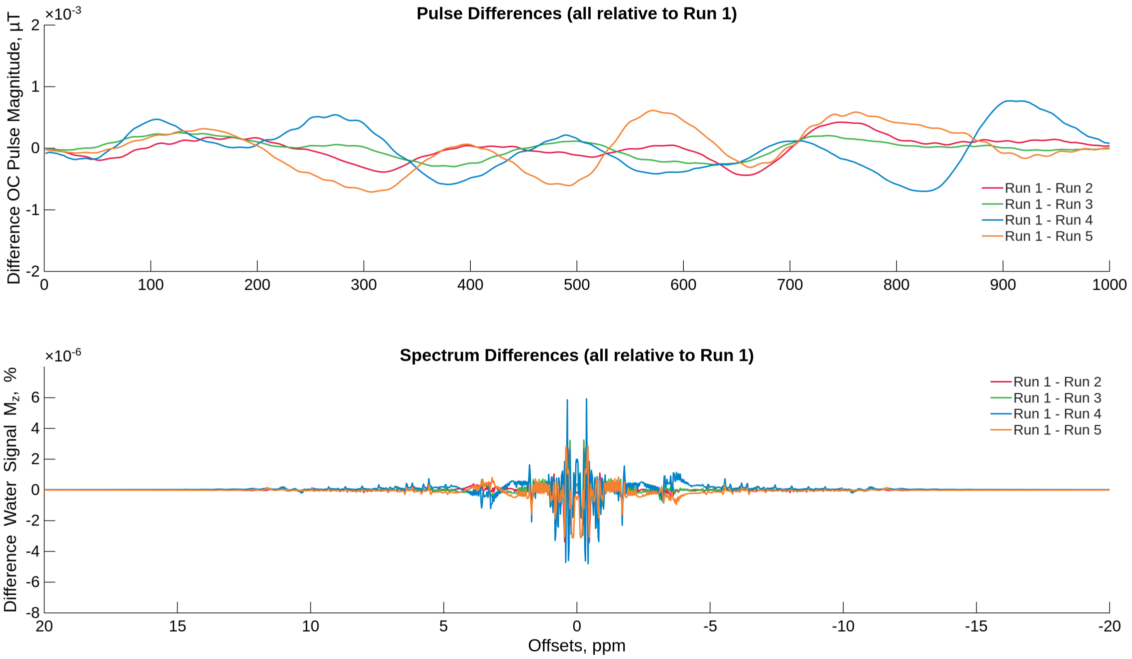


Figure S14: OC optimization results from 5 runs with random intiialization. Difference in pulse shape (top) and difference in simulated spectra (bottom).

#### 9 In vivo thigh $B_0$ map

WASABI $B_0$ map for the thigh measurement in Figure S15.

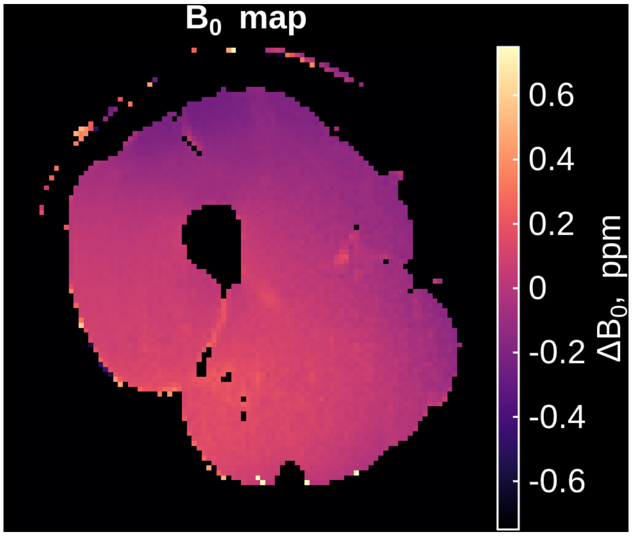


Figure S15: WASABI $B_0$ map for the thigh measurement.

#### 10 Performance of the different pulses in simulation

Pulseq CEST simulation of all 100 ms pulses used in the paper over different $B1$ scalings (Figure S16). The simulated values were:

Pulse train parameters: 8 pulses, $t_p = 100$ ms, DC = 90 %.

BM pool model: water pool ($f = 1.0$, $T_1 = 1.2$ s, $T_2 = 0.080$ s) and four exchangeable pools: creatine ($f = 0.0035$, $T_1 = 1.2$ s, $T_2 = 0.160$ s, $k = 250$ Hz, $\Delta\omega = 1.7$ ppm), IOP ($f = 0.0021$, $T_1 = 1.2$ s, $T_2 = 0.160$ s, $k = 1000$ Hz, $\Delta\omega = 4.2$ ppm), NA ($f = 0.0017$, $T_1 = 1.2$ s, $T_2 = 0.160$ s, $k = 250$ Hz, $\Delta\omega = 3.2$ ppm), and OH ($f = 0.002$, $T_1 = 1.2$ s, $T_2 = 0.160$ s, $k = 1000$ Hz, $\Delta\omega = 1.2$ ppm).

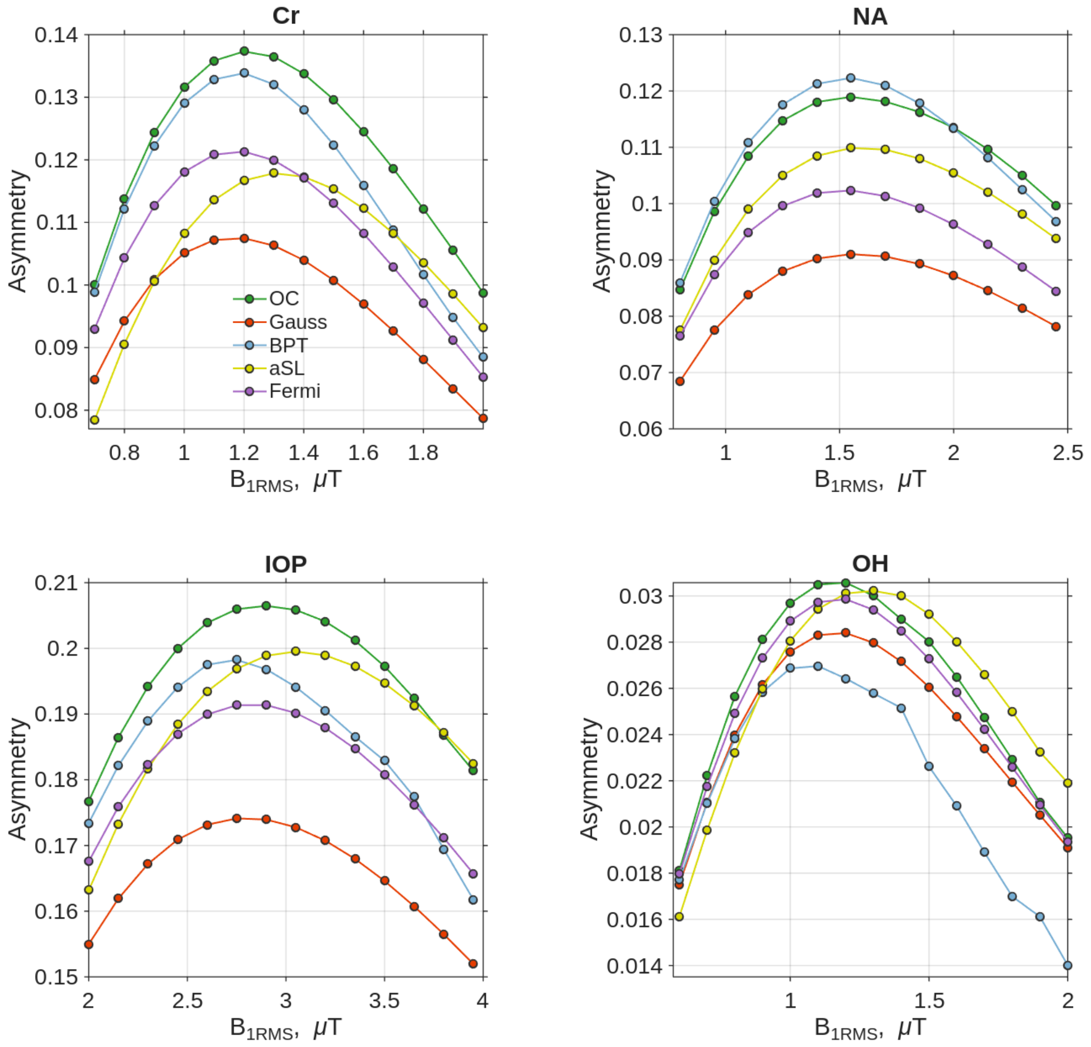


Figure S16: Pulseq CEST simulation: $MTR_{asym}$ for all pulse shapes used in manuscript, simulated for parameters expected in the phantom measurements. $B1$ level was adjusted to show the maximum point for every pulse

### 11 Performance comparison of temporally stretched and compressed OC pulses

To minimize artifacts in the OC spectra shown in Figure 3 e,f,g,h of the paper, the 100 ms OC pulse was used for scalings between 100 and 76 ms. For scalings between 75 and 50 ms the 50 ms pulse was used. The correct pulses and interpolations are implemented in the `makeOCPulse` in pulseq CEST and is automatically selected based on the user input of `td`.

Pulseq CEST simulation for the creatine parameters in section 10 for different pulse times in 1 s pulse train with DC of approximately 90 %.

For different pulse times the OC shows constant highest saturation (Figure S18). The 100 ms pulse generates slightly higher contrast than the 50 ms, independent of the scaling in time.

High resolution (0.01 ppm) spectra for all pulse shapes and different pulse times can be seen in Figure S19. These spectra are simulated with a $B_0$ inhomogeneity of 0.1 ppm.

The pulse train parameters for both Figures S18, S19 were chosen to have a constant DC of 91 % a $T_{sat}$ of approximately 1 s and a $B_{1rms}$ of 1 µT.

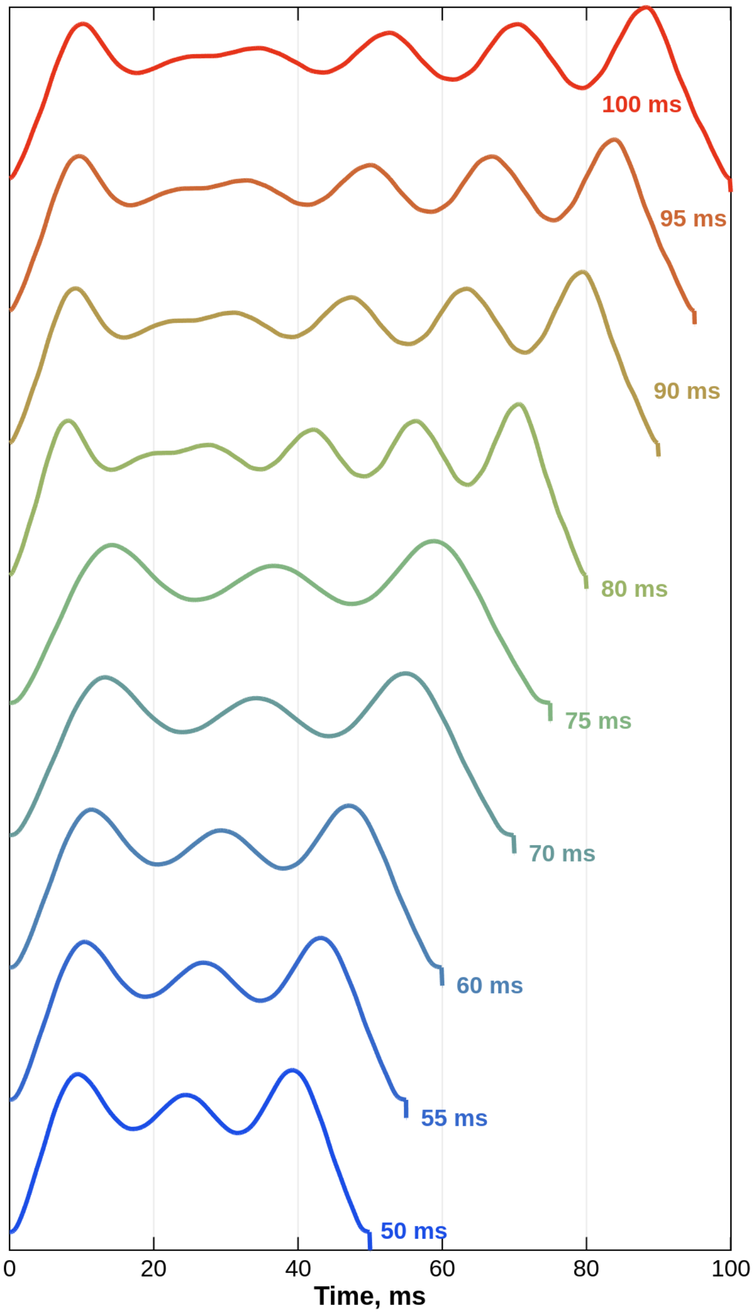


Figure S17: Pulse shapes stretched and compressed to different times $t_p$.

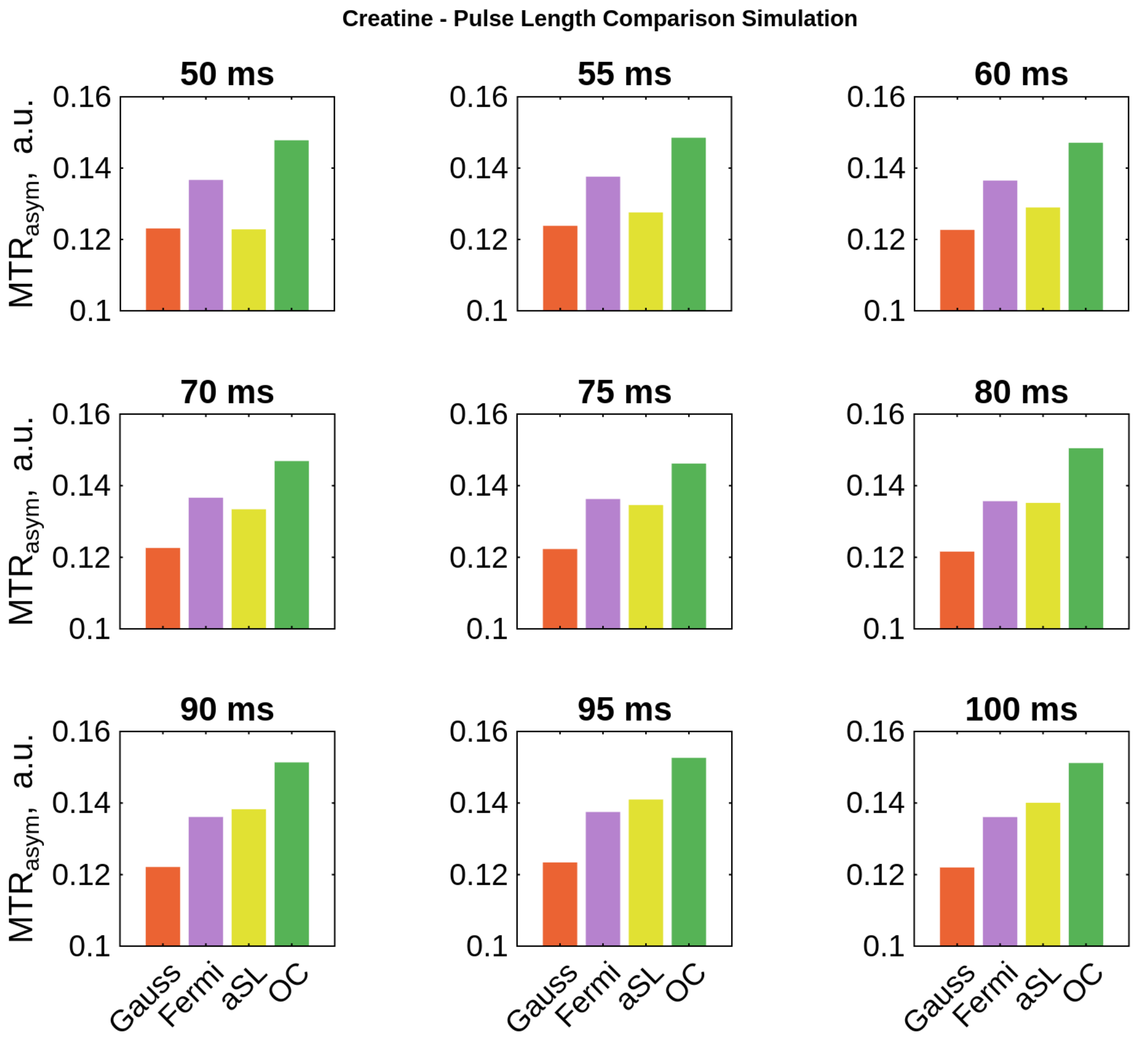


Figure S18: $MTR_{asym}$ for different pulse times and Gauss, Fermi, aSL and OC pulse simulated for the creatine pahntom.

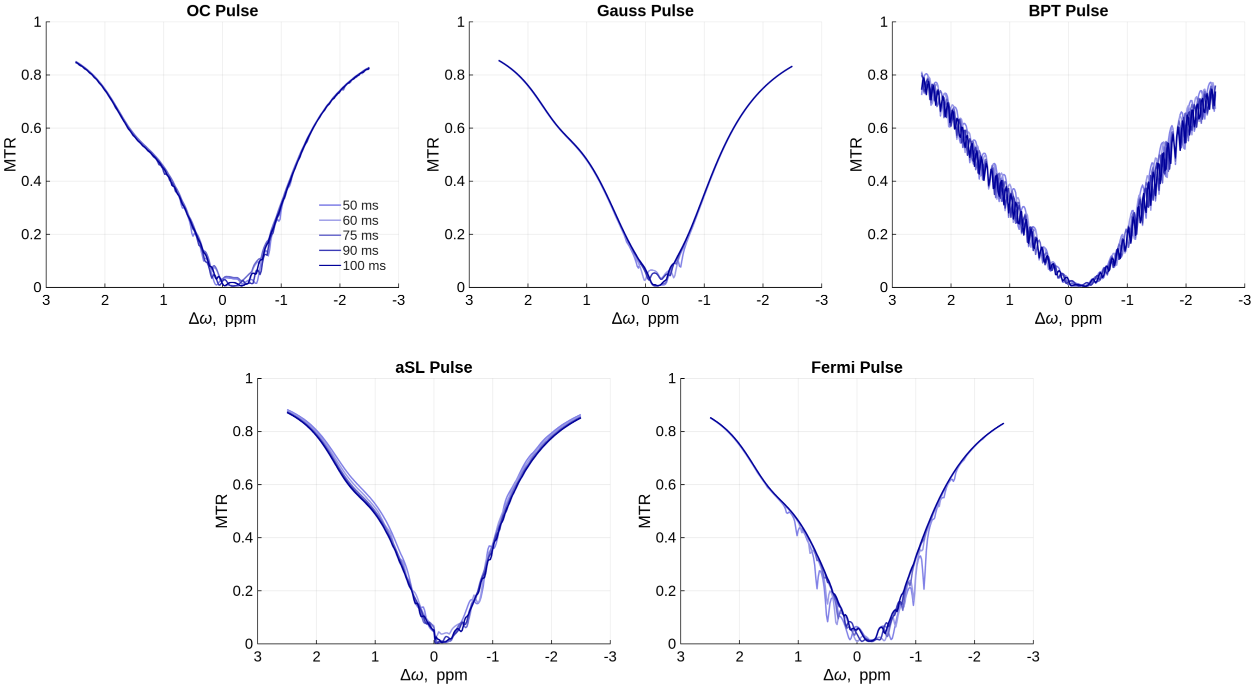


Figure S19: Spectra simulated with different pulse lengths and a high frequency resolution of 0.01 ppm and a $B_0$ inhomogeneity of 0.1 ppm.

#### 12 Lorentzian fitting

Parameter maps for the APT brain Lorentzian fitting can be seen in Figure S21. Water and MT were fitted with a 2 pool Lorentzian model. For the fit a pixel wise Lorentzian model was fitted with a least squares algorithm. Both saturation lead to approximately the same MT maps. The on resonant saturation of the OC was higher than the Gaussian saturation. This is in agreement with simulations. The OC APT contrast generated is approximately 30 % higher than with Gaussian saturation.

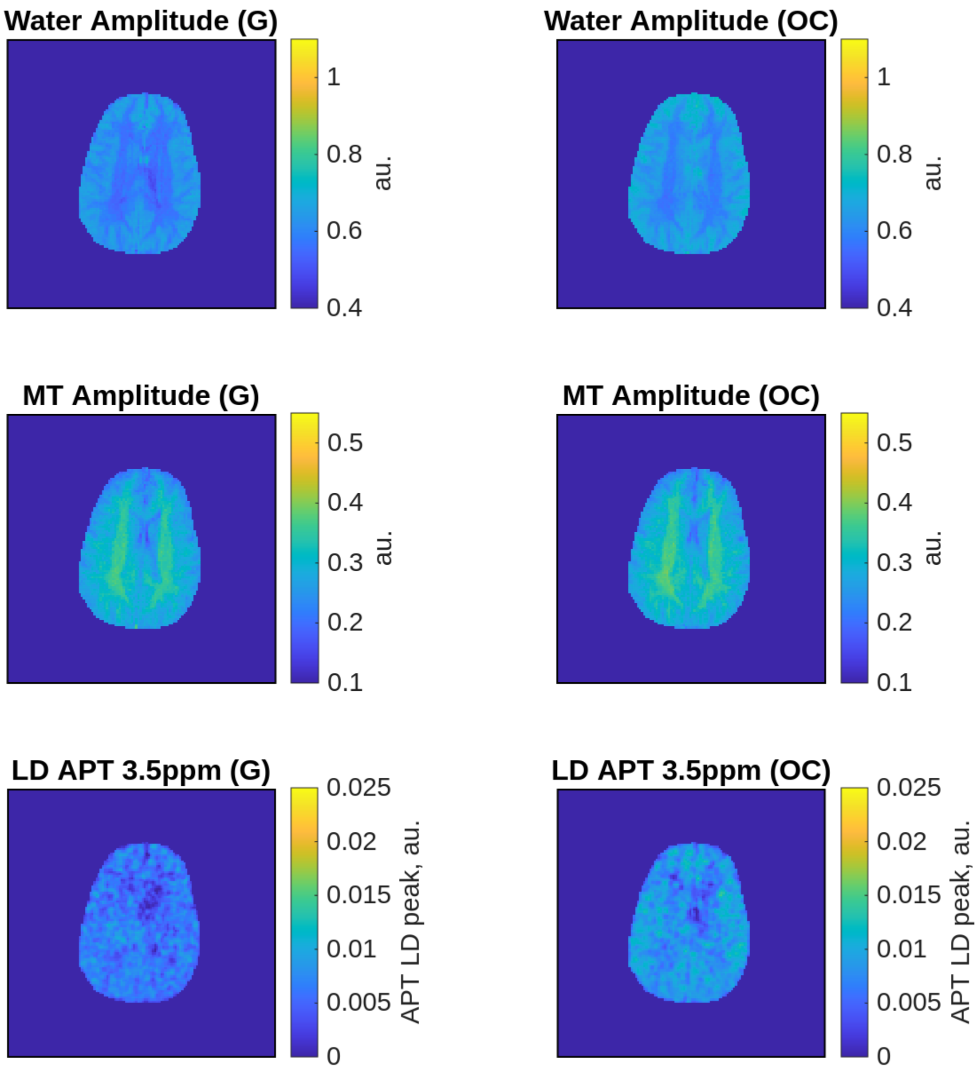


Figure S20: Parameter maps from the Lorentzian fitting. And the Lorentzian difference maps resembling the extracted APT contrast.

### 13 ROI for the SNR calculation

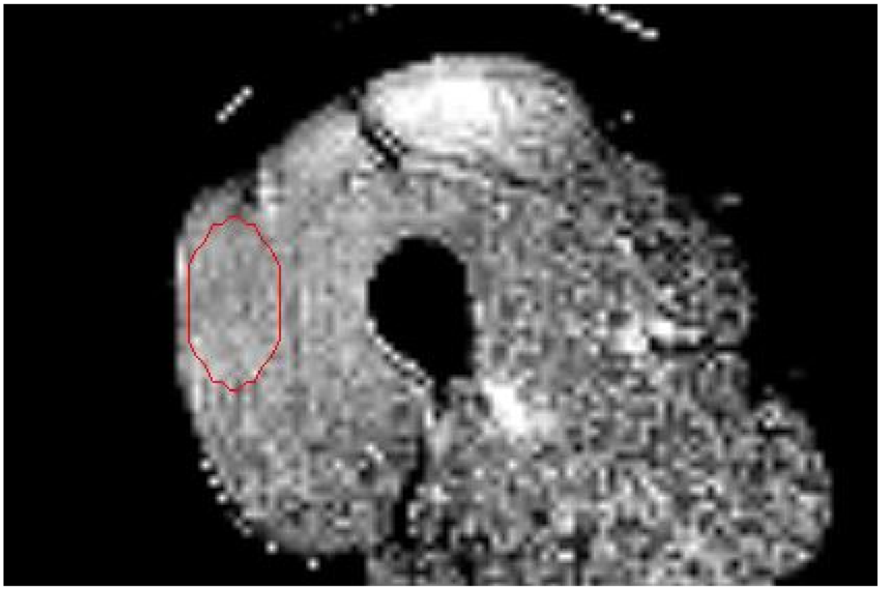

Figure S21: ROI in the PCA-denoised and $B_0$-corrected OC $MTR_{asym}$ image used for the calculation of the SNR.
